# Interaction Between Transcribing RNA Polymerase and Topoisomerase I Prevents R-loop Formation in *E. coli*

**DOI:** 10.1101/2021.10.26.465782

**Authors:** Dmitry Sutormin, Alina Galivondzhyan, Olga Musharova, Dmitrii Travin, Anastasiya Rusanova, Kseniya Obraztsova, Sergei Borukhov, Konstantin Severinov

## Abstract

Bacterial topoisomerase I (TopoI) removes excessive negative supercoiling and is thought to relax DNA molecules during transcription, replication and other processes. Using ChIP-Seq, we show that TopoI of *Escherichia coli* (EcTopoI) is co-localized, genome-wide, with RNA polymerase (RNAP) in transcription units. Treatment with transcription elongation inhibitor rifampicin leads to EcTopoI relocation to promoter regions, where RNAP also accumulates. When a 14 kDa RNAP-binding EcTopoI C-terminal domain (CTD) is overexpressed, co-localization of EcTopoI and RNAP along the transcription units is reduced. Pull-down experiments directly show that the two enzymes interact *in vivo*. Using ChIP-Seq and Topo-Seq, we demonstrate that EcTopoI is enriched and in and upstream (within up to 12-15 Kbs) of highly-active transcription units, indicating that EcTopoI relaxes negative supercoiling generated by transcription. Uncoupling of the RNAP-EcTopoI interaction by either overexpression of EcTopoI CTD or deletion of EcTopoI domains involved in the interaction is toxic for cells and leads to excessive negative plasmid supercoiling. Moreover, the CTD overexpression leads to R-loops accumulation genome-wide, indicating that the RNAP-EcTopoI interaction is required for prevention of R-loops formation.

**Article Highlights:**

- TopoI colocalizes genome-wide and interacts with RNAP in *E. coli*
- Disruption of the interaction between TopoI and RNAP decreases cells viability, leads to hypernegative DNA supercoiling, and R-loops accumulation
- TopoI and DNA gyrase are enriched, respectively, upstream and downstream of transcription units in accordance with twin-domain model of Liu and Wang
- TopoI recognizes its cleavage sites through a specific motif and by sensing negative supercoiling

## Introduction

An optimal level of DNA supercoiling is required for DNA replication and transcription (Postow *et al*., 2001; Dorman, 2019), DNA compaction, effective bulk segregation of chromosomes, and site-specific DNA recombination and repair (Saha *et al*., 2013; Wang, Montero Llopis and Rudner, 2013). Topoisomerases, a conserved and ubiquitous group of enzymes, contribute to and regulate the extent of DNA supercoiling and its other topological properties (Maxwell, Bush and Evans-Roberts, 2015). Topoisomerases are divided into two types: type I enzymes introduce a transient single-strand break into DNA while type II enzymes introduce a transient double-strand break (Maxwell, Bush and Evans-Roberts, 2015).

Topoisomerase I of *Escherichia coli* (EcTopoI, encoded by the *topA* gene) belongs to the A class of type I topoisomerases (Maxwell, Bush and Evans-Roberts, 2015). EcTopoI relaxes only negatively supercoiled DNA and is thought to maintain the steady-state level of supercoiling by compensating the activity of another topoisomerase – the DNA gyrase, a IIA type enzyme, which introduces negative supercoiling utilizing the energy of ATP hydrolysis (Menzel and Gellert, 1978; Tse-Dinha, 1985; Liu *et al*., 2017). Deletion of *topA* leads to rapid accumulation of suppressor mutations, mostly in genes encoding the DNA gyrase subunits. By reducing gyrase activity these mutations balance the level of DNA supercoiling inside the cell (Dinardo *et al*., 1982; Pruss *et al*., 1982). Amplification of a chromosome region containing the *parC* and *parE* genes encoding topoisomerase IV (TopoIV) is also frequently reported in *topA* null mutants (Dorman *et al*., 1989; Kato *et al*., 1990; Brochu *et al*., 2018). Conversely, *topA* deletions complement growth and replication defects of temperature-sensitive (Ts) *gyrB* mutants at non-permissive temperatures (Usongo *et al*., 2013).

Hypernegative supercoiling is a hallmark of *topA* mutants resulting from uncompensated gyrase activity (Baaklini *et al*., 2008). It was proposed that hypernegative supercoiling leads to stabilization of R-loops – RNA-DNA heteroduplexes forming when nascent RNA transcripts anneal to the template DNA strand upstream of the transcribing RNA polymerase (RNAP) (Masse and Drolet, 1999; Baaklini *et al*., 2008; Stolz *et al*., 2019). Indeed, R-loops have been recently detected in *topA* mutants by performing dot-blots with the RNA:DNA hybrid specific antibody S9.6 (Brochu, Vlachos-Breton and Drolet, 2021). Since in the R-loop one DNA strand is unpaired, a hub that accumulates excessive negative supercoiling is created, leaving the nearby DNA less negatively supercoiled. Since such DNA is a substrate for DNA gyrase, which introduces more negative supercoiling, a positive feedback loop is created, leading to further accumulation of R-loops and more negative supercoiling (negative supercoiling R-loops masking of negative supercoiling increase gyrase activity excessive negative supercoiling) (**Figure 1**) (Drolet, 2006). Nascent transcripts in R-loops are degraded, leading to rapid growth arrest (Baaklini *et al*., 2008).

**Figure 1.**
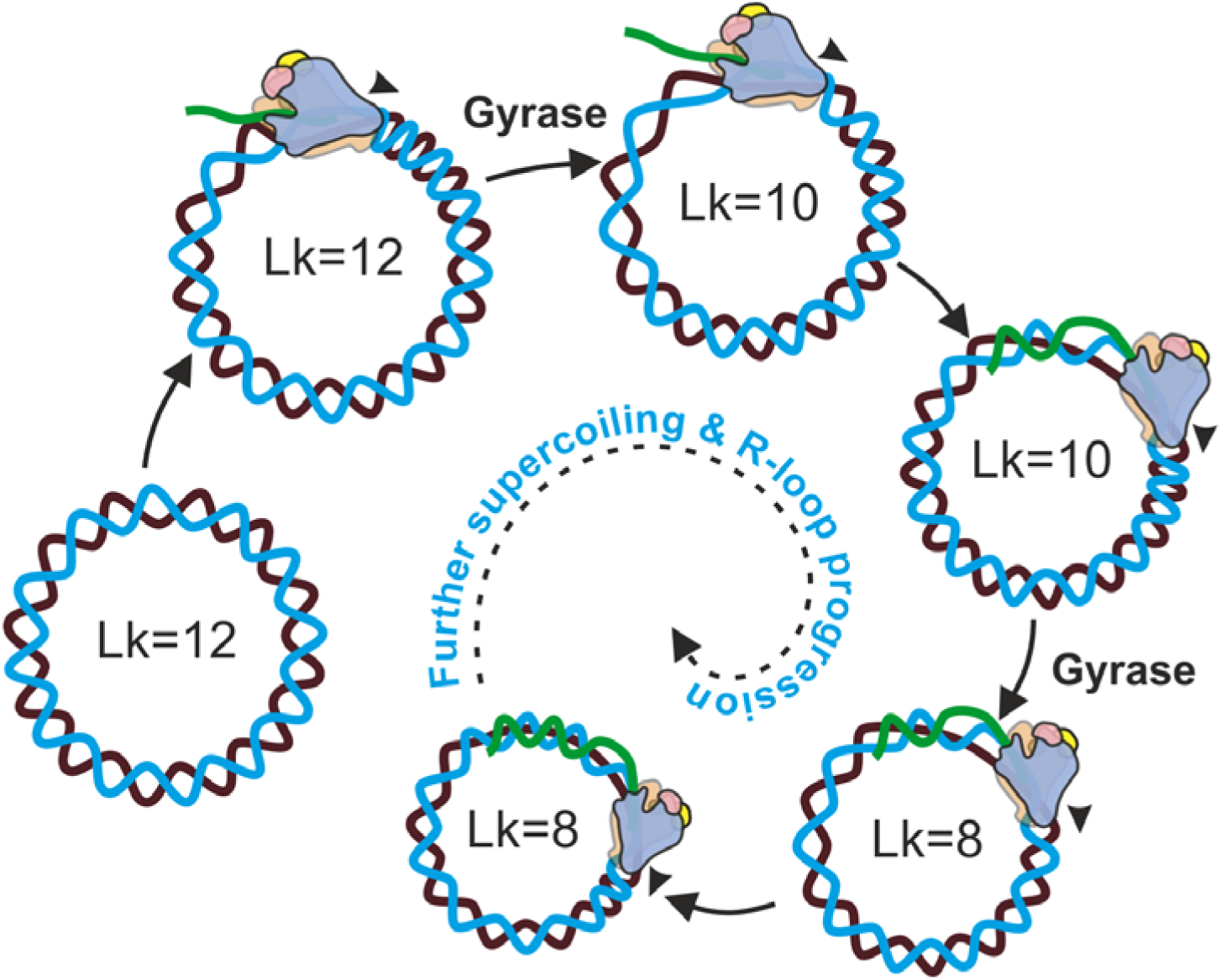
A positive feedback cycle that leads to hypernegative supercoiling of plasmid DNA and R-loops accumulation when transcription proceeds in the presence of DNA gyrase and in the absence of TopoI.

Overexpression of RNAse HI, an enzyme which degrades RNA in the R-loops (Kogoma, 1997; Drolet and Brochu, 2019) and should thus break the feedback loop, was reported to partially suppress the negative effects of *topA* deletion (Drolet *et al*., 1995; Hraiky, Raymond and Drolet, 2000), although this finding was disputed (Stockum, Lloyd and Rudolph, 2012). Conversely, deletion of the RNAse HI *rnhA* gene exacerbates the *topA* null phenotype (Usongo *et al*., 2008; Stockum, Lloyd and Rudolph, 2012). Stabilized R-loops can also prime *oriC*-independent replication – a phenomenon called “constitutive stable DNA replication” (cSDR) initially observed in cells lacking RNAse HI. It was demonstrated that cells lacking type-I topoisomerases also exhibit cSDR, which is suppressed by overexpression of RNAse HI (Martel *et al*., 2015; Brochu *et al*., 2018). Together, these data indicate that hypernegative supercoiling is the likely cause of severe growth defects of non-suppressed *topA* mutants (Baaklini *et al*., 2004). The EcTopoI was shown to bind RNAP, and the interaction was mapped to the C-terminal portion of EcTopoI and the β’ subunit of RNAP (Cheng *et al*., 2003). The TopoI-RNAP interaction was also reported for mycobacteria (Banda, Cao and Tse-Dinha, 2017) and *Streptococcus pneumoniae* (Ferrandiz, Hernandez and de la Campa, 2021). It was hypothesized that association with RNAP enables TopoI to rapidly relax negative DNA supercoils forming behind the elongating RNAP, thereby preventing the R-loops formation (Cheng *et al*., 2003; Yang *et al*., 2015). The chromosomal distribution of EcTopoI is currently unknown, although some sequence preference *in vitro* has been reported (Y. Tse, Kirkegaard and Wang, 1980; Kirkegaard, Pflugfelder and Wang, 1983). Recently, the genome-wide distribution of TopoI from *M. tuberculosis* (MtTopoI), *M. smegmatis* (MsTopoI), and *S. pneumoniae* (SpTopoI) was investigated using ChIP-Seq (Ahmed *et al*., 2017; Rani and Nagaraja, 2018; Ferrandiz, Hernandez and de la Campa, 2021). In all cases, the topoisomerase was shown to associate with actively transcribed genes with particular enrichment upstream of RNAP peaks at promoter regions. These findings agree with the twin-domain model proposed by Liu and Wang (Liu and Wang, 1987; Wu *et al*., 1988), but do not necessarily imply direct TopoI-RNAP association.

In the present work we map the EcTopoI binding sites on the *E. coli* chromosome using ChIP-Seq. We demonstrate that during exponential growth the enzyme is accumulated in regions with high levels of transcription, where it colocalizes with RNAP at promoters and transcription unit (TU) bodies. EcTopoI is also significantly enriched in extended, 12-15 Kbs, regions upstream of transcribed TUs. When transcription is inhibited by rifampicin (Rif), both EcTopoI and RNAP redistribute from TUs bodies toward promoter regions and EcTopoI becomes absent in the upstream regions. When a 14 kDa C-terminal domain of EcTopoI known to interact with RNAP (Cheng *et al*., 2003) is overexpressed, EcTopoI enrichment is decreased in TU bodies and promoter regions, but unaffected at the upstream regions. By mapping of cleavage sites induced by an “intrinsically-poisoned” EcTopoI mutant we reveal that EcTopoI catalytic activity is increased in upstream region of highly active TUs. Based on these data and pull-down experiments we conclude that EcTopoI physically interacts with RNAP in TUs in a CTD-dependent manner. At the same time, and independently of RNAP, EcTopoI is attracted to extended upstream regions in front of TUs by negative supercoiling generated by transcription and removes supercoils. We finally demonstrate that long-term overexpression of the 14 kDa CTD is lethal for cells and results in hypernegative supercoiling and accumulation of R-loops. We propose that the interaction between RNAP and EcTopoI is essential for DNA duplex restoration immediately upstream of the elongating RNAP, either by *in situ* relaxation of negative superhelicity or by structural clamping of DNA by the topoisomerase.

## Materials & Methods

### Strains and plasmids

*E. coli* DY330 *topA-SPA* (W3110 *lacU169 gal490 cI857* Δ (*cro-bioA*) *topA*-SPA) strain with *topA* gene fused with the sequence encoding the SPA tag (purchased from Dharmacon) was used for EcTopoI ChIP-Seq and Topo-Seq experiments, supercoiling experiments, and toxicity experiments. *E. coli* DY330 (W3110 *lacU169 gal490 cI857* Δ (*cro-bioA*)) was used for amplification of *topA* gene. *E. coli* DH5α strain was used for standard cloning. *E. coli* BW25113 was used for *topA* gene editing. *E. coli* DY330 *rpoC-TAP* (W3110 *lacU169 gal490 cI857* Δ (*cro-bioA*) *rpoC*-TAP) strain with *rpoC* gene fused with the sequence encoding the TAP tag (purchased from Dharmacon) was used for RNAP ChIP-Seq experiments. pCA24 plasmid was used for cloning and overexpression of EcTopoI 14kDa CTD, EcTopoI, or GFP. pCA24 topA plasmid was obtained from *E. coli* ASKA collection (Kitagawa *et al*., 2005). pET28 plasmid was used for overproduction and purification of EcTopoI in *E. coli* BL21(DE3) strain.

### Cloning of EcTopoI and EcTopoI CTD

Full length *topA* gene was amplified from genomic DNA extracted from *E. coli DY330* (Genomic DNA extraction kit, ThermoFisher Scientific) as two overlapping fragments to remove NcoI restriction site at the end of the gene with following primers: topA_NcoI_fw + NcoI_mut_rev, NcoI_mut_fw + topA_HindIII_strepII_rev. Primer sequences can be found in **Table S1**. Full length *topA* gene was reconstituted from fragments with overlap extension PCR using topA_NcoI_fw and topA_HindIII_strepII_rev primers. Resulting PCR product was cloned into pET28 plasmid by NcoI and HindIII restriction sites giving pET28 topA_strep plasmid. C-terminal StrepII tag was introduced with topA_HindIII_strepII_rev primer.

DNA fragment encoding EcTopoI 14kDa CTD was amplified from genomic DNA of *E. coli* DY330 with TopoA_14_kDa_CTD_BamHI_fw and TopoA_CTD_HindIII_rev primers (**Table S1**) and cloned by BamHI and HindIII sites into pCA24 plasmid giving pCA24 14kDa CTD construct. To prevent T5-lac promoter leaking, strains transformed with the plasmid were grown on media containing 0.5% glucose for catabolite repression. Obtained plasmids were verified by Sanger sequencing.

### Construction of pBAD33 topA_strepII G116S M320V plasmid

To incorporate substitutions *topA* gene fused with strepII-coding sequence was PCR-amplified from pET28 topA_strep plasmid in three overlapping fragments using the following primer pairs: topA_XbaI_RBS_fw+topA_G116S_out_rev, topA_G116S_in_fw+topA_M320V_in_rev, and topA_M320V_out_fw+topA_strepII_HindIII_rev. The fragments were fused using overlap extension PCR with primers topA_XbaI_RBS_fw+topA_strepII_HindIII_rev (**Table S1**). The final amplicon preliminarily treated with DpnI was cloned into pBAD33 by XbaI and HindIII restriction sites resulting in a pBAD33 topA_strepII G116S M320V plasmid. To prevent promoter leaking, strains transformed with the plasmid were grown on media containing 0.5% glucose for catabolite repression. Obtained plasmid was verified by Sanger sequencing.

### Genome editing, editing of topA gene

Editing of *topA* gene in *E. coli* BW25113 cells was performed by recombineering approach using Lambda Red system on pKD46 plasmid (Datsenko and Wanner, 2000). Recombination cassettes were obtained by PCR from pKD4 plasmid with primers having flanks homologous to the sites of desired recombination: topA_delta_topA66_kanR_F+topA_SPA_kanR_cysB_R, topA_delta_14kDa_kanR_F+topA_SPA_kanR_cysB_R, topA_delta_30kDa_kanR_F+topA_SPA_kanR_cysB_R (**Table S1**). Three versions of *topA* truncations from the 5’-end were obtained: *topA*Δ*11kDa*, *topA*Δ*14kDa*, and *topA*Δ*30kDa*.

### Whole genome sequencing, identification of mutations

DNA was extracted from 3 mL of *E. coli* BW25113 or *E. coli* BW25113 *topA* mutants night culture using GeneJET Genomic DNA purification kit (ThermoFisher) according to the manufacturer protocol. NGS libraries were made using NEBNext Ultra II DNA Library Prep kit (NEB). DNA sequencing was performed on Illumina MiniSeq with 150+150 bp paired-end protocol.

Raw reads were filtered and trimmed with Trimmomatic (Bolger, Lohse and Usadel, 2014) and then were aligned to the *E. coli* W3110 MuSGS genome (*E. coli* W3110 genome with the insertion of *cat*-Mu SGS cassette may be downloaded from GEO: GSE95567) using BWA-MEM (Li and Durbin, 2010). BAM and bed files were prepared with Samtools (Li *et al*., 2009) and visualized in IGV (Thorvaldsdóttir, Robinson and Mesirov, 2013). SNPs and short indels were identified with bcftools mpileup followed by bcftools call programs. Large scale chromosomal deletions and region multiplications were inspected by manual analysis of genome coverage depth.

### Toxicity of EcTopoI CTD

Long term toxicity of the topoisomerase I CTD was assessed by CFU counting of *E. coli* DY330 transformed with pCA24 14kDa CTD plasmid. *E. coli* DY330 without plasmids and *E. coli* DY330 transformed with pCA24 GFP served as controls. Overnight cultures were grown at 37°C in LB or LB supplemented with 0.5% glucose or LB supplemented with 1 mM IPTG. Chloramphenicol (34 µg/mL) was added to cell cultures harboring a plasmid. Serial dilutions of the cultures were applied on LB plates supplemented with corresponding additions (0.5% glucose or 1 mM of IPTG) or without additions. CFU were counted after overnight incubation at 37°C.

To assess short-term toxicity of the topoisomerase I CTD a culture of *E. coli* DY330 transformed with pCA24 14kDa CTD was grown in LB supplemented with chloramphenicol (34 µg/mL) and glucose (0.5%) until reaching OD_600_=0.2. Then the culture was bisected and one half was induced with 1mM IPTG. After 1 h of additional culturing CFU were counted on LB plates without the inducer by serial dilution method.

### Analysis of plasmids supercoiling

Topoisomers were separated in 1% TAE agarose gel containing chloroquine (concentrations range from 0.5 to 7.5 µg/ml, indicated for each experiment separately) with subsequent washing in TAE and staining with ethidium bromide.

### EcTopoI ChIP-Seq

For EcTopoI ChIP-Seq we used *E. coli* DY330 strain (Dharmacon) encoding EcTopoI fused with SPA-tag (Butland *et al*., 2005). Cell culture (V=40 mL) grown to mid-exponential phase OD_600_∼0.5-0.7 in LB containing 50 µg/mL of kanamycin was crosslinked by adding fresh formaldehyde to the final concentration of 1%, followed by incubation for 20 min at room temperature, with agitation. Crosslinking was stopped by the addition of sterile glycine to a final concentration of 0.125 M. Cells were incubated for 10 min at room temperature and harvested by centrifugation at 4500 g, 5 min, 4°C. Cell pellets were washed three times with 10 mL of ice-cold PBS and finally resuspended in 1 mL of FA lysis buffer (50 mM HEPES-NaOH pH 7.5, 1 mM EDTA, 0.1% deoxycholate, 0.1 % SDS, 1% Triton X-100, 150 mM NaCl), followed by 10 min incubation on ice. Then, protease inhibitors cocktail (1x concentration, cOmplete ULTRA, Sigma-Aldrich) and RNase A (0.1 mg/ml, Thermo Scientific) were added. To disrupt cells and shear DNA, cells were sonicated in a 1.5 mL Eppendorf tube in ice-water bath to achieve 100-1000 bp fragments range. Lysates were clarified by centrifugation at 10000 g for 5 min at 4°C, and the resulting supernatant was used for further analysis.

For Input DNA 100 µL of lysate was treated and de-crosslinked by incubation with proteinase K (Thermo-Fisher Scientific) for 4 h at 55°C and DNA was purified using DNA clean-up kit (Thermo Fisher Scientific). DNA fragmentation range was assessed by electrophoresis.

For EcTopoI immunoprecipitation, 900 µL of the remaining lysate were diluted with 1 mL of TES buffer (10 mM Tris-HCl pH 7.5, 1 mM EDTA, 250 mM NaCl) and mixed with 80 µL of ANTI-FLAG® M2 affinity gel (Sigma-Aldrich). Immunoprecipitation was performed at room temperature on a rotating mixer for 1.5 h. Then affinity resin was washed consecutively with the following solutions 1 mL each: TES buffer (10 mM Tris-HCl pH 7.5, 1 mM EDTA, 250 mM NaCl); twice with TESS buffer (10 mM Tris-HCl pH 7.5, 250 mM NaCl, 1 mM EDTA, 0.1% Tween-20, 0.05% SDS); and TE buffer (10 mM Tris-HCl pH 7.5, 1 mM EDTA). All wash procedures were carried out by resuspending the beads in the wash solution and inverting the tube several times, followed by brief centrifugation and removing the supernatant. For proteolysis and de-crosslinking after washing procedure, affinity resin was diluted with 200 µL of TES buffer, proteinase K (Sigma-Aldrich) was added up to 0.5 mg/mL and samples were incubated at 55°C for 4 h. Next, agarose gel was removed by centrifugation (2000 g, 2 min, 25°C) and IP-DNA was finally purified from supernatant using AMPure XP magnetic beads (Beckman Coulter). Enrichment was assessed with qPCR for several genomic loci (**Table S1**). For further details see **Supplementary Materials & Methods**.

NGS libraries were prepared using TruSeq kit (Illumina). DNA sequencing was performed by Illumina NextSeq 75+75 bp paired-end protocol. Libraries preparation and sequencing were performed at Skoltech Genomics Core Facility.

ChIP-Seq experiments were performed in triplicate.

### EcTopoI ChIP-Seq with RNAP inhibition by rifampicin

ChIP-Seq experiments with inhibition of RNAP with rifampicin (Rif) were performed as described above, except that cells were pretreated with 100 µg/mL Rif for 20 min before fixation with formaldehyde. Subsequent steps of samples preparation were identical to ChIP-Seq of untreated cells.

### EcTopoI ChIP-Seq with overexpression of EcTopoI 14kDa CTD

40 mL of exponentially growing culture of *E. coli* DY330 *topA-SPA* cells harboring pCA24 14kDa CTD plasmid were cultivated in LB supplemented with 50 µg/mL kanamycin and 34 µg/mL chloramphenicol until reaching OD_600_∼0.2. Then, IPTG was added to final concentration 1 mM to induce CTD expression and the cultivation was proceeded for 1 h before fixing with formaldehyde. The following steps of samples preparation were continued identical to ChIP-Seq for untreated cells.

### E. coli RNAP ChIP-Seq

For TAP-tagged RNAP-ChIP, we used *E. coli* DY330 strain (Dharmacon) encoding RpoC fused with TAP-tag (Butland *et al*., 2005). Cell culture (V=200 mL) grown to mid-exponential phase OD_600_∼0.5-0.7 in LB containing 50 µg/mL of kanamycin was crosslinked by adding formaldehyde to the final concentration of 1%, followed by incubation for 20 min at room temperature, with agitation. Crosslinking was stopped by the addition of sterile glycine to a final concentration of 0.25 M. Cells were incubated for 20 min at room temperature and harvested by centrifugation at 3000 g, 10 min, 4°C. Cell pellets were washed twice with 20 mL of ice-cold 20 mM Tris-HCl pH 7.6, containing 60 mM NaCl (TBS), resuspended in 0.5 mL of ChIP Lysis Buffer 1 (10 mM Tris-HCl pH 8.0, 50 mM NaCl, 10 mM EDTA, 20% sucrose), followed by addition of 0.5 mL of 2x ChIP Lysis Buffer 2 (200 mM Tris-HCl pH 8.0, 600 mM NaCl, 4% Triton X-100), containing RNaseA (1 µg/mL). Resuspended cells were pre-lysed by the addition of 1 µg of Lysozyme and incubated at 37°C for 10 min, followed by sonication in a 2-mL Eppendorf tube in an ice-water bath to achieve a maximum yield of 300-400 bp fragments. Lysates were clarified by centrifugation at 13000 g for 10 min at 4°C, and the resulting supernatant was used for further analysis.

For the Input DNA fragment size analysis, a 100 µL aliquot of the clarified lysate was first treated with proteinase K (Thermo-Fisher Scientific) at 0.125 mg/mL in the presence of 1% SDS for 4 h at 37°C followed by DNA de-crosslinking by incubation at 65°C for 6 h. The Input DNA material was purified using Qiagen PCR purification kit (Qiagen) and quantified using Nano-Drop spectrophotometer. The DNA fragment size distribution was assessed by DNA electrophoresis.

For the preparation of RNAP-ChIP DNA, the remaining lysate (∼900 µL) was purified by immunoprecipitation (IP). The lysate was mixed with 10 µL of IgG-agarose (GE Healthcare) and incubated at 4°C overnight on a rotating mixer in the presence of Protease Inhibitor Cocktail (Sigma). Then affinity resin was washed consecutively with 1 mL each of the following solutions: 40 mM Tris-HCl pH 7.9, 0.5% Tween-20, 2 M NaCl; 40 mM Tris-HCl pH 7.9, 0.5% Tween-20, 1 M NaCl; 40 mM Tris-HCl pH 7.9, 0.5% Tween-20, 200 mM NaCl; and twice with RIPA buffer (50 mM Tris-HCl pH 7.4, 140 mM NaCl, 1% NP-40, 0.1% deoxycholate, 0.1% SDS). All wash procedures were carried out by resuspending the beads in the wash solution and inverting the tube several times, followed by brief centrifugation and removing the supernatant by vacuum aspiration. The last wash was done for 20 min at 4°C on a rotary mixer. The RNAP-DNA crosslinks were eluted by incubation with ChIP Elution Buffer (10 mM Tris pH 8.0, 30 mM EDTA, 1% SDS) at 65°C for 4 hours (or O/N) on a shaker at 800-900 rpm. The resulting material was treated with Proteinase K (Thermo Fisher Scientific) (0.2 mg/mL) at 37°C for 3-5 h, followed by IP-DNA decrosslinking by incubation at 95°C for 2h. The IP-DNA was finally purified using ChIP DNA Cleaning & Concentrator kit (Zymo Research).

A 50-100 ng of IP-DNA or Input-DNA were end repaired by a mix of T4 DNA polymerase (NEB), T4 PNK (NEB), and Klenow DNA polymerase (NEB) and purified by Qiaquick PCR DNA purification kit (Qiagen). The eluted DNA material was A-tailed by Klenow Fragment (3C to 5C exo minus) (NEB) followed by purified by MinElute PCR purification Kit (Qiagen). Illumina Multiplex Adapters (MPA) were ligated with Quick DNA ligase (NEB) and DNA was purified by AMPure XP beads (Beckman Coulter). Resulting library was separated by agarose electrophoresis with subsequent size selection of DNA bands corresponding to 220 bp, which were excised by Gel X-tracta tool (USA scientific) and purified with Gel Extraction Kit (Qiagen). The purified DNA material was PCR-amplified (18 cycles) using Phusion polymerase (NEB) and purified by MinElute PCR purification kit (Qiagen).

NGS libraries were prepared using TruSeq kit (Illumina). DNA sequencing was performed by Illumina HiSeq 50+50 bp paired-end protocol.

### ChIP-Seq data analysis

Reads, filtered and trimmed with Trimmomatic (Bolger, Lohse and Usadel, 2014), were aligned to the *E. coli* W3110 MuSGS genome (*E. coli* W3110 genome with the insertion of *cat*-Mu SGS cassette may be downloaded from GEO: GSE95567) using BWA-MEM (Li and Durbin, 2010). BAM and bed files were prepared with Samtools (Li *et al*., 2009) and visualized in IGV (Thorvaldsdóttir, Robinson and Mesirov, 2013). For EcTopoI ChIP-Seq data peak calling was performed with MACS2 (Zhang, Liu, Clifford A. Meyer, *et al*., 2008) with the following parameters: nomodel, Q-value<0.001. Motif identification was performed by ChIPMunk (Kulakovskiy *et al*., 2010) and visualization by WebLogo (Crooks *et al*., 2004). Fold enrichment tracks were further analyzed using custom python scripts (https://github.com/sutormin94/TopoA_ChIP-Seq). Detailed analysis is described in **Supplementary Materials & Methods**.

Genome annotation with open reading frames (ORFs) for *E. coli* W3110 genome was obtained from Ensembl Bacteria (Howe *et al*., 2020). Annotation of operons for *E. coli* W3110 is based on DOOR database (Mao *et al*., 2009). Annotation of transcription units (TUs) for *E. coli* MG1655 was taken from EcoCyc database and transferred to *E. coli* W3110 (Karp *et al*., 2018). Information about subcellular localization of *E. coli* proteins was taken from PSORTdb 4.0 (Peabody *et al*., 2016). Annotation of promoters and transcription factor sites was taken from RegulonDB (Santos-Zavaleta *et al*., 2019).

### E. coli total RNA-Seq and data analysis

Total RNA was extracted with ExtractRNA reagent (Evrogen) from 2 mL of exponentially growing *E. coli* DY330 culture in LB medium, when OD_600_ reached ∼0.6. RNA sample was treated with DNAse I (ThermoFisher Scientific) and purified with RNAClean XP beads (Beckman Coulter). Sequencing libraries were prepared without rRNA depletion using NEBNext Ultra II Directional RNA Library kit (NEB) with the following modifications: 10 min of fragmentation instead of 15 and 10 PCR cycles. Then they were quantified using qPCR, pooled in equal amounts, diluted to 2 nM and sequenced on HiSeq4000 instrument (Illumina, USA) with 50 bp long reads protocol. Libraries preparation and sequencing were performed at Skoltech Genomics Core Facility. RNA-Seq was performed in triplicate.

Raw reads were trimmed and filtered with Trimmomatic (Bolger, Lohse and Usadel, 2014) and aligned to the *E. coli* W3110 MuSGS genome (*E. coli* W3110 genome with the insertion of *cat*-Mu SGS cassette may be downloaded from GEO: GSE95567) using BWA-MEM (Li and Durbin, 2010). BAM and bed files were prepared with Samtools (Li *et al*., 2009). RSeQC package was used for FPKM and genes expression level calculation (Wang, Wang and Li, 2012).

### Strand specific EcTopoI Topo-Seq and data analysis

*E. coli* DY330 *topA-SPA* were transformed with pBAD33 topA_strepII G116S M320V plasmid. Culture grown overnight at 37°C and with shaking 180 rpm in LB supplemented with chloramphenicol (34 µg/mL) and 0.5% glucose was diluted with fresh medium, supplemented with the antibiotic and glucose, to get 100 ml. Cultivation continued until reaching OD_600_=0.4, then it was bisected, and one half (+Ara) was induced by addition of arabinose to 10 mM. The other half of the culture served as a non-induced control (-Ara). 30 min after the induction cells were harvested by centrifugation (3000 g). Cell pellet at this step can be frozen in liquid N_2_ and stored at -80°C until further processing. The cell pellet was resuspended in 1 mL of strep-Tactin lysis buffer (50 mM Tris HCl pH 8.0, 150 mM NaCl) containing protease inhibitors cocktail (1x concentration, cOmplete ULTRA, Sigma-Aldrich) and RNase A (0.1 mg/ml, Thermo Scientific). To disrupt cells and shear DNA, cells were sonicated in a 1.5 mL Eppendorf tube in an ice-water bath to achieve 100-1000 bp fragments range. Lysates were clarified by centrifugation at 8000 g for 5 min at 4°C, and the resulting supernatant was used for further analysis.

For Input DNA (+Ara-IP and -Ara-IP samples) 100 µL of the lysate was treated with proteinase K (Thermo-Fisher Scientific) for 2 h at 55°C and DNA was purified using DNA clean-up kit (Thermo Fisher Scientific). DNA fragmentation range was assessed by electrophoresis.

For cleavage complexes immunoprecipitation (+Ara+IP and -Ara+IP samples), 900 µL of the remaining lysate was mixed with 80 μL of strep-Tactin Superflow Plus affinity resin (Qiagen) pre-equilibrated with strep-Tactin lysis buffer supplemented with 0.05% SDS. After 1 h of incubation at 25°C with constant rotation the resin was washed three times with strep-Tactin lysis buffer, the proteins bound were eluted using 100 μL of elution buffer (50 mM Tris HCl pH 8.0, 150 mM NaCl, 2.5 mM desthiobiotin). 20 µL of the elution were subjected for analysis using SDS PAGE. Remaining 80 µL of the elution were treated with proteinase K (Thermo-Fisher Scientific) overnight at 50°C. IP-DNA was purified using AMPure XP magnetic beads (Beckman Coulter). Topo-Seq experiments were performed in triplicate.

NGS libraries were prepared using strand-specific Accel NGS 1S kit (Swift Bioscience) suitable for damaged DNA. DNA sequencing was performed by Illumina NextSeq 150+150 bp paired-end protocol. Libraries preparation and sequencing were performed at Skoltech Genomics Core Facility.

Reads, filtered and trimmed with Trimmomatic (Bolger, Lohse and Usadel, 2014), were aligned to the *E. coli* W3110 MuSGS genome (*E. coli* W3110 genome with the insertion of *cat*-Mu SGS cassette may be downloaded from GEO: GSE95567) using BWA-MEM (Li and Durbin, 2010). The number of DNA fragments’ 3’-ends was calculated per position (N3E) separately for forward and reverse strands, based on read alignments stored in SAM files. The tracks were scaled by the total number of aligned reads to get normalized coverage across samples and biological replicates were averaged. After that -IP tracks (+Ara-IP and -Ara-IP) were subtracted from +IP (+Ara+IP and -Ara+IP respectively) tracks strand-wise resulting in +Ara and -Ara tracks. Finally, -Ara tracks were subtracted from +Ara tracks strand-wise to get enriched signal. Resultant tracks were further analyzed using custom python scripts (https://github.com/sutormin94/TopoI_Topo-Seq).

### Strand-specific DRIP-Seq and data analysis

Single colony of *E. coli* DY330 *topA*-SPA was inoculated into 5 mL of LB supplemented with 50 µg/mL kanamycin and 0.5% glucose and grown overnight. The starter culture (0.5 mL) was diluted 100 times with 50 mL of fresh LB with 0.5% glucose and cultivation continued at 37°C and shaking till mid-exponential phase (OD_600_∼0.6). Cells were harvested by centrifugation and total nucleic acid was purified using GeneJET Genomic DNA purification kit (ThermoFisher) according to the manufacturer’s protocol but omitting the RNAse A treatment. Extracted nucleic acid was sheared up to 150 bp fragments using Covaris ultrasonicator.

10 µg of S9.6 antibodies (Kerafast, ENH001) were incubated with 20 µL of protein-G Sepharose (Bialexa) equilibrated with PBS in 70 µL of IP-buffer (10 mM Tris-HCl pH 7.5, 1 mM EDTA, 0.1% sodium deoxycholate, 1% Triton X-100, 140 mM NaCl) for 2 h at 4°C with gentle shaking. After the pre-incubation, an aliquot (containing 6 µg of sheared DNA, concentrations measured using Qubit 1X dsDNA HS Assay) was added and the immunoprecipitation continued for 4 h. Then, the resin was washed with 1 mL of IP-buffer, 1 mL of IPS-buffer (10 mM Tris-HCl pH 7.5, 1 mM EDTA, 0.1% sodium deoxycholate, 1% Triton X-100, 500 mM NaCl) and twice with 1 mL of Wash-buffer (10 mM Tris-HCl pH 8, 1 mM EDTA, 0.5% sodium deoxycholate, 1% Triton X-100, 250 mM NaCl). TES-buffer (10 mM Tris-HCl pH 7.5, 1 mM EDTA, 250 mM NaCl) was mixed with the washed resin to get 200 µL of slurry and proteinase K (ThermoFisher) was added to a final concentration 0.5 mg/mL. Proteolysis was performed at 55°C for 1 h. After that, the resin was spun down by centrifugation and nucleic acids were purified from supernatant with AMPure XP magnetic beads (Beckman Coulter). As a control, an aliquot of sonicated nucleic acids (containing 6 µg of sheared DNA, concentrations measured using Qubit 1X dsDNA HS Assay) was treated with 5U of RNAse HI (NEB) in RNAse HI buffer for 30 min at 37°C and then nucleic acids were purified with GeneJET Gel Extraction and DNA cleanup micro kit (General cleanup protocol, ThermoFisher). IP for treated aliquot using S9.6 antibodies and nucleic acid purification were performed as described for the untreated aliquot.

DRIP with cells expressing EcTopoI 14kDa CTD was performed similarly but with *E. coli* DY330 *topA*-SPA harboring pCA24 14kDa CTD plasmid. Cultures were grown in LB supplemented with chloramphenicol 34 µg/mL and 0.5% glucose. 14kDa CTD overexpression was induced with 1mM IPTG at OD_600_∼0.2 and cultivation continued for 1 h. DRIP with cells treated with RNAP inhibitor rifampicin was performed similarly but rifampicin (SigmaAldrich) was added in a concentration 100 µg/mL at OD_600_∼0.6 (after 1 h min after 14kDa CTD induction) and cultivation continued for 20 min. DRIP-Seq experiments were performed in triplicate.

NGS libraries were prepared using strand-specific Accel NGS 1S kit (Swift Bioscience) suitable for single-stranded DNA. DNA sequencing was performed using Illumina NextSeq with 75+75 bp paired-end protocol. Libraries preparation and sequencing were performed at Skoltech Genomics Core Facility.

Reads, filtered and trimmed with Trimmomatic (Bolger, Lohse and Usadel, 2014), were aligned to the *E. coli* W3110 MuSGS genome (*E. coli* W3110 genome with the insertion of *cat*-Mu SGS cassette may be downloaded from GEO: GSE95567) using BWA-MEM (Li and Durbin, 2010). For samples with overexpression of EcTopoI 14kDa CTD reads were also aligned to pCA24 14kDa CTD plasmid. BAM and bed files were prepared with Samtools (Li *et al*., 2009) and visualized in IGV (Thorvaldsdóttir, Robinson and Mesirov, 2013). Strand-specific coverage for forward and reverse strands were obtained based on read alignments stored in SAM files. The tracks were scaled by the total number of aligned reads to get normalized coverage across samples and biological replicates were averaged. Then coverage depth for reverse strand was subtracted from coverage for the forward strand. After the scaling, tracks of control samples (treated with RNAse HI) were subtracted from corresponding experimental tracks. Resultant tracks were further analyzed using custom python scripts (https://github.com/sutormin94/E_coli_DRIP-Seq_analysis).

### EMSA

26mer oligonucleotides (Consensus, Poly-T, and Random) and their complementary sequences were synthesized in non-labeled and 5’-Cy5-labeled forms (Syntol) (**Table S1**). For binding evaluation 0.4 pmoles of Cy5-labeled oligonucleotide was mixed with increasing amount of purified EcTopoI (0-16 pmoles; see **Supplementary Materials & Methods** for the purification procedure) in the binding buffer (10 mM Tris-HCl pH 7.5, 50 mM NaCl, 6 mM MgCl_2_). 20 µL reactions were incubated at 37°C for 10 min and DNA was separated in 10% acrylamide gel (acrylamide/bisacrylamide 29:1) prepared on 1xTGB with magnesium (25 mM Tris-HCl, 250 mM glycine, 6 mM MgCl2, pH 8.3). Separation was performed in TGB with magnesium at room temperature at 100 V. Cy5-labeled bands were visualized in ChemiDoc imaging system (Biorad).

For competition experiments 0.4 pmoles of labeled oligonucleotide was mixed with saturating amount of EcTopoI (16 pmoles) and increasing amount of non-labeled competitor oligonucleotide (0-51.2 pmoles – molar excess over labeled oligonucleotide up to 128x) in the binding buffer (10 mM Tris-HCl pH 7.5, 50 mM NaCl, 6 mM MgCl_2_). Reactions and DNA separation were performed as described for oligonucleotides.

*dps*, *potF* or *nuoN* DNA fragments were PCR-amplified from *E. coli* DY330 genomic DNA (for primers see **Table S1**) and purified with GeneJET Gel Extraction and DNA cleanup micro kit (PCR cleanup protocol, ThermoFisher). For binding 2.85 pmoles of DNA fragment was mixed with increasing amount of purified EcTopoI (0-16 pmoles and 0-32 pmoles correspondingly) in binding buffer (10 mM Tris-HCl pH 7.5, 50 mM NaCl, 0.1 mM EDTA) (Narula and Tse-Dinh, 2012). 20 µL reactions were incubated at 37°C for 10 min and DNA was separated in 10% acrylamide gel (acrylamide/bisacrylamide 29:1) prepared on 1xTAE buffer. Separation was performed in TAE at room temperature at 100 V. For DNA visualization gel was stained with EtBr.

### Microscale thermophoresis (MST)

10 fmoles of Cy5-labeled 26-mer oligonucleotide (Consensus, Poly-T, Random and complementary sequences - **Table S1**) were mixed with increasing amount of purified EcTopoI (0-17 pmoles) in the binding buffer (10 mM Tris-HCl pH 7.5, 50 mM NaCl, 6 mM MgCl_2_, 0.05% Tween 20). 20 µL reactions were incubated at room temperature for 5 min and 5 µL of the mixture was loaded into NT.115 capillaries (NanoTemper). MST was performed in Monolith NT.115 (NanoTemper) at 22°C, excitation power 100% and MST power 40%. Binding curve fitting, K_D_ measurement (K_D_ model) and statistic evaluation were performed automatically with default parameters using MO.Affinity software (NanoTemper).

### Microscopy

Induced and non-induced *E. coli* DY330 *topA-SPA* pCA24 14kDa CTD cells were cultivated as described for EcTopoI ChIP-Seq. *E. coli* BW25113 cultures were grown in LB till OD_600_∼0.6. Cells were spotted to agarose pads (1.2% agarose in PBS) and imaged at ×100 magnification using Nikon Eclipse Ti microscope equipped with the Nikon Plan Apo VC 100X1.40 oil objective and Nikon DS-Qi2 digital monochrome camera. Images were processed using ImageJ software (Schneider, Rasband and Eliceiri, 2012).

## Results

### EcTopoI is widely distributed over the *E. coli* genome, co-localized with RNAP, and enriched in regions with negative supercoiling

The topoisomerase I distribution along the *E. coli* chromosome was determined using ChIP-Seq with a DY330 strain derivative carrying a fusion of the *topA* gene with the SPA-tag encoding sequence (**Figure 2**, orange track). Three biological replicas were made and a good correspondence between them was observed (**Figure S1A**, Pearson correlation > 0.6). Using the MACS2 analysis pipeline (Zhang, Liu, Clifford A Meyer, *et al*., 2008), we detected 403 significantly enriched regions (e-value < 0.001) observed in all three replicas (**Figures S1B-E**). EcTopoI peaks tend to have a lower GC-content than the genome average (**Figure S2**) and a positive correlation between peaks log-fold enrichment and the AT-content was observed (Spearmen correlation 0.36, p-value 2.3e-5). The peaks appeared to be uniformly distributed over the entire chromosome. Of note, there was no notable enrichment of the EcTopoI signal at the terminator region of chromosome replication, in contrast to observations made for *M. smegmatis* topoisomerase I (Rani and Nagaraja, 2018).

**Figure 2.**
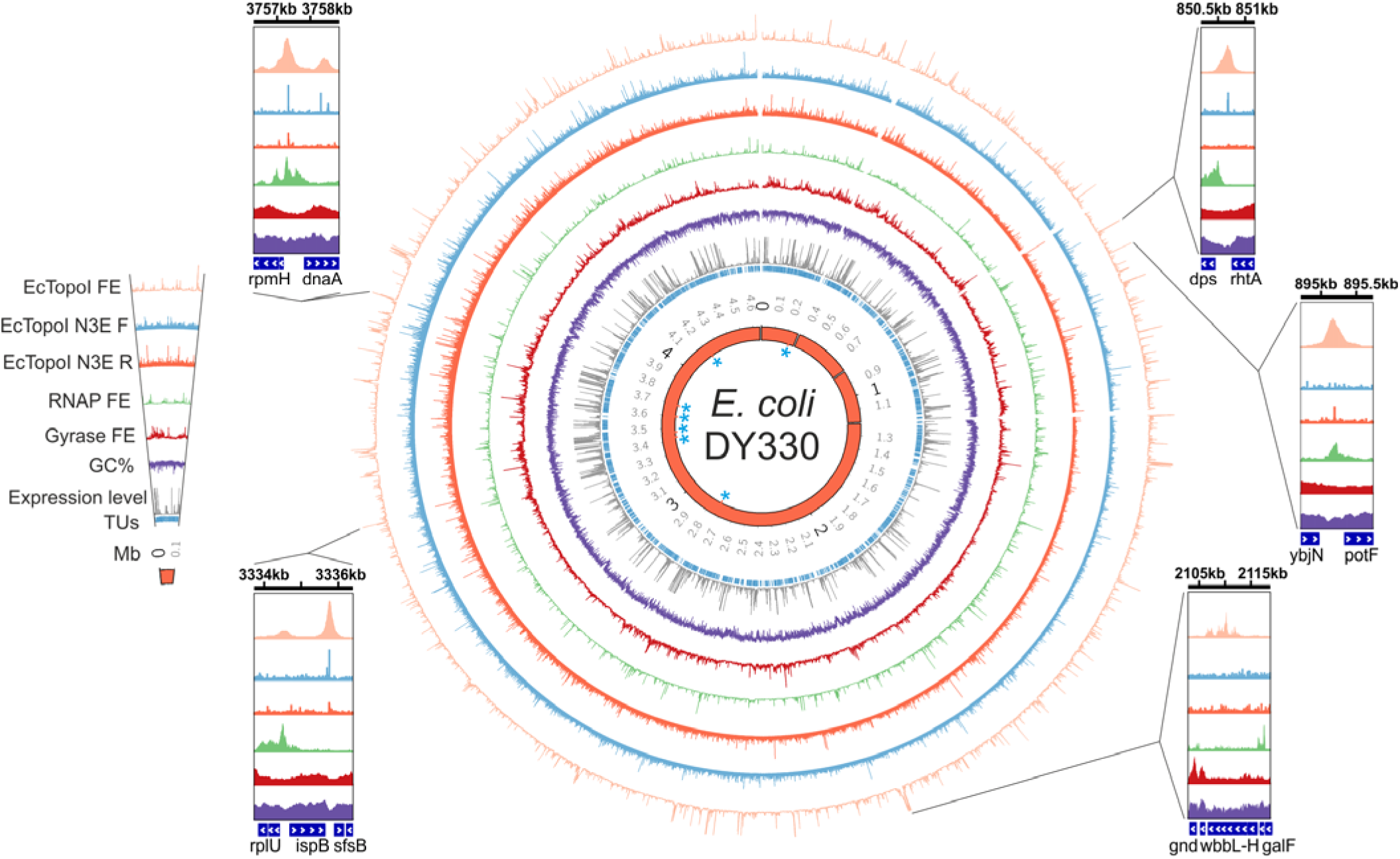
Distribution of EcTopoI, RNAP, and gyrase enrichment peaks over the *E. coli* chromosome. Circular maps demonstrate enrichment of EcTopoI (ChIP-Seq, orange; Topo-Seq, blue and red for two strands separately), RNAP (ChIP-Seq, green), and DNA gyrase (Topo-Seq, dark red). Additionally, GC% (purple) and mean expression level (RNA-Seq, grey) of transcription units (TUs annotation, inner blue circle) are shown. The rRNA operons are highlighted with blue asterisks in the inner red ring representing *E. coli* DY330 genome. The numbers on the outside of the ring give genome coordinates in megabase pairs (Mbs). Three gaps at around ∼0.3 Mb, ∼0.8 Mb, and ∼1.2 Mb correspond to deletions in the *E. coli* DY330 genome relative to the *E. coli* W3110 reference genome. Insets give a closer look into representative regions with high EcTopoI signals. Coordinates given in Kbs are indicated on the top of each inset. The maps were constructed with the Circos tool (Krzywinski *et al*., 2009), insets were prepared using the IGV (Thorvaldsdóttir, Robinson and Mesirov, 2013).

We next determined whether the EcTopoI ChIP-Seq signal overlaps with the RNAP signal, a result that might be expected based on the published data about the interaction between the two enzymes (Cheng *et al*., 2003). А ChIP-Seq experiment with a DY330 strain derivative carrying a fusion of the *rpoC* (RNAP β’ subunit) gene with a TAP-tag encoding sequence was performed (**Figure 2**, green track). The RpoC ChIP-Seq signal correlated well with published ChIP-Seq obtained for RpoB (β RNAP subunit) (Kahramanoglou *et al*., 2011) (Spearman correlation 0.59, **Figure S3E**). Overall, we found 3635 RpoC peaks with fold enrichment of at least 3, ∼25% of which overlapped with earlier reported RpoB peaks (Monte-Carlo simulation with 10000 iterations, p-value<1e-308, **Figures S3F** and **S4C**). 60% of topoisomerase peaks (243/403, Monte-Carlo simulation with 10000 iterations, p-value=4.9e-6, **Figure S4A**) overlapped with the RpoC peaks (**Figure 3A**). Consistently, enrichment of RpoC is significantly higher within the EcTopoI-enriched regions compared to the outside regions (Welch t-test, p-value<1e-308) and enrichment of EcTopoI is significantly higher inside the RpoC-occupied regions than outside of these regions (Welch t-test, p-value<1e-308) (**Figure 3B**). Colocalization between the RpoB and EcTopoI signals was also observed with a publicly available RpoB ChIP-Seq dataset for *E. coli* MG1655 (Kahramanoglou *et al*., 2011) (**Figures S3G-H**). Overall, we conclude that EcTopoI is significantly co-localized with RNAP on the *E. coli* chromosome in exponentially growing cells. We also performed an RNA-Seq experiment with *E. coli* DY330 (**Figure S3A-C**). As expected, a strong correlation was observed between the RpoC ChIP-Seq signal and transcription level (**Figure S3D**, Spearman correlation 0.55, p-value 1.4e-133). The signal of EcTopoI was also roughly proportional to transcript abundance, with highest enrichment values observed for 200 most highly-expressed transcription units (HETUs, expression level > 31 FPKM) and, particularly, for rRNA operons (**Figures S8.II** and **III**). In contrast, little or no EcTopoI enrichment was observed for 200 least expressed TUs (LETUs, expression level < 0.31 FPKM) (**Figure 3C**).

**Figure 3.**
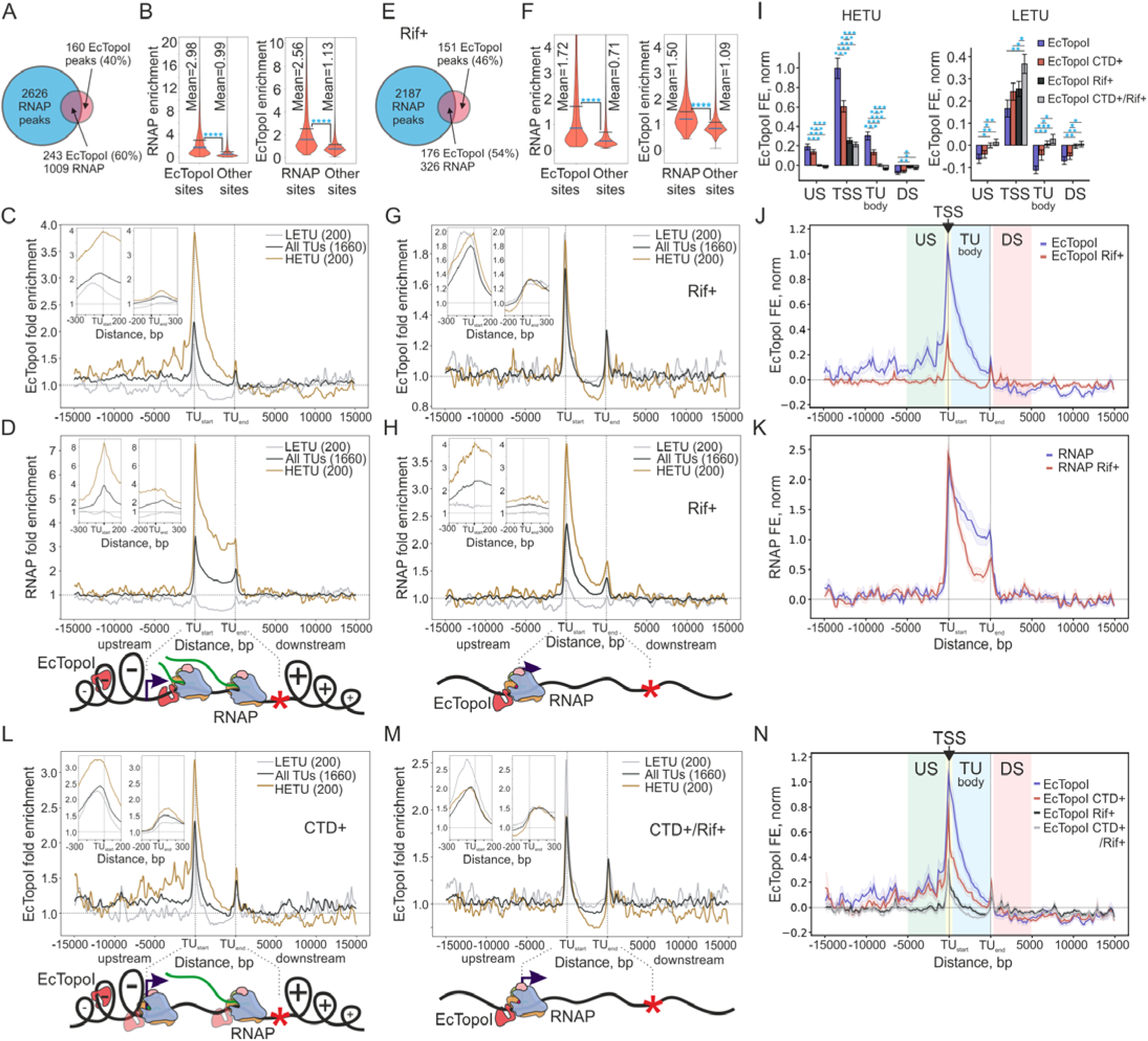
EcTopoI is associated with RNAP and regions of expected transcription-induced negative supercoiling. (**A**) Number of EcTopoI peaks overlapping with RNAP peaks. The Venn diagram represents an overlap of EcTopoI peaks (403 in total) and RNAP peaks (3635 totally). (**B**) Violin plots of RNAP enrichment in EcTopoI peaks and in outside regions (left). Violin plots of EcTopoI enrichment in RNAP peaks and in outside regions (right). The means and medians are indicated by black and blue lines, respectively. Statistically significant differences between means (t-test, p-value<<0.05) are indicated by asterisks. (**C**) Metagene plot of EcTopoI enrichment within TUs, and in their upstream (left) and downstream (right) regions. Enrichment is shown for all TUs (black curve), highly-expressed (HETU, orange curve), and least-expressed (LETU, grey curve) sets. The number of TUs in each group is indicated in parentheses. The two insets show zoom-in views of EcTopoI enrichment near transcription start (TU start) and termination (TU end) sites. (**D**) A metagene plot of RNAP enrichment within TUs and in their upstream and downstream regions. Analysis and groups of TUs are same as in **C**. A graphical representation of the Liu & Wang twin-domain model (Liu and Wang, 1987) and localization of RNAP and EcTopoI according to the metagene plots is shown below the panel. (**E**) Number of EcTopoI peaks overlapping with RNAP peaks in cells treated with Rif. The Venn diagram represents an overlap of EcTopoI peaks (327 in total) and 2513 RNAP peaks. (**F**) Violin plots of RNAP enrichment in EcTopoI peaks and in outside regions (left) and of EcTopoI enrichment in RNAP peaks and in outside regions (right) for cells treated with Rif (an RpoC ChIP-chip dataset for cells treated with Rif was taken from (Mooney *et al*., 2009)). (**G**) A metagene plot of EcTopoI enrichment within TUs, in their upstream and downstream regions for cells pre-treated with RNAP inhibitor Rif before ChIP-Seq. (**H**) A metagene plot of RNAP enrichment within TUs, in their upstream and downstream regions for cells treated with Rif. A graphical representation of the localization of RNAP and EcTopoI according to metagene plots is shown below the panel. (**I**) Comparison of EcTopoI enrichment for untreated condition (blue bars), cells pretreated with Rif (black bars), cells overexpressing 14kDa CTD (red bars), and cells overexpressing 14kDa CTD followed by Rif treatment (grey bars) for HETUs (left) and LETUs (right). Enrichment was quantified for normalized tracks for regions near the transcription start sites (TSS, ±200 bp from transcription start site), 5 Kbs upstream regions (US), 5 Kbs downstream regions (DS), and TU bodies (TU). Differences in enrichment were tested using Welch t-test. Significance is indicated by asterisks. Error bars are represented by ±SEM. (**J**) A metagene plot of normalized EcTopoI enrichment for untreated condition (blue curve) and for cells pretreated with Rif (red curve) for HETUs. Confidence bands are represented by ±SEM. Regions used for quantification of enrichment in panel **I** (US, TSS, TU body, and DS) are shown on the plot as colored areas. (**K**) A metagene plot of normalized RNAP enrichment for untreated condition (blue curve) and for cells pretreated with Rif (red curve) for HETUs. (**L**) A metagene plot of EcTopoI enrichment in TU bodies and, upstream and downstream regions in cells overexpressing 14kDa CTD. A graphical representation of the localization of RNAP and EcTopoI according to the metagene plot is shown above the panel. (**M**) A metagene plot of EcTopoI enrichment in TU bodies, and upstream and downstream regions in cells overexpressing 14kDa CTD followed by Rif treatment. A graphical representation of the localization of RNAP and EcTopoI according to the metagene plot is shown below the panel. (**N**) A metagene plot for HETUs demonstrating normalized EcTopoI enrichment for untreated condition (blue curve), cells with induced 14kDa CTD (red curve), cells treated with Rif (black curve), and a combination of CTD induction and Rif treatment (grey curve). Confidence bands are represented by ±SEM. Regions used for quantification of enrichment in panel **I** (US, TSS, TU body, and DS) are shown by colored areas on a plot.

Enrichment of EcTopoI and RNAP within transcription units (TUs), and in upstream and downstream regions was next analyzed (**Figures 3C, D**, and **S3I**). Metagene analysis indicated colocalization of EcTopoI and RNAP within the TU bodies and, particularly, close to the transcription start sites (TSS), where the highest enrichment for both enzymes was observed. A decreasing RNAP enrichment gradient towards the ends of TUs, presumably caused by premature transcriptional termination (De Smit *et al*., 2008; Zhu *et al*., 2019), was observed. A gradient with a similar slope was detected for EcTopoI enrichment, suggesting that EcTopoI either directly follows elongating RNAPs or physically associates with the enzyme.

EcTopoI accumulated upstream but was depleted downstream of TUs, a result that is consistent with the predictions of the Liu & Wang twin-domain model that posits accumulation of negative supercoiling (a substrate of TopoI) upstream, i.e., behind the elongating RNAP (Liu and Wang, 1987). Excessive enrichment of EcTopoI over the background could be tracked up to 12-15 Kbs upstream of TSS for HETUs (**Figure 3C**), suggesting that negative supercoiling can diffuse over significant lengths of *E. coli* chromosome. A small peak of EcTopoI and RNAP enrichment at TU ends may correspond to enrichment at promoter regions of closely packed adjacent genes or result from physical association of the two enzymes at transcription termination sites.

Overall, our observations strongly support the association of EcTopoI with RNAP at transcription start sites and within TUs and with negatively supercoiled DNA upstream of actively transcribed genes.

### The RNAP inhibitor rifampicin causes EcTopoI re-localization to promoter regions

If EcTopoI interacts with RNAP, then it should redistribute to promoters upon the treatment with rifampicin (Rif), an inhibitor that prevents RNAP escape into elongation (Campbell *et al*., 2001). On the other hand, if EcTopoI association with extended regions upstream of TUs were driven by excessive transcription-generated negative supercoiling, then Rif treatment shall abolish this association. To test these predictions, we performed EcTopoI ChIP-Seq in cells treated with Rif prior to formaldehyde fixation. According to metagene analysis, EcTopoI enrichment along the lengths of HETUs bodies disappeared in Rif-treated samples reaching lower than background enrichment values (**Figure 3G**). This corresponded with the disappearance of elongating RNAP from TU bodies in Rif-treated samples (**Figures 3H, K**, an RpoC ChIP-chip dataset for cells treated with Rif was taken from (Mooney *et al*., 2009)). Association of EcTopoI with upstream regions of HETUs was also abolished upon Rif treatment (**Figures 3G, J**). Yet, the enrichment of EcTopoI on promoter regions of HETUs remained, although at lower levels compared to untreated control (**Figures 3I, J**). The decrease may be caused by dissipation of transcription-induced negative supercoiling and/or by redistribution of RNAP. The latter scenario is supported by the observation that in Rif-treated samples enrichment of both EcTopoI and RNAP is increased at LETU promoters (**Figures 3H, I**). Be that as it may, EcTopoI and RNAP remained co-localized in Rif-treated cells (Welch t-test, p-value<1e-308, **Figures 3E, F**), sharing a significant number of enrichment peaks (Monte-Carlo simulation with 10000 iterations, p-value<1e-308, **Figure S4D**). Consistent with the re-localization of EcTopoI to promoters, EcTopoI peaks found by MACS2 in Rif-treated cells were narrower (median width 311 bp) and more AT-rich (43.5% GC) than peaks in untreated samples (**Figure S2B**). Overall, these results further support EcTopoI interaction with elongating RNAP, promoter initiation complexes, and regions upstream of transcribed genes.

### Overexpression of EcTopoI CTD impairs interaction with RNAP

The 14 kDa CTD of EcTopoI was previously shown to interact with *E. coli* RNAP *in vitro* (Cheng *et al*., 2003), however, the physiological role of this interaction is unknown. As expected, we observed that RNAP can be purified by affinity chromatography from cells encoding SPA-tagged EcTopoI and from cells overexpressing His-tagged CTD (**Figure S5C**) by using EcTopoI or CTD as baits (**Figures S5D** and **E**). Overexpression of CTD decreased the amount of RNAP co-purified with SPA-tagged EcTopoI from *E. coli* DY330 cells, while overexpression of the GFP control had no such effect (**Figure S5F**) (see **Supplementary Materials & Methods** for procedure details). We therefore reasoned that overproduction of CTD may impair the EcTopoI-RNAP interaction *in vivo* and thus change the distribution of EcTopoI. Accordingly, we carried out EcTopoI ChIP-Seq experiments with cells overexpressing the EcTopoI CTD. To avoid possible biases caused by toxicity of prolonged overexpression of the CTD (**Figures S5A-B**), we grew cells until OD_600_∼0.2, induced them with high, 1 mM, concentration of IPTG for one hour and then performed ChIP-Seq. As can be seen from **Figure 3N**, CTD overproduction indeed decreased EcTopoI enrichment in TUs bodies and in promoter regions (**Figure 3I**). In contrast, EcTopoI enrichment in upstream regions of TUs was similar in samples from cells overproducing CTD and control samples (**Figures 3N, I**). This enrichment was dependent on the level of transcription (**Figure 3L**) and extended up to 12 Kbs for HETUs, implying that it was caused by transcription-induced negative supercoiling. It therefore follows that CTD does not compete with EcTopoI for the binding to negatively supercoiled DNA but strongly affects EcTopoI interaction with RNAP. Treatment of CTD-expressing cells with Rif led to EcTopoI enrichment profiles that were similar to those observed for Rif-treated control (compare **Figure 3M** with **Figure 3G**). Additionally, using RT-qPCR, we showed that CTD overexpression does not affect transcription elongation (**Figures S5H-I**).

### EcTopoI is recruited to chromosomal regions with excessive negative supercoiling surrounded by topological barriers

EcTopoI distribution in 1529 *E. coli* intergenic regions (IRs, **Figure S6A**) was examined in more detail. We found high levels of enrichment at IRs flanked by highly-transcribed genes and/or having a high level of RNAP enrichment (**Figures S6B**, **C**). Independently of RNAP enrichment/expression levels, high levels of EcTopoI enrichment were characteristic for IRs that *i*) were located between divergently transcribed genes (**Figures S6D**), *ii*) harbored transcription factor-binding sites (**Figure S6E**, particularly high enrichment of EcTopoI was associated with IRs harboring sites of Cra and FNR binding, see **Figure S7**), and *iii*) were flanked by genes coding for membrane proteins (**Figures S6F** and **G**). IRs with the highest level of EcTopoI enrichment fulfilled all three criteria and were located between highly transcribed genes (**Figures 4A** and **S6H-I**).

**Figure 4.**
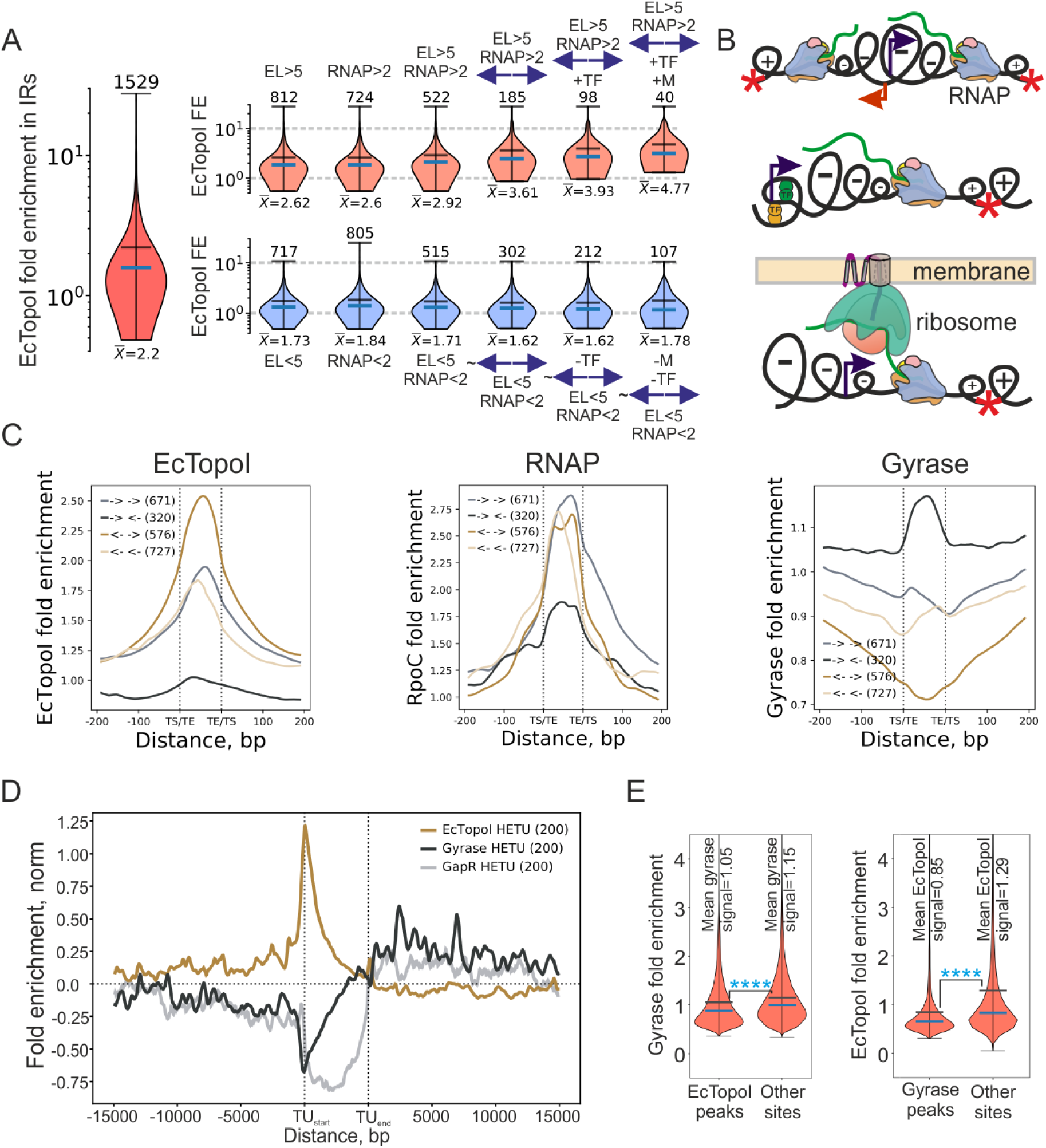
Features of intergenic regions associated with the increased enrichment of EcTopoI and mutual exclusion of EcTopoI and DNA gyrase genome-wide. (**A**) Cumulative effect of transcription and local topological borders on EcTopoI signal in IRs. Expression level of adjacent genes (EL), fold enrichment of RNAP (RNAP), orientation of adjacent genes (indicated by arrows), presence of annotated sites of transcription factors (TF), and localization of proteins encoded by flanking genes to the membrane (M) were tested for association with EcTopoI enrichment. Fold enrichment of EcTopoI in 1529 IRs analyzed (on the left) is compared with EcTopoI enrichment in IRs defined by features positively associated with EcTopoI signal (top row of violin plots), and with EcTopoI enrichment in IRs defined by features negatively associated with EcTopoI signal (bottom row of violin plots). Positively associated features are: high transcription level and high level of RNAP enrichment (EL>5, RNAP>2), - divergently transcribed genes, +TF – at least one annotated site of a transcription factor is present in an IR, M – at least one gene flanking IR encodes a membrane protein. Negatively associated features are: low level of transcription and RNAP enrichment (EL<5, RNAP<2), ∼ - genes separated by an IR are NOT in a divergent orientation, -TF – no annotated sites of transcription factors binding are present, -M – genes flanking an IR do not encode membrane proteins. Means are shown with horizontal black lines and numeric values are shown below each violin-plot, medians are shown as blue horizontal lines; vertical axes are log-scaled. (**B**) Graphical representation of topological borders that trap negative supercoiling: divergent orientation of genes, complexly organized promoters, transertion. (**C**) Meta-intergene plots of EcTopoI (on the left), RNAP (in the center), and gyrase (on the right) enrichments in IRs. IRs were classified according to orientation of flanking genes. Number of regions comprising a group is indicated in brackets. (**D**) Comparison of EcTopoI ChIP-Seq data (CTD-/Rif- conditions), gyrase Topo-Seq data (experiments with ciprofloxacin (Sutormin *et al*., 2019)), and GapR-Seq (Guo *et al*., 2021). Metagene analysis performed for a HETU set. (**E**) Violin plots of gyrase enrichment in EcTopoI peaks and outside of these regions (left) and of EcTopoI enrichment in gyrase peaks and outside of these regions (right). Mean and median are indicated by black and blue lines, respectively. Statistically significant difference between means (t-test, p-value<<0.05) is indicated by asterisks.

Since, meta-intergene analysis demonstrated that IRs flanked by divergent genes exhibit, on average, had a much higher EcTopoI signal than those between convergent genes (**Figure 4C**), we propose that EcTopoI is preferentially enriched at negative supercoils trapped by local topological borders, which are formed by divergent transcription from highly complex promoters and by coupled transcription and translation/polypeptide chain translocation into membrane (transertion) (Woldringh, 2002) of polypeptides into cell membrane (**Figure 4B**).

### EcTopoI and DNA gyrase have mutually exclusive localization on the *E. coli* chromosome

EcTopoI and DNA gyrase have opposite binding preferences and activities: while EcTopoI is attracted to and relaxes negative supercoils, DNA gyrase is attracted to and removes positive supercoils (Terekhova *et al*., 2012; Ashley *et al*., 2017; Liu *et al*., 2017; Sutormin *et al*., 2019). Comparison of ChIP-Seq data for EcTopoI and Topo-Seq data for DNA gyrase (Sutormin *et al*., 2019) directly demonstrates that *in vivo* gyrase enrichment is significantly lower in regions occupied by EcTopoI and *vice versa* (Welch t-test, p-value<1e-308, **Figure 4E**). While EcTopoI is enriched upstream of HETUs (where transcription-induced negative supercoiling should be high) and is depleted in downstream regions (where positive supercoiling should be accumulated) (**Figure 4D**), the gyrase enrichment pattern is the opposite. A signal of GapR-Seq, a method recently designed to detect positive supercoiling (Guo *et al*., 2021), matches the enrichment of DNA gyrase, indicating that gyrase accumulation in the downstream regions is indeed colocalized with increased positive supercoiling. While EcTopoI is particularly enriched in IRs flanked by divergent genes (see above), where cumulative negative supercoiling expected, the DNA gyrase signal is the highest for IRs between convergent genes (cumulative positive supercoiling expected) (**Figure 4C**). Finally, gyrase enrichment in downstream TU regions, as well as EcTopoI enrichment upstream positively correlates with transcription activity and is abolished by Rif (**Figure S8.I**). Together, these data indicate that EcTopoI and gyrase have opposing patterns of distribution genome-wide and these patterns are fully consistent with the predictions of the Liu & Wang model (Liu and Wang, 1987).

The DNA topoisomerases binding and cleavage sites may not fully overlap as was reported earlier for eukaryotic Top1 and prokaryotic *E. coli* TopoIV (El Sayyed *et al*., 2016; Baranello *et al*., 2017). To globally identify EcTopoI *in vivo* cleavage sites (TCSs), we constructed EcTopoI G116S M320V, an “intrinsically-poisoned” double mutant that forms stable covalent complexes with DNA (Cheng, Sorokin and Tse-Dinh, 2008). As expected, continuous production of EcTopoI G116S M320V from a plasmid led to growth inhibition (**Figure S9B**) and SOS-response (**Figure S9A**). EcTopoI G116S M320V was transiently (30 min) expressed in *E. coli* DY330 and trapped cleavage complexes were purified through a C-terminal strepII tag fused with the mutant topoisomerase (**Figure S9C**). At the time interval chosen, expression of EcTopoI G116S M320V had no apparent effect of cell culture growth (**Figure S9B**). Topoisomerase-associated DNA fragments were prepared for sequencing with Accel NGS 1S kit (Swift Biosciences), which allows strand-specific sequencing of single-stranded DNA, and the reads were mapped to the reference genome. Below, we refer to this experimental pipeline as “Topo-Seq”. The number of 3’-ends (N3E) was counted for every genomic position strand-specifically. Since EcTopoI forms a covalent intermediate with the 5’-end of a single-stranded break it introduces and leaves the 3’-end unmodified, an increase in the N3E should mark a TCS. A total of 262 TCSs were identified in the *E. coli* genome (125 on the forward and 137 on the reverse strand). The TCSs determined by Topo-Seq mark the sites of EcTopoI activity with single-base precision and significantly overlap with EcTopoI peaks detected with ChIP-Seq (Monte-Carlo simulation with 10000 iterations, p-value 3.5e-13, **Figure 5A**, **Figures S9D**, **E**). Interestingly, some regions characterized by increased EcTopoI binding (as evidenced by ChIP-Seq and ChIP-qPCR, **Figure S10D**) and cleavage (as evidenced by Top-Seq), demonstrated increased affinity to purified EcTopoI *in vitro*, revealing sequence specificity of the enzyme (**Figure 5B**, see also below).

**Figure 5.**
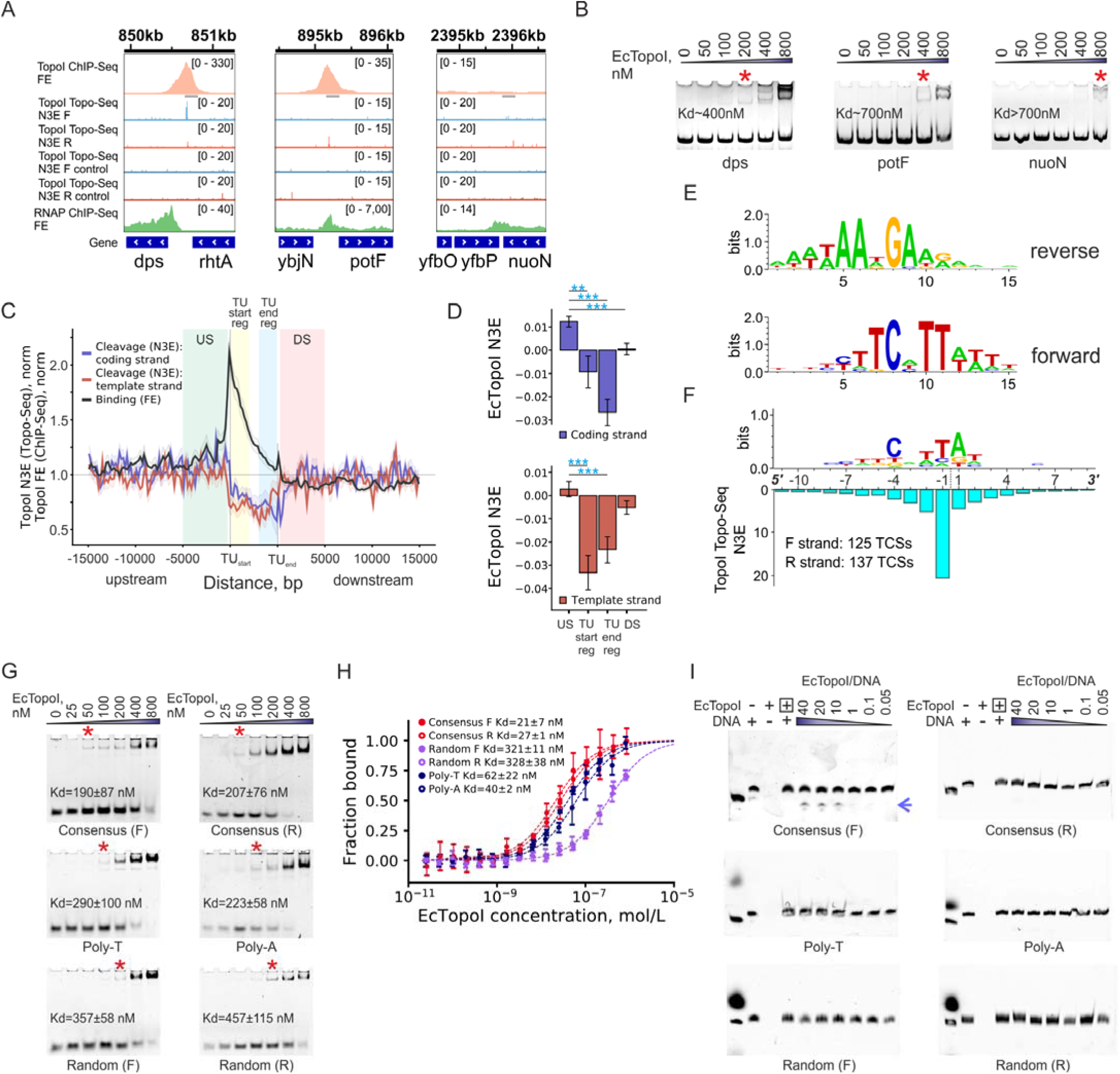
Topoisomerase I cleavage sites (TCSs) identified by Topo-Seq and sequence-specificity of binding and cleavage. (**A**) Representative regions of *E. coli* chromosome demonstrate the matching of EcTopoI peaks identified by ChIP-Seq and EcTopoI TCSs obtained by Topo-Seq (*dps*, *potF*) and a region lacking EcTopoI binding and activity (*nuoN*). EcTopoI cleavage activity is shown strand specifically for forward and reverse strands separately. As a control, a non-induced culture was used. Positions of amplified regions used for ChIP-qPCR and affinity measurements are indicated with grey rectangles. (**B**) Affinity of purified EcTopoI to three amplified regions of the genome indicated in panel **A** measured by EMSA: *nuoN* (control, low ChIP-Seq and Topo-Seq signals), *potF*, and *dps* (two highest peaks identified by ChIP-Seq and strong TCSs). DNA fragments (28.5 nM) were combined with EcTopoI (from 0 to 800 nM), reaction components were separated in a 10% native acrylamide gel and stained with EtBr. Red stars mark the lowest concentration of EcTopoI at which a gel-shift was detected. (**C**) Comparison of EcTopoI ChIP-Seq data (untreated condition, black curve) and EcTopoI Topo-Seq data. EcTopoI Topo-Seq signal is strand-specific and shown for coding (blue curve) and template (red curve) strands separately. Confidence bands around the mean metagene signal are represented by ±SEM. Metagene analysis was performed for the HETU set. Regions used for further quantification of enrichment in panel D (US, TU start region, TU end region, and DS) are shown by colored areas on the plot. (**D**) Comparison of EcTopoI cleavage signal in different regions relative to HETUs. Differences in the signal were tested using Welch t-test. Significance is indicated by asterisks. Error bars represent ±SEM. (**E**) A logo representing a binding motif of EcTopoI identified by ChIPMunk (Kulakovskiy *et al*., 2010) in sequences under the ChIP-Seq peaks. The motif is shown in both orientations. (**F**) EcTopoI cleavage motif identified by alignment of TCSs (cleavage is introduced between nucleotides at positions -1 and 1 and is indicated by a dashed line). The N3E signal is plotted below the motif. (**G**) EMSA experiments with EcTopoI and different Cy5-labled oligonucleotides. Binding of oligonucleotides in forward orientation is shown on the left, of complementary reverse oligonucleotides – on the right. Oligonucleotides in 20 nM concentration and a series of EcTopoI concentrations ranging from 0 to 800 nM were mixed for binding. Red stars mark the lowest concentration of EcTopoI at which a gel-shift was detected. (**H**) Results of MST experiments for Cy5-labled oligonucleotides and EcTopoI. Oligonucleotides in a concentration 0.5 nM and EcTopoI in concentrations ranging from 0 to 850 nM were mixed and MST was performed using Monolith NT.115. (**I**) DNA oligonucleotide cleavage by EcTopoI *in vitro*. A control with EcTopoI inactivated by high temperature is indicated by a boxed + sign. Cleavage products are marked with a blue arrow.

The binding and cleavage activities of EcTopoI were compared by metagene analysis. Both signals were increased upstream of active TUs, while the GapR-Seq signal was significantly decreased, indicating the relaxation of transcription-generated negative supercoiling by EcTopoI (**Figures 5C, D**). Interestingly, the cleavage activity of EcTopoI was significantly decreased at promoters and within the TUs bodies, i.e., at sites where formation of complexes with RNAP is expected. Indeed, the TCSs’ overlap with RNAP peaks is decreased (Monte-Carlo simulation with 10000 iterations, p-value 1.6e-2, **Figures S9D**, **F**). A similar pattern was reported for Top1 and RNAPII in eukaryotes (Baranello *et al*., 2017), implying that topoisomerase activity may be negatively regulated within the RNAP complexes both in pro- and eukaryotes.

### Identification of the EcTopoI binding and cleavage motif

We used ChIPMunk to find overrepresented motifs in EcTopoI ChIP-Seq peaks sequences. A strong motif was detected in more than 90% of enrichment peaks for all conditions tested (**Figure S10**). The motif was AT-rich, strongly asymmetric, and contained a conserved central TCNTTA/T part (**Figure 5E**). Single-stranded DNA oligonucleotides containing the consensus sequence were tested for their ability to bind EcTopoI *in vitro* by EMSA. Since the putative EcTopoI binding motifs is asymmetric, oligonucleotides corresponding to both strands of the consensus motif (“F”-forward, “R”- reverse) were used. Poly-T, Poly-A, and two complementary random-sequence oligonucleotides of equivalent length served as controls. As can be seen from **Figure 5G**, both consensus oligos bound EcTopoI with comparable affinities (K_D_ for “F” - 190±87 nM, for “R” - 207±76 nM); random oligos demonstrated the lowest affinities (K_D_ for “F” - 357±58 nM, for “R” - 457±115 nM), Poly-T and Poly-A had intermediate affinities with a higher affinity for Poly-A (K_D_ of 223±58 nM compared to 290±100 for Poly-T) (**Table S3**). In competition experiments, Consensus F oligo, when bound to EcTopoI, could not be displaced by 16-32 molar excess of random or Poly-T competitors. In contrast, Consensus F oligo readily displaced both random or Poly-T DNA from the EcTopoI complex at 0.125-4-fold molar excess. Consensus R oligo also readily displaced random oligos from a complex with EcTopoI, indicating stronger binding. Yet Consensus R oligo appeared to bind EcTopoI as efficiently as Poly-A DNA in competition experiments, which corresponds to K_D_ values reported above (**Figure S10E**).

We also used microscale thermophoresis (MST) to determine EcTopoI binding preferences. Again, the random sequence oligos had the lowest affinity (K_D_ for “F” - 321±11 nM, for “R” - 328±38 nM), while consensus oligo binding was the tightest (K_D_ for “F” - 21±7 nM, for “R” - 27±1 nM). The Poly-T and Poly-A oligos demonstrated intermediate affinities with a higher affinity for Poly-A (a K_D_ 40±2 nM versus 62±22 nM for Poly-T) (**Figure 5H** and **Table S3**). Overall, the binding affinities for EcTopoI revealed in our experiments rank as Consensus F>Consensus R∼Poly-A>Poly-T>>random oligos.

To determine whether there is a specific EcTopoI cleavage motif, we aligned sequences around the TCSs. The revealed cleavage motif was highly similar to the binding motif identified using ChIP-Seq. As can be seen from **Figure 5F**, EcTopoI cleaves a TA dinucleotide located 4 nucleotides downstream of a conserved C residue. The cleavage motif was validated *in vitro*: cleavage was only observed for Consensus F oligo, bearing a sequence that completely matches the deduced cleavage motif (**Figure 5I**). Overall, we conclude that while EcTopoI prefers to bind to AT-rich single-stranded sequences, a single C residue within an AT-rich patch is needed for efficient cleavage. Earlier *in vitro* experiments, demonstrated that type-IA topoisomerases, including EcTopoI, specifically recognize a C and cleave DNA 4 nucleotides downstream (Y. Tse, Kirkegaard and Wang, 1980; Zhang, Cheng and Tse-Dinh, 2011; Narula and Tse-Dinh, 2012). Our results extend these observations and show that a C in a specific context is required for EcTopoI cleavage *in vivo*.

### Impairing the RNAP:EcTopoI interaction mimics inactivation of EcTopoI and is deleterious for cell growth

If the RNAP:EcTopoI complex has a physiological role, uncoupling of the RNAP-EcTopoI interaction shall have an impact on cell viability. Indeed, overnight overexpression of the 14 kDa EcTopoI CTD dramatically inhibited colony formation, while 1-hour overexpression had a bacteriostatic effect and led to ∼2-fold decrease in the number of CFUs (**Figures S5A** and **B**). Consistently, overexpression of CTD slowed culture growth ∼75 min post-induction (**Figure 6Aiii**). Overexpression of GFP or full-length EcTopoI had no such effect (**Figures 6Ai, ii**). Cells expressing the CTD formed filaments and underwent SOS response, indicating accumulation of DNA breaks (**Figure 6B, Figure S9A**).

**Figure 6.**
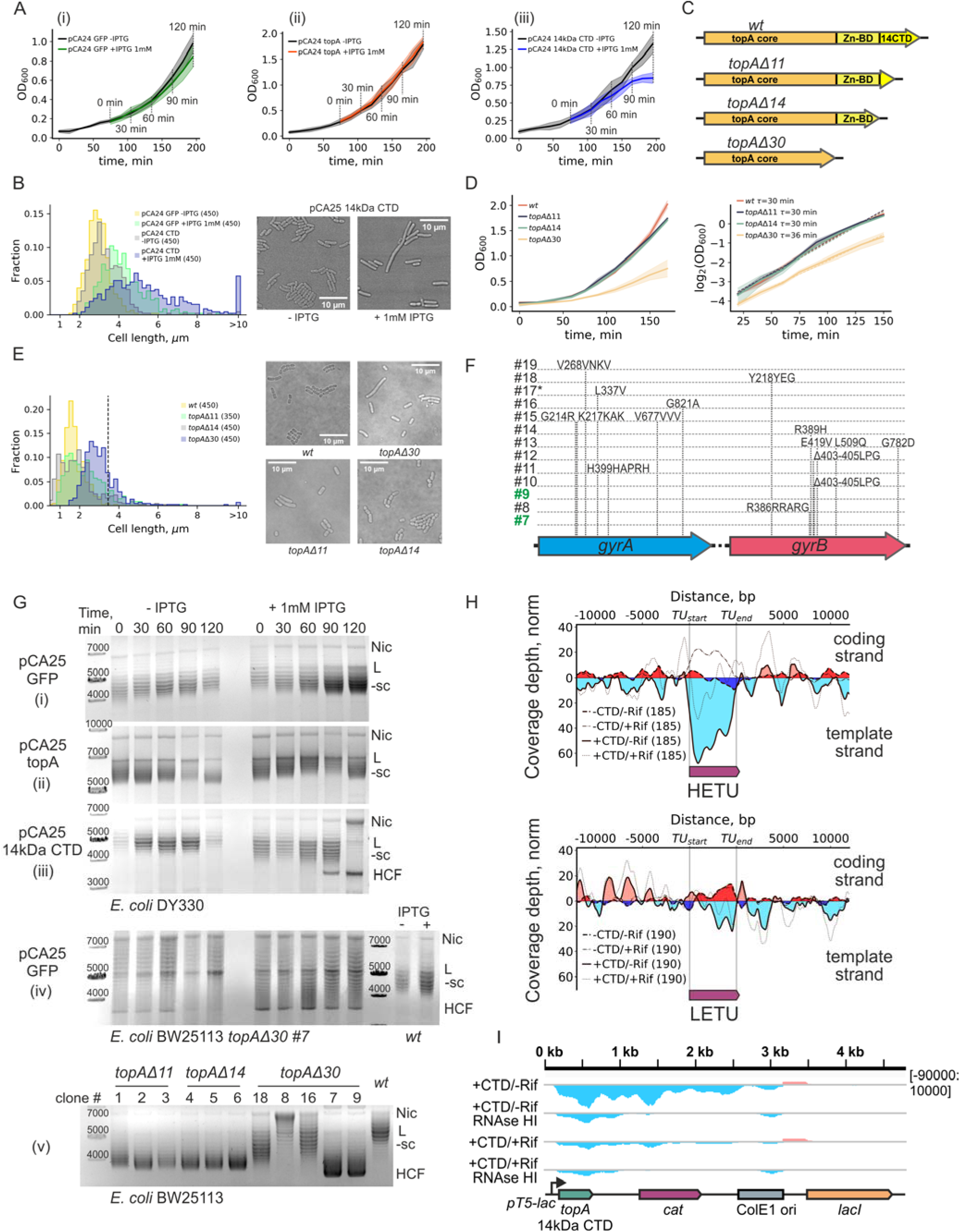
Uncoupling of RNAP:EcTopoI complex is toxic for cells, leads to hypernegative supercoiling of plasmids, and accumulation of R-loops. (**A**) Growth curves of *E. coli* DY330 *topA*-SPA harboring pCA24 GFP (i), pCA24 14kDa CTD (ii) or pCA24 topA (iii) plasmids in LB supplemented with 0.5% glucose. At OD_600_∼0.2 1 mM IPTG was added to indicated cultures (“+ IPTG 1 mM”, means of three biological replicates are shown), non-induced cultures served as controls (“- IPTG”, means of three biological replicates shown). Shade represents 0.95 confidential interval of the mean. Grey lines mark time-points when aliquots were taken for plasmid extraction. (**B**) Quantification of cell length in CTD or GFP (control) producing cultures. Distributions of cell lengths are shown on the left. Cells longer than 10 µm are collected into an overflow bin. Numbers of cells are indicated in brackets. Representative fields are shown on the right. *E. coli* DY330 *topA-SPA* cells harboring indicated plasmids were grown and induced as for ChIP-Seq experiments and examined by bright-field microscopy. (**C**) Graphical representation of truncated versions of a *topA* gene constructed by recombineering in *E. coli* BW25113. (**D**) Growth curves of *E. coli* BW25113 strains with truncated versions of *topA* and of wild-type control (on the left). Shade represents 0.95 confidential interval of the mean. On the right - quantification of doubling time for exponential regions of growth curves. (**E**) Quantification of cell length in cultures of *E. coli* BW25113 strains with truncated versions of *topA* and the wild-type. Distributions of cell lengths is shown on the left; dashed vertical line marks 2*mean cell length for wild-type. Representative fields are shown on the right. The strains were grown in LB until mid-exponential phase (OD_600_∼0.6) and examined by bright-field microscopy. (**F**) Mutations in gyrase genes *gyrA* and *gyrB* found in different clones of *E. coli* BW25113 *topA* Δ *30*. A star for clone #17 indicates amplification of a chromosomal region containing TopoIV topoisomerase genes *parC* and *parE*; clones ##7 and 9, lacking compensatory mutations, are indicated in green. (**G**) Supercoiling of pCA24 GFP (i), pCA24 topA (ii), and pCA24 14 kDa CTD (iii) plasmids extracted from exponentially growing *E. coli* DY330 *topA*-SPA cultures. Supercoiling of pCA24 GFP (iv) plasmid extracted from exponentially growing *E. coli* BW25113 *topA* Δ30 culture (time-course, on the left) or *E. coli* BW25113 *wt* (two rightmost lanes). At OD_600_∼0.2 cultures were either induced with IPTG (“+1 mM IPTG”) or continued growth without induction (“- IPTG”). Cultures were sampled every 30 min, corresponding to time-points indicated in panel **A**. (v) Supercoiling level of the pCA24 GFP plasmid extracted from overnight cultures of different clones of *E. coli* BW25113 *topA* mutants and from the wild-type control. Clone numbers match those in panel **F**. Nic – nicked plasmid, L – linear plasmid, -sc – negatively supercoiled plasmid, HCF – hypercompacted form of a plasmid. Plasmid forms were separated by electrophoresis in 1% agarose gels in TAE buffer supplemented with 5 µg/mL chloroquine and visualized by EtBr staining. (**H**) Metagene analysis of normalized strand-specific read coverage depth obtained in DRIP-Seq experiments with exponentially growing *E. coli* DY330 *topA*-SPA cultures grown at conditions identical to those used for ChIP-Seq with EcTopoI. Coverage depth for -CTD/-Rif experiment is shown with a dashed line, enrichment on the “+” and “-” strands is shown by dark-red and dark-blue fillings respectively. Coverage depth for +CTD/-Rif experiment is shown with a solid line, enrichment on the “+” and “-” strands is shown by light-red and light-blue fillings respectively. Right plot represents coverage for a set of highly-transcribed transcription units (HETU, the rRNA operons were excluded), left – for low-transcribed units (LETU). Numbers of transcription units used in metagene analysis are indicated in parenthesis. Schematic transcription units are shown in purple below. (**I**) Normalized strand-specific read coverage depth of the pCA24 14 kDa CTD expression plasmid obtained in DRIP-Seq experiments with exponentially growing *E. coli* DY330 *topA*-SPA cultures. Cultures were either treated with RNAP inhibitor rifampicin or 10 minutes before cells harvesting (+Rif) or remained untreated (-Rif). For a control, coverage for samples treated with RNAse HI before immunoprecipitation with the S9.6 antibodies is shown (RNAse HI tracks). Coverage for “-” and “+” strands is shown in light-blue and light-red, respectively. A linearized map of the pCA24 14kDa CTD plasmid is shown below.

As an alternative strategy to disrupt the RNAP-EcTopoI interaction, we constructed *E. coli* strains with *topA*Δ*11* (recapitulates the well-known *topA66* mutation (Biekt and Cohen, 1989)), *topA*Δ*14*, and *topA*Δ*30* mutations. These mutations lead to production of EcTopoI lacking, correspondingly, an 11 kDa portion of CTD, the entire 14 kDa CTD, or a longer 30 kDa fragment that includes both the CTD and the Zn-binding domain (**Figure 6C**). Since EcTopoI interacts with RNAP through both the CTD and the Zn-binding domain (Cheng *et al*., 2003), we expected that *topA* Δ *11* and *topA* Δ *14* will decrease the RNAP interaction, while *topA* Δ *30* will abolish it. In fact, the *topA*Δ*30* deletion shall inactivate EcTopoI (Zumstein and Wang, 1986; Ahumada and Tse-Dinh, 2002). While *topA* Δ *11* and *topA* Δ *14* strains formed colonies that were indistinguishable from wild-type, colonies formed by the *topA*Δ*30* mutant were heterogeneous: most were much smaller than wild-type others had a wild-type appearance. Whole-genome sequencing revealed that cells from *topA* Δ *11* and *topA* Δ *14* colonies had no additional mutations, while cells from nearly all fast-growing *topA* Δ *30* colonies harbored mutations in the gyrase genes (**Figure 6F**). Amplification of the chromosomal region containing the *parC* and *parE* genes encoding topoisomerase TopoIV was detected in one of the *topA* Δ *30* clones (#17). Two *topA* Δ *30* clones (##7 and 9) had no additional mutations. Clone #7 was used for further analysis. The growth curve analysis showed that while the doubling time of *topA* Δ *11* and *topA* Δ *14* strains was indistinguishable from that of the wild-type (30 min), the *topA*Δ*30* clone grew slower, with a doubling time of 36 min (**Figure 6D**). The fraction of long (>2* mean length of wild-type cells) cells was considerably higher in *topA* Δ *11* and *topA* Δ *14* cultures (11%) compared to the wild-type (0.3%). The fraction of long cells was 23% in *topA* Δ *30* culture, and these cells were much longer than the wild-type or other mutant cells (**Figure 6E**). Consistent with increased frequency of longer cells, the *topA* Δ *11* and *topA* Δ *14* mutants were outperformed by the wild-type in long-term competition experiments (**Figure S11**). Overall, we conclude that uncoupling of the RNAP:EcTopoI complex formation by CTD overexpression phenotypically mimics inactivation of EcTopoI in *topA* Δ *30* clones. Suppressor mutations in *topA* Δ *30* clones might reduce global negative supercoiling by the gyrase, thus compensating the deleterious effects of EcTopoI inactivation.

### Impairing of the RNAP:EcTopoI interaction leads to excessive negative supercoiling and accumulation of R-loops

It has been suggested that the interaction of EcTopoI with RNAP may prevent the formation of R-loops behind the elongating transcription complex, thus helping restore the DNA duplex and increasing the processivity of transcription (Yang *et al*., 2015). To test this idea, we examined changes in the topological state of plasmids in cells overexpressing the 14 kDa CTD, a condition that impairs the RNAP:EcTopoI interaction (above). If uncoupling leads to accumulation of R-loops, hypernegative supercoiling of plasmid DNA shall be expected (Masse and Drolet, 1999). Indeed, in agreement with earlier reported data of Cheng et al. (Cheng *et al*., 2003), negative supercoiling of plasmids increased dramatically in cells overexpressing the CTD. In fact, over time, a hypercompacted plasmid form appeared in these cells. This form migrated faster than any other topoisomer (**Figure 6Giii**) and may have corresponded to plasmids containing R-loops (Masse and Drolet, 1999). No excessive supercoiling or plasmid hypercompaction was observed upon overexpression of GFP or full-length EcTopoI (**Figures 6Gi, ii**, the latter condition led to plasmid relaxation, as expected).

Another condition that affects RNAP:EcTopoI interaction is deletion of the C-terminal region of EcTopoI (above). To access plasmid supercoiling level in BW25113 *topA* mutants and the wild-type strain, plasmids were extracted from overnight cultures and topoisomers were resolved by electrophoresis. Plasmids purified from *topA* Δ *11* and *topA* Δ*14* clones, where the strength of RNAP:EcTopoI interaction is reduced, had higher levels of negative supercoiling than plasmids purified from the wild-type control. Supercoiling levels varied dramatically for plasmids purified from different *topA*Δ*30* clones, where EcTopoI is inactivated and there is no interaction with RNAP. The highest level of negative supercoiling, approaching that of hypernegatively supercoiled plasmid from CTD-expressing cells, was observed for plasmids purified from clones ##7 and 9 that lacked suppressing mutations (**Figure 6Gv**). Plasmid hypercompaction in the *topA*Δ*30 clone #7* was more prominent when expression of plasmid-borne *gfp* gene was induced with IPTG, indicating that both uncoupling of RNAP:EcTopoI complex/EcTopoI inactivation *and* active transcription contribute to this phenomenon, consistent with the R-loops accumulation hypothesis (**Figure 6Giv**).

To directly observe accumulation of R-loops, we performed strand-specific DRIP-Seq at conditions identical to those used for ChIP-Seq. R-loops accumulation in HETUs was revealed upon overexpression of the 14 kDa CTD (**Figure 6H**). In addition, a large portion of the pCA24 14 kDa CTD expression plasmid, including the *topA* fragment encoding the 14 kDa CTD and the *cat* antibiotic resistance gene transcribed in the same direction, was covered by R-loops upon induction of transcription. Rifampicin abolished R-loops accumulation (**Figure 6I**). Dot-blot analysis also demonstrated increased level of R-loops in response to CTD overexpression (**Figure S5G**) (see **Supplementary Materials & Methods** for procedure details). We conclude that uncoupling of RNAP:EcTopoI leads to accumulation of transcription-induced R-loops, which may be the basis of toxicity of overexpressed 14 kDa CTD.

## Discussion

In this study, several important observations are made. First, we show interaction between TopoI and RNAP is required for *E. coli* cell viability. Disruption of the interaction leads to hypernegative DNA supercoiling and to dramatic R-loops accumulation. Our data provides a mechanistic explanation for TopoI –RNAP complex function and show that TopoI is required for R-loops formation control. Second, we demonstrate directly that TopoI and DNA gyrase are localized in extended upstream and downstream regions of TUs, illustrating the diffusion of unconstrained supercoils generated by transcription in accordance with the twin-domain model. Finally, we revealed that both DNA topology and local sequence patterns define localization and activity of TopoI genome-wide.

*In vitro*, bacterial topoisomerase I efficiently relaxes negatively supercoiled DNA (Terekhova *et al*., 2012; Liu *et al*., 2017). Therefore*, in vivo* the enzyme should be able to relieve negative supercoiling generated by transcription and, possibly, replication, and compensate the DNA gyrase activity to maintain physiological level of supercoiling (Massé and Drolet, 1999; Zechiedrich *et al*., 2000). *E. coli* topoisomerase I was demonstrated to interact through its Zn-binding and C-terminal domains with the β’ subunit of the RNA polymerase (Cheng *et al*., 2003; Tiwari *et al*., 2017). Eukaryotic TOP1 and RNAPII, as well as mycobacterial and streptococcal TopoI and RNAP were also shown to interact. These interactions must have evolved independently, since TOPI and prokaryotic TopoI are evolutionarily distant from each other (McKie, Neuman and Maxwell, 2021), while MtbTopoI appears to interact with MtbRNAP through a domain other than the one used on the case of the *E. coli* enzyme (Gupta *et al*., 2006). Yet, the ubiquitous presence of this interaction suggests the existence of common topological problems associated with transcription that need to be solved (Banda, Cao and Tse-Dinha, 2017; Baranello *et al*., 2017).

### Genome-wide localization patterns of EcTopoI and DNA gyrase supports the twin-domain model

Considering the fact that EcTopoI interacts with RNAP and has an increased affinity to negatively supercoiled DNA, it should be particularly enriched at highly-transcribed regions of the bacterial chromosome. In full agreement with this expectation, we observed that the ChIP-Seq signal from EcTopoI is enriched in the bodies and promoters of active transcription units. We also detected prominent EcTopoI enrichment in extended, up to ∼12-15 Kbs, regions upstream of active TUs. We propose that this enrichment defines the range that transcription-induced supercoiling can diffuse along the *E. coli* chromosome. Analysis of EcTopoI enrichment in intergenic regions supports that the enzyme is attracted to topologically constrained regions that accumulate negative supercoiling. An inverted enrichment pattern is observed for DNA gyrase and GapR, which are known to act upon/interact with positively supercoiled DNA. These observations provide, for the first time, a whole-genome validation of the twin-domain model proposed by Liu & Wang (Liu and Wang, 1987).

### EcTopoI interacts with RNAP *in vivo*

The EcTopoI signal in transcription unit bodies overlaps with the RNAP signal and is abolished by rifampicin. In contrast, EcTopoI enrichment, as well as RNAP enrichment at promoters is unaffected by Rif. These data further support a tight linkage of EcTopoI with RNAP. Furthermore, over-expression of EcTopoI CTD, which squelches EcTopoI-RNAP interaction, decreases the enrichment of topoisomerase in transcription units and promoters, while leaving signals in extended upstream regions unaffected. Since CTD does not influence transcription elongation, the results support the existence of an RNAP:EcTopoI complex that is formed during transcription initiation and persists during transcription elongation *in vivo*. In contrast to *E. coli*, enrichment of streptococcal SpTopoI is abolished by Rif, possibly indicating a distinct mechanism of RNAP:TopoI interaction (Ferrandiz, Hernandez and de la Campa, 2021).

### *E. coli* and *Mycobacterium* rely on different versions of the twin-domain model

Genome-wide binding of mycobacterial topoisomerases (DNA gyrase and TopoI) and RNAP was studied by Nagaraja group, who showed colocalization of the topoisomerases and RNAP (Ahmed *et al*., 2017; Rani and Nagaraja, 2018), see also re-analysis of their ChIP-Seq data in **Figures S3** and **S4E-F**). Surprisingly, in *Mycobacterium*, both topoisomerases are enriched within the bodies of transcription units with the highest signal near the transcription start sites. In contrast to *E. coli,* there is no evidence for supercoiling of any sign diffusing away from the TUs (**Figures 7A, B**). These patterns define two possible variations of the original Liu & Wang scheme: an “open” one for *E. coli* (and, likely, *S. pneumoniae*, **Figure S3R**) in which supercoiling domains extend over a substantial distance from moving RNAP complexes and a “closed” one for *Mycobacterium*, where the domains are trapped within polymerase-topoisomerases complex without escape of supercoils. Possibly, mycobacterial topoisomerases in complex with RNAP are able to fully relax supercoils generated in TUs and, therefore, their activity is not needed in adjacent regions. Hypothetical “semi-open” sub-models may also exist: i) when gyrase is highly-active in complex with RNAP and TopoI does not interact with RNAP, allowing negative supercoiling to diffuse freely upstream of TUs; ii) an opposite situation, when only TopoI is active within the complex with RNAP, with only positive supercoiling diffusing downstream of TUs where it is relaxed by RNAP-free gyrase (**Figure 7C**). Additional variations may be possible depending on the balance of topoisomerase and RNAP activities. If topoisomerase activity exceeds the rate at which RNAPs generate supercoiling, diffusion of supercoiling will be constrained by rapid relaxation by topoisomerases. Based on this logic, EcTopoI in complex with its cognate RNAP allows a portion of unconstrained supercoiling to diffuse upstream, where it is relaxed by RNAP-free topoisomerase.

**Figure 7.**
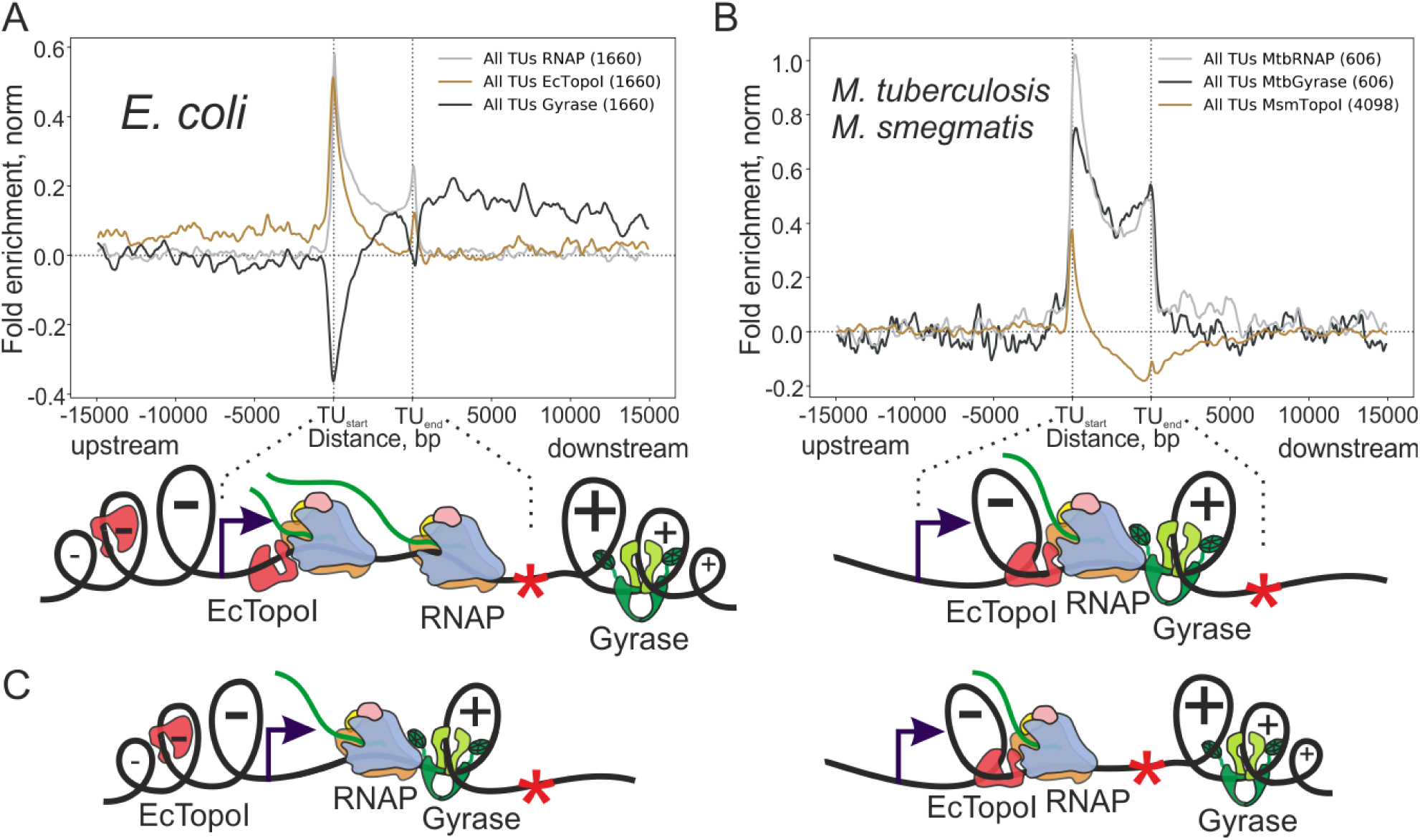
Possible variations of the twin-domain model. Average normalized enrichment of TopoI, DNA-gyrase, and RNAP over transcription units of *E. coli* (**A**, “open” model) and *Mycobacterium* (**B**, “closed” model). Graphical representation of twin-domain sub-models is shown below. ChIP-Seq data for *M. tuberculosis* MtbRNAP, MtbGyrase, and *M. smegmatis* MsmTopoI was taken from publicly available datasets (Uplekar *et al*., 2013; Ahmed *et al*., 2017; Rani and Nagaraja, 2018). (**C**) Other “semi-open” hypothetical variations of the twin-domain model, based on interaction of key topoisomerases (TopoI, DNA-gyrase) with RNAP and their activity within a complex.

### EcTopoI cleavage activity might be negatively regulated in complex with RNAP

By using a transient expression of an “intrinsically-poisoned” topoisomerase mutant we performed Topo-Seq, an approach which allowed us to identify EcTopoI cleavage sites genome-wide. A similar approach was earlier applied for TopoI from *M. smegmatis* (MsmTopoI) (Rani and Nagaraja, 2018). Interestingly, despite the prominent binding of both EcTopoI and MsmTopoI to promoter regions, neither enzyme cleaves DNA there. However, the pattern of activity in TU bodies and in regions upstream of TUs is different for the two enzymes. First, MsmTopoI was shown to be active (i.e., cleaves DNA) in TUs bodies. In contrast, EcTopoI is inactive in the beginning of TUs but its activity is rising toward the end of TUs, particularly in LETUs. This can reflect activation of EcTopoI by extensive torsional stress generated by individual RNAP molecules expected in LETUs. Conversely, in HETUs, convoys of RNAP molecules should mutually annihilate positive and negative supercoils (Kim *et al*., 2019; Alena and Kolomeisky, 2021). Second, no activity of MsmTopoI was detected in extended regions upstream of TUs, while EcTopoI activity in these regions is significantly increased, again illustrating the proposed “open” and “closed” variants of the twin-domain model. Overall, the data for EcTopoI resemble the activity pattern of eukaryotic Top1, which is bound to RNAPII but is inactive until a signal is triggered by a RNAPII stalled by a torsional stress (Baranello *et al*., 2017). A similar, independently evolved mechanism could be operational for bacterial topoisomerases. A naturally occurring modification of EcTopoI, N^ε^-acetylation of lysins, was reported to reduce activity of the enzyme *in vivo* (Zhou *et al*., 2017). We speculate that this modification can be involved in the regulation of EcTopoI activity in its complex with RNAP. We predict that EcTopoI activity is inhibited by acetylation at promoters and in the beginning of TU bodies and is activated by de-acetylation (probably, by the CobB protein) in the ends of TUs. This de-acetylation may be triggered by conformational changes within the EcTopoI-RNAP complex caused by RNAP stalling or by extensive torsional stress.

Topoisomerase I can catenate and decatenate single-stranded DNA circles and double-stranded circles containing nicks *in vitro* (Tse and Wang, 1980; Bhaduri *et al*., 1998; Rani and Nagaraja, 2018). Recently, enrichment of TopoI activity was found at the Ter region of *M. smegmatis* chromosome (Rani and Nagaraja, 2018). This bacterium is lacking classical topoisomerases-decatenases, TopoIII and TopoIV, and, thus, its TopoI is likely to be involved in chromosomes decatenation. Despite similarity in *in vitro* activities, we did not observe any specific binding or cleavage by EcTopoI near Ter regions of the *E. coli* chromosome, indicating that this enzyme is not involved in chromosome decatenation.

### EcTopoI has sequence specificity *in vivo*

By combining ChIP-Seq and Topo-Seq we identified the *in vivo* binding/cleavage motif of EcTopoI, which was validated *in vitro*. The EcTopoI binding motif is asymmetric, AT-rich, and contains a single conserved C residue. This C is located 4 nucleotides upstream of the cleavage site, which occurs at a TA dinucleotide. Our data indicate that AT-rich sequences are good binding substrates for EcTopoI but appropriately positioned C residue is strictly required for cleavage, which agrees with previous *in vitro* observations made for different type IA topoisomerases (Y.-C. Tse, Kirkegaard and Wang, 1980; Kovalsky, Kozyavkin and Slesarev, 1990; Annamalai *et al*., 2009; Zhang, Cheng and Tse-Dinh, 2011; Narula and Tse-Dinh, 2012). Likely, this requirement is characteristic for the entire protein family. For optimal activity of EcTopoI, the binding/cleavage motif should be “activated” by melting of DNA, excessive negative supercoiling upstream of TUs, or, perhaps, by hypothetical signaling such as de-acetylation of EcTopoI when in complex with RNAP at the ends of TUs.

### RNAP:EcTopoI complex is required for R-loops formation control

Overexpression of EcTopoI CTD is toxic for *E. coli* (Cheng *et al*., 2003). We demonstrate that cells overexpressing the CTD form filaments and accumulate R-loops genome-wide. In addition, plasmids purified from these cells are hypernegatively supercoiled. Earlier, such changes were detected in *topA-null* mutants (Pruss, 1985; Masse, Phoenix and Drolet, 1997; Baaklini *et al*., 2008; Usongo *et al*., 2013) as also observed in this work for the *topA*Δ*30* mutant, which lacks the relaxation activity (Zumstein and Wang, 1986; Lima, Wang and Mondragón, 1993; Ahumada and Tse-Dinh, 2002). It was proposed, that the primary role of topoisomerase I is to prevent extensive R-loops formation by relaxation of excessive negative supercoiling (Masse and Drolet, 1999; Brochu *et al*., 2018). Since EcTopoI remains fully active upon CTD overexpression, as indicated by enrichment in upstream regions of active transcription units, it appears that physical interaction between RNAP and EcTopoI, which is affected by CTD, is required to efficiently prevent R-loops formation during transcription.

Why then *E. coli topA*Δ*11* and *topA*Δ*14* mutants have close to wild-type viability (despite slightly increased cell length, more negatively supercoiled DNA, and decreased fitness in competition experiments)? We suggest that the Zn-binding domain of EcTopoI is primarily responsible for the binding to RNAP and/or its affinity to RNAP is sufficient for required levels of complex formation. Of note, molecular dynamics simulation predicted residues of Zn-binding domain, but not of CTD, to be involved in the interaction with RNAP (Tiwari *et al*., 2017). We assume that when CTD is overexpressed and binds to RNAP, it sterically excludes topoisomerase from the complex.

We did not observe signs of mechanisms compensating the uncoupling of the RNAP:EcTopoI complex. Rather, cells rapidly acquired mutations in the gyrase genes and/or amplified genes of TopoIV (Dinardo *et al*., 1982; Pruss *et al*., 1982; Brochu *et al*., 2018). Therefore, the complex EcTopoI-RNAP interface may be a promising target for a new class of antibacterials. In line with this conjecture, it was demonstrated that the *topA66* mutation in *E. coli* led to decreased SOS-response and increased sensitivity to antibiotics (Yang *et al*., 2015). Similarly, overexpression of *M. tuberculosis* TopoI CTD resulted in increased susceptibility to antibiotics and oxidative stress (Banda, Cao and Tse-Dinha, 2017). We expect that uncoupling of the TopoI-RNAP complex by a drug can lead to fixation of mutations in gyrase genes, decreasing the gyrase activity. If so, administration of such a hypothetical drug followed by a course of gyrase-targeting inhibitor would be beneficial, as gyrase inhibitor will act on an essential enzyme whose activity is already decreased by mutations, leaving less space on the mutational landscape for accumulation of additional mutations conferring resistance to antibiotics.

## Data and software availability

Sequencing data for *E. coli* RNA-Seq (GSE181687), *E. coli* TopoI ChIP-Seq and Topo-Seq (GSE181915 and GSE182473, respectively), *E. coli* RpoC ChIP-Seq (GSE182850), *E. coli* DRIP-Seq (GSE181945) were deposited in GEO with corresponding dataset accession numbers. Sequencing data for *E. coli topA* mutants’ whole genome sequencing was deposited in SRA (PRJNA757761). See full list of datasets used in the study in Table S2. Custom scripts used for data analysis are available from Github repository https://github.com/sutormin94.

## Funding & Acknowledgements

This work (bioinformatic analysis, microscopy, and micro-scale thermophoresis) was supported by grant 075-15-2019-1661 from the Ministry of Science and Higher Education of the Russian Federation. This work was also supported by Skoltech NGP Program (Skoltech-MIT joint project) and RFBR grant, project number 20-34-90069. Sequencing was supported by the Skoltech Life Sciences Program grant. Sequencing was performed by Skoltech Genomics Core Facility. We acknowledge Dr. Marina Serebryakova for mass spectrometry and Dr. Svetlana Dubiley for extensive and fruitful project discussions.

## Author contributions

**D.S.** conceived the study and designed experiments. **A.G.** performed EcTopoI ChIP-Seq and DRIP-Seq experiments. **D.S.** conducted Topo-Seq experiments. **K.O.** and **S.B.** performed RpoC ChIP-Seq experiments. **D.S.** obtained *E. coli* BW25113 *topA* derivatives. **O.M.** sequenced *E. coli* BW25113 strains with edited *topA* gene. **D.S.** performed all NGS data analysis. **A.G.** conducted DNA cleavage and DNA binding experiments. **D.S.** analyzed topology of plasmids. **A.R.** performed microscopy. **D.T.** conducted pull-down experiments, purified EcTopoI and EcTopoI CTD. **D.S.** prepared all figures. **D.S.**, **S.B.**, and **K.S.** wrote the manuscript, which was read, edited, and approved by all authors.

## Competing interests

The authors declare no competing interests.

**Supplementary Information** is available for this paper.

**Correspondence and requests** for materials should be addressed to Dmitry Sutormin or Konstantin Severinov.

## List of acronyms

EcTopoI: *E. coli* topoisomerase I
MsmTopoI: *M. smegmatis* topoisomerase I
MtbTopoI: *M. tuberculosis* topoisomerase I
MtbRNAP: *M. tuberculosis* RNA polymerase
MtbGyrase: *M. tuberculosis* DNA-gyrase
SpTopoI: *S. pneumoniae* topoisomerase I
CTD: C-terminal domain of topoisomerase I
RNAP: RNA polymerase
Rif: RNAP inhibitor rifampicin
Kb, Mb: kilo- and megabase pairs respectively
TU: transcription unit
LETU: transcription units with low level of expression: 200 Tus
HETU: transcription units with high level of expression: 200 Tus
TSS: transcription start site
IR: intergenic region
TF: transcription factor
EMSA: electrophoretic mobility shift assay
MST: microscale thermophoresis
FE: fold enrichment

