## Supplementary Figures for "Interaction Between Transcribing RNA Polymerase and Topoisomerase I Prevents R-loop Formation in *E. coli*"

**Figure S1** related to Figures 2, 3. Correspondence among biological replicas of EcTopoI ChIP-Seq experiments. (**A**) Heatmaps represent pair-wise Pearson correlation between EcTopoI ChIP-Seq datasets (fold enrichment). From left to right: correlations between CTD-Rif- (no EcTopoI 14kDa CTD overexpression, no treatment with Rif) samples; correlations between CTD-Rif+ (no EcTopoI 14kDa CTD overexpression, treatment with Rif) samples; correlations between CTD+Rif- (EcTopoI 14kDa CTD overexpression, treatment with Rif) samples; correlation between CTD+Rif+ (EcTopoI 14kDa CTD overexpression, treatment with Rif) samples. (**B**) Heatmaps represent pair-wise number of enrichment peaks identified with MACS2 shared between EcTopoI ChIP-Seq datasets**.** From left to right: numbers of peaks shared between CTD-/Rif- biological replicas; numbers of peaks shared between CTD-/Rif+ biological replicas; numbers of peaks shared between CTD+/Rif- biological replicas; numbers of peaks shared between CTD+/Rif+ biological replicas. (**C**) Numbers of EcTopoI peaks shared between different experimental conditions. Peaks considered are presented in all biological replicas of particular experimental condition. (**D**) Presented matrices of shared EcTopoI peaks are not symmetric, because several enriched regions (peaks) of sample 1 can overlap with one long peak of sample 1. In this example, number of shared peaks will be higher for sample 1 than for sample 2.

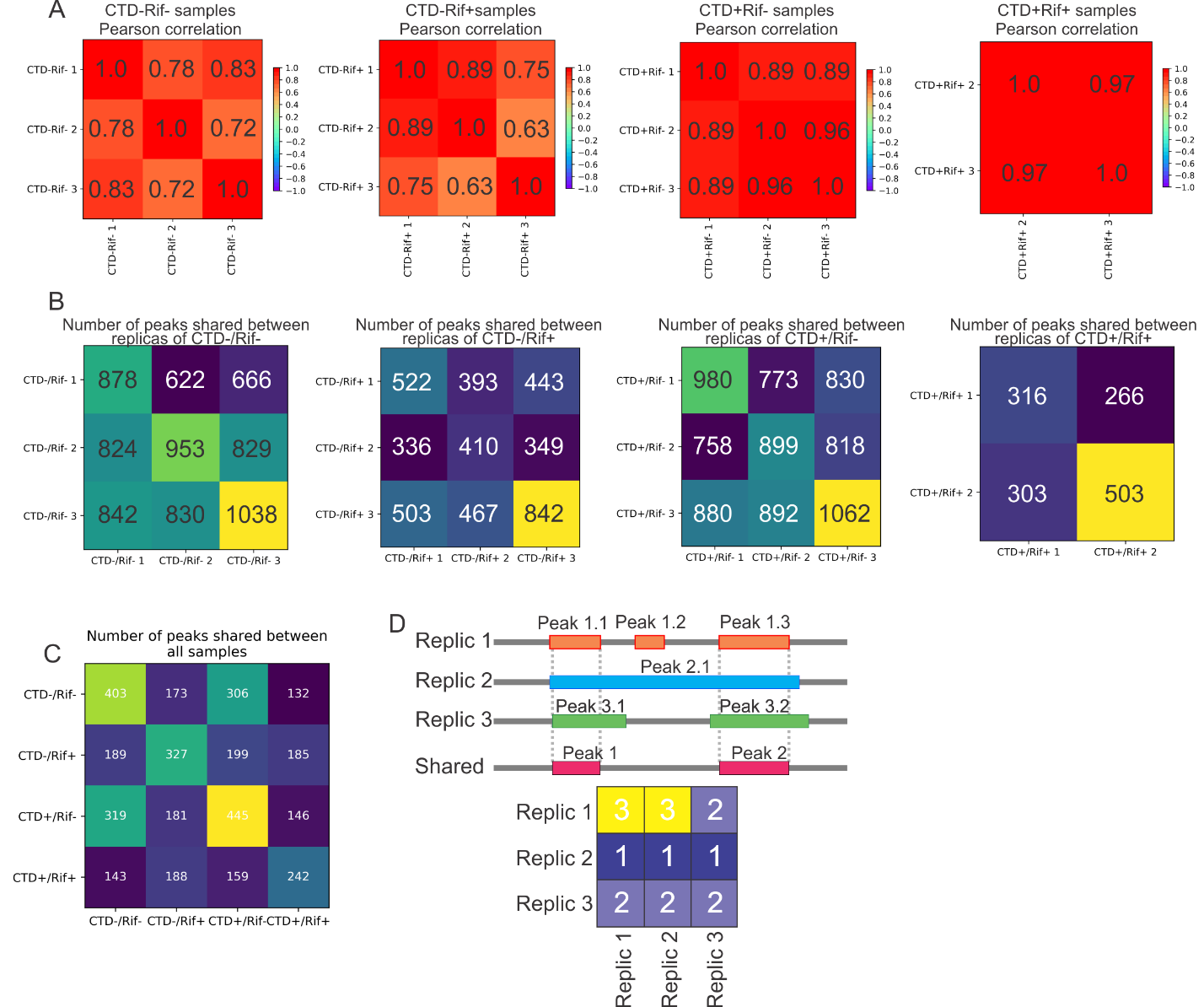

**Figure S2** related to Figure 3**.** Description of peaks called with MACS2 in EcTopoI ChIP-Seq datasets. **A**. CTD-/Rif- peaks. **B**. CTD-/Rif+ peaks. **C**. CTD+/Rif- peaks. **D.** CTD+/Rif+ peaks. **E.** Peaks shared among all datasets. **F.** Fold enrichment of peaks shared (**S – for shared**) among all datasets for different datasets.

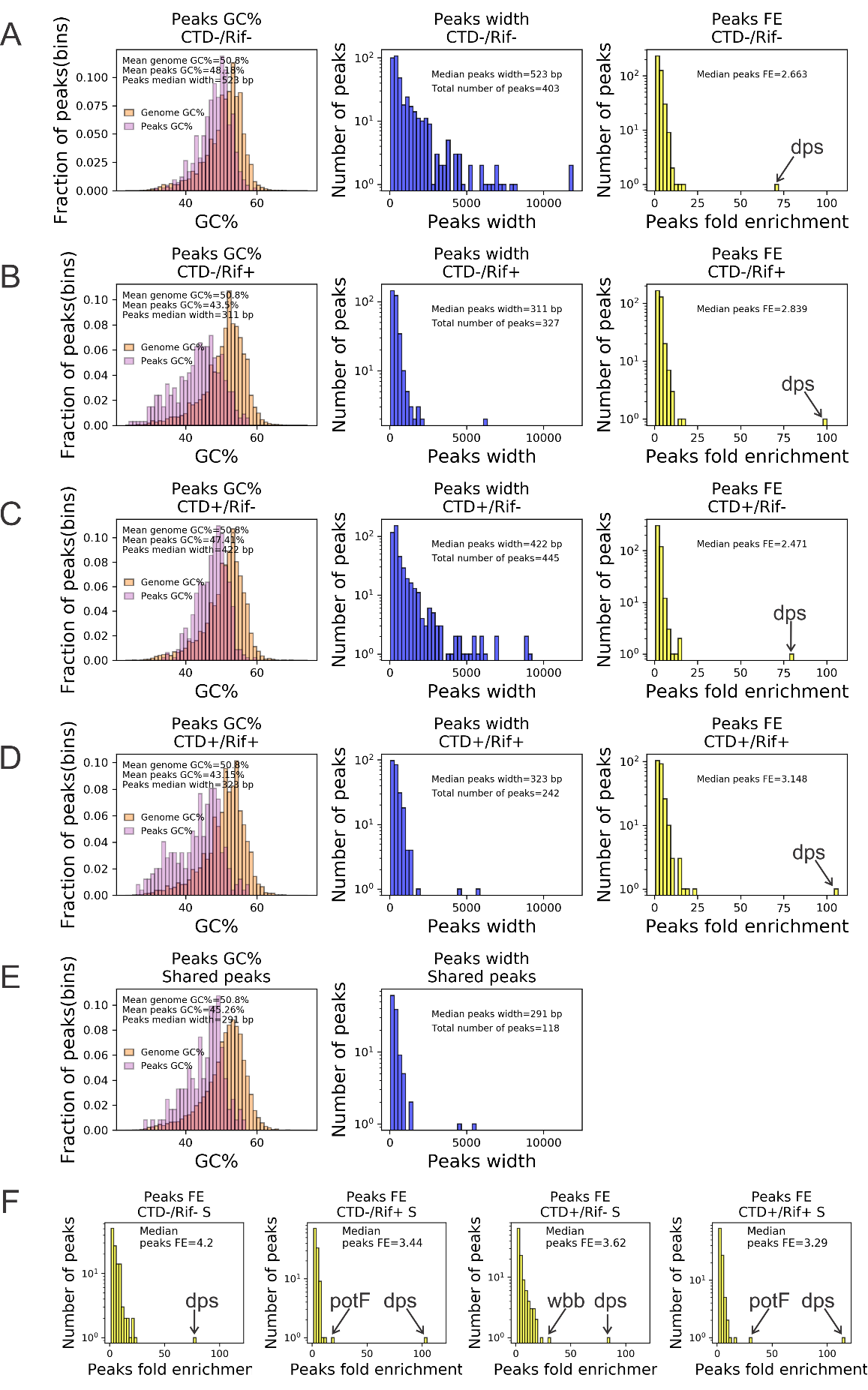

**Figure S3** related to Figures 3 and 7. Association between transcription, RNAP, TopoI, and DNA-gyrase. (**A**) Correspondence among biological replicas of *E. coli* RNA-Seq experiments performed for cells growing exponentially in LB medium at 37^o^C: EP1, EP2, EP3. Heatmap represents pair-wise Pearson correlation between datasets (expression level of TUs expressed in FPKM). Totally, 2123 TUs were analyzed, TUs with extremely high level of expression (rRNAs and tRNAs) were excluded from the analysis. (**B**) Distribution of TUs length. Totally, 1673 TUs were analyzed (**C**) Distribution of TUs expression level. rRNA has an extremely high expression level. Lower panel demonstrates a zoom-in of TUs with expression level lower 1000 FPKM. (**D**) Correlation between RpoC fold enrichment and transcription level of TUs. (**E**) Correlation between RpoC fold enrichment and RpoB signal for TUs. (**F**) Number of RpoC peaks (our data) overlapping with RpoB peaks (data of Kahramanoglou and colleagues [1]). Venn diagram represents an overlap of RpoC peaks (3635 in total) and RpoB peaks (2357 totally). Raw sequencing data for RpoB (β-subunit of RNAP) ChIP-Seq performed for exponentially growing *E. coli* MG1655 is taken from [1] and mapped on *E. coli* DY330 genome. Due to lacking a mock control for ChIP-Seq, regions having coverage depth > 450 were defined as RpoB peaks (~3x excess over background coverage depth). (**G**) Number of RpoB peaks overlapping with EcTopoI peaks. Venn diagram represents an overlap of RpoB peaks (2357 in total) and EcTopoI peaks (403 totally). (**H**) Violin plots of RpoB coverage depth in EcTopoI peaks regions and other sites of *E. coli* genome (left). Violin plots of EcTopoI enrichment in RpoB peaks regions and other sites of genome (right). Mean and median are indicated by black and blue lines, respectively. The statistically significant difference between means (t-test, p-value<<10e-3) is indicated by stars. (**I**) Signal of RpoB in TUs, their upstream (left) and downstream (right) regions. The metagene plot shows the distribution of average ChIP-Seq signal for all TUs, highly-expressed (HETU), and low-expressed (LETU) sets. The number of TUs used for analysis in each group is indicated in parentheses. The two insets show the zoom-in views of RpoB enrichment near transcription start and termination sites. (**J**) Metagene plot represents enrichment of MsmRNAP over the active transcription units (HETU), silent transcription units (LETU) and all transcription units (All TUs). Number of TUs considered for each set indicated in parentheses. Transcription level of *Mycobacterium smegmatis* TUs is assessed based on RNA-Seq data taken from [13], MsmRNAP ChIP-Seq data is taken from [11]. (**K**) Metagene plot represents enrichment of MsmTopoI over sets of TUs. MsmTopoI ChIP-Seq data is taken from [12]. (**L**) Venn diagram represents number of overlapping RNAP and TopoI peaks in *Mycobacterium smegmatis*. (**M**) Enrichment of MsmRNAP in regions occupied by MsmTopoI peaks and enrichment of MsmTopoI in regions occupied by MsmRNAP peaks. (**N**) Metagene plot represents enrichment of MtbRNAP over the active transcription units (HETU), silent transcription units (LETU) and all transcription units (All TUs). Number of TUs considered for each set indicated in parentheses. Transcription level of *Mycobacterium tuberculosis* TUs is assessed based on RNA-Seq data taken from [10]. MtbRNAP ChIP-Seq data is taken from [10]. (**O**) Metagene plot represents enrichment of MsmGyrase over sets of TUs. MtbGyrase ChIP-Seq data is taken from [9]. (**P**) Venn diagram represents number of overlapping RNAP and gyrase peaks in *Mycobacterium tuberculosis*. (**Q**) Enrichment of MtbGyrase in regions occupied by MtbRNAP peaks and enrichment of MtbRNAP in regions occupied by MtbGyrase peaks. (**R**) Metagene plot represents enrichment of SpTopoI and SpRNAP over a set of all *S. pneumoniae* TUs. ChIP-Seq data is taken from [14].

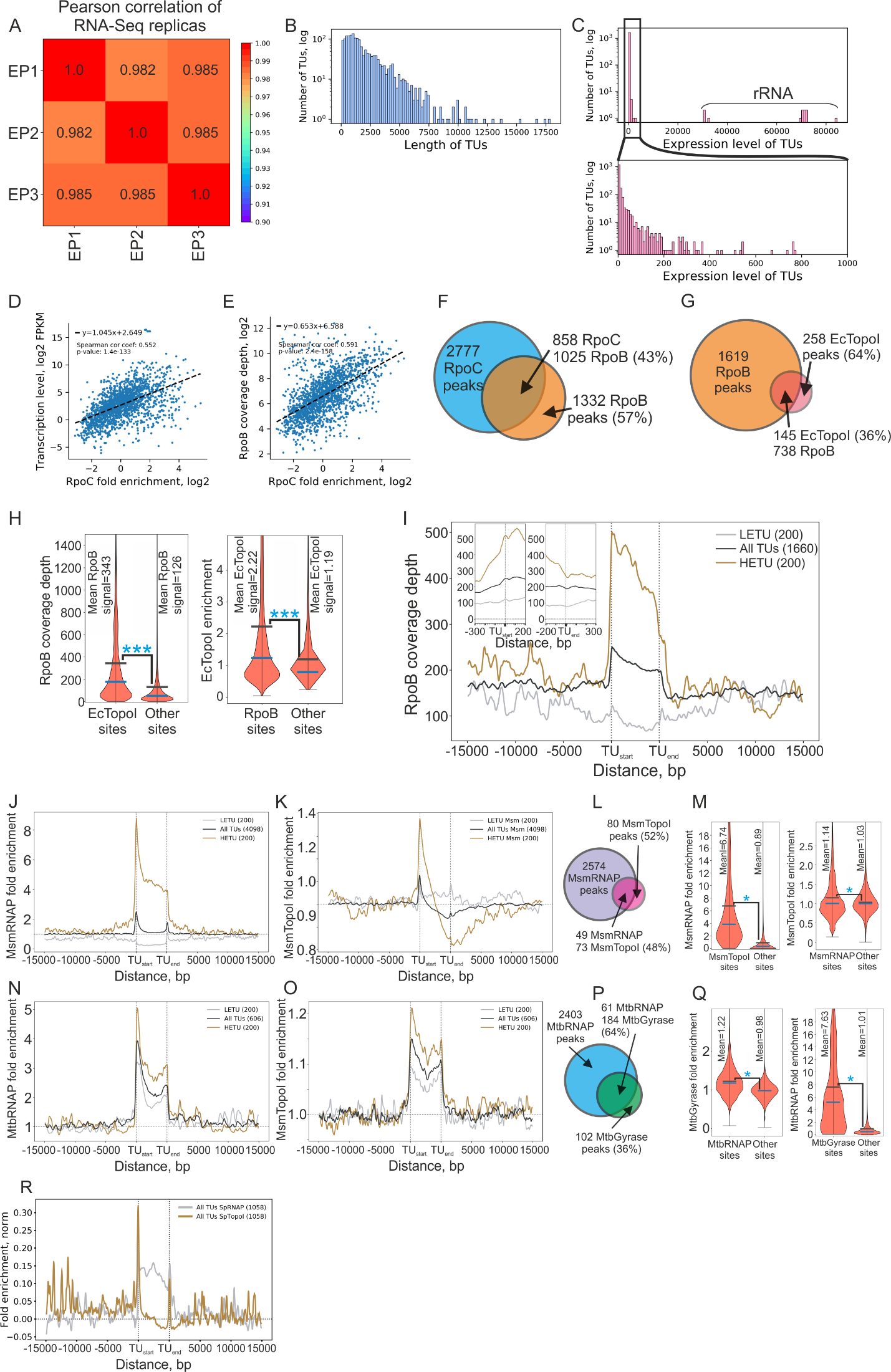

**Figure S4** related to Figure 3 and 7. Monte-Carlo simulation of peak sets overlay. (**A**) Sampling the overlay of EcRNAP (RpoC) and EcTopoI peak sets. (**B**) Sampling the overlay of EcRNAP (RpoB) and EcTopoI peak sets. (**C**) Sampling the overlay of EcRpoC and EcRpoB peak sets. (**D**) Sampling the overlay of EcRNAP (RpoC) and EcTopoI peak sets for Rif+ conditions. (**E**) Sampling the overlay of MsmRNAP and MsmTopoI peak sets. (**F**) Sampling the overlay of MtbRNAP and MtbGyrase peak sets.

Simulations are performed using **Peak_overlap_simulation.py** script, in each test 10000 simulations were performed.

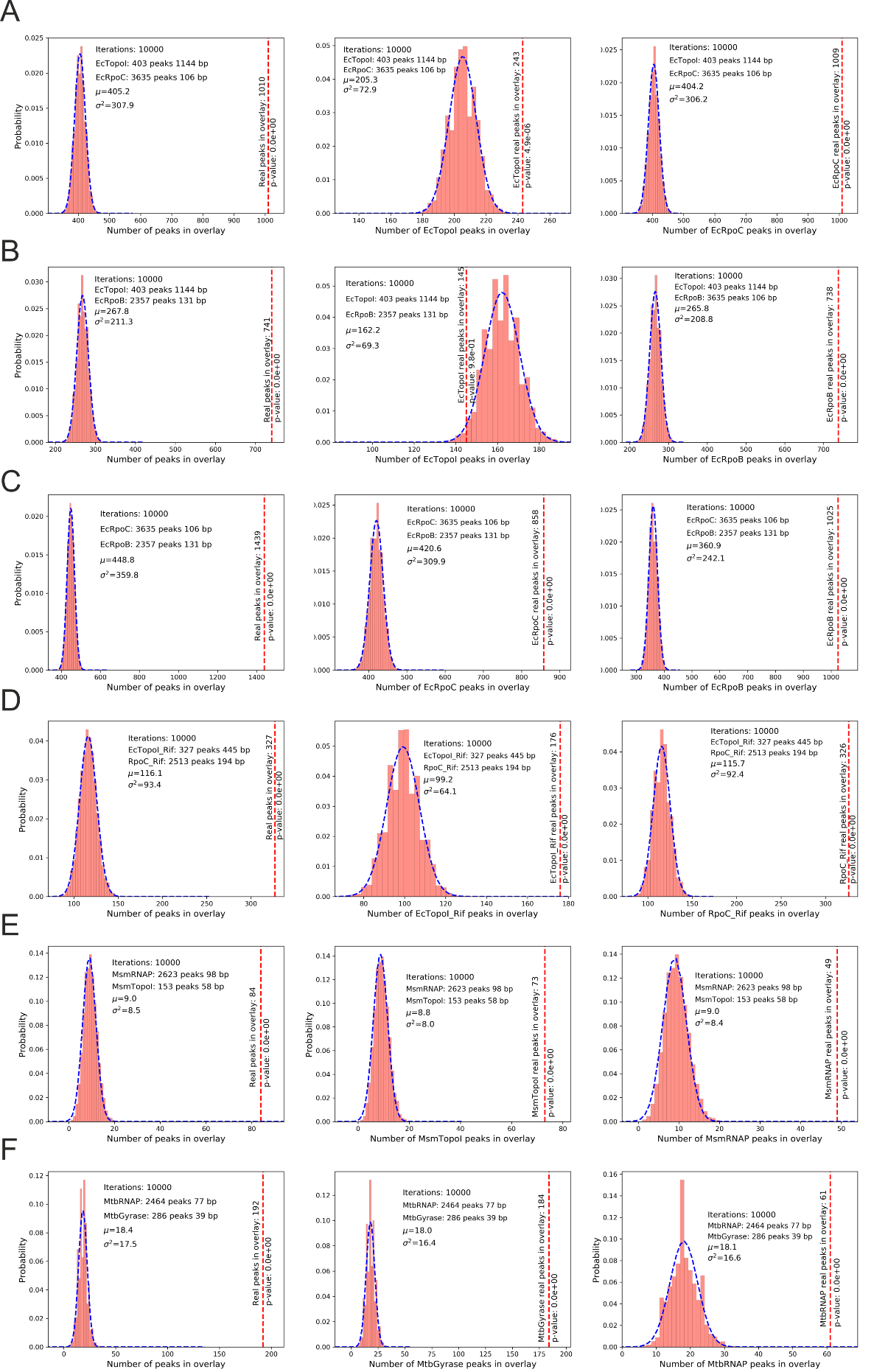

**Figure S5** related to Figures 3 and 6. (**A**) Long-term overexpression of 14kDa CTD is toxic for *E. coli*. CFU counting of *E. coli* DY330 transformed with pCA25 14kDa CTD or pCA25 GFP or without plasmid (- plasmid). Overnight cultures were grown in pure LB medium (-Glc/-IPTG), LB supplemented with either 0.5% Glc (Glc 0.5%/-IPTG) or 1mM IPTG for plasmid induction (-Glc/IPTG 1mM). CFU were counted by a serial dilution method on plates with corresponding medium. Experiments were performed in triplicates. Barplot (on the right) represents quantification of a CFU number (see CFU values in **STab. 8**). (**B**) CFU counting of *E. coli* DY330 transformed with pCA25 14kDa CTD. Culture was grown in LB supplemented with antibiotics until reaching OD_600_=0.2. Then the culture was bisected and one half was induced with 1mM IPTG. After 1h of additional culturing CFU were counted on LB plates without the inducer by a serial dilution method. Experiments were performed in triplicates. Barplot (on the right) represents quantification of a CFU number (see CFU values in **STab. 7**). (**C**) Validation of protein overexpression for EcTopoI 14kDa CTD overexpression from pCA25 14kDa CTD plasmid. Bands correspond to 14kDa CTD (validated by MS) are marked with black triangles. (**D**) Pull-down experiments with His-tagged CTD as a bait. Pull-down expreminets were performed with *E. coli* DY330 topA-SPA and *E. coli* DY330 rpoC-TAP strains. Protein bands were analyzed with MS (Coumassie gel – on the left), RpoC and EcTopoI were detected by Western blotting (on the right) using primary anti-FLAG monoclonal antibodies produced in mouse and secondary anti-mouse antibodies conjugated with HRP produced in rabbit. Protein bands examined by MS are shown by enumearted black triangles on the left gel: RNAP subunit β – bands 1 and 7, RNAP subunit β’ – bands 1 and 6, RNAP subunit α – bands 3 and 8 (see table on the right). EcTopo-SPA and RpoC-TAP are shown by white and black triangles corrspondingly on the right blot. (**E**) Pull-down experiments with EcTopoI-SPA. As a control *E. coli* DY330 strain with native (non-tagged) *topA* was used (*wt*). EcTopoI-SPA is marked with a white triangle; RNAP subunits β and β’ are marked with a black triangle. (**F**) Pull-down experiments with EcTopoI-SPA. Pull-down was performed upon over-expression of GFP or 14kDa CTD from corresponding pCA25 plasmids. As a control *E. coli* DY330 strain with native (non-tagged) *topA* was used (*wt*). EcTopoI-SPA is marked with a white triangle; RNAP subunits β and β’ are marked with a black triangle. Quantification of the pull-down experiments is shown on the right (see values in **STab. 10**). Bands intensity was measured using Photoshop. (**G**) Short-term (1 h**)** CTD overexpression leads to R-loops accumulation. Dot-blot picture is on the left; quantification of the dot-blot signal is shown on the right (see intensity relative values in **STab. 9**). Samples were treated with RNAse III to remove double-stranded RNA. For dot-blot procedure details see Supplementary Materials & Methods. (**H**) The toxicity of the 14 kDa CTD may be caused by alterations of transcription or by degradation of mRNA. Additionally, the effects of CTD on EcTopoI enrichment distribution revealed by ChIP-Seq may be caused by redistribution of RNAP itself. To test these possibilities, we probed transcript coverage over *topA* transcript with RT-qPCR 1-hour post-induction of 14 kDa CTD expression (sequences of the primers see in **STab. 2**). Bars represent coverage normalized on a 5’-proximal region of a gene (topA1 pair of primers). Bars colors correspond to conditions of experiment: green – non-induced cells with CTD expression from pCA25 plasmid repressed by 0.5% glucose, blue – cells harboring pCA25 CTD induced with 1mM IPTG. Below is a map of topA gene with CTD fragments and pairs of primers indicated. (**I**) Abundance of transcripts over the transcription unit comprising of *rpmH-rnpA-yidD-yidC* genes probed by RT-qPCR for untreated condition and for 1-hour induction of 14 kDa CTD. Bars represent coverage normalized on a 5’-proximal region of a transcription unit (rpmH1 pair of primers). See raw qPCR data in **STab. 16**.

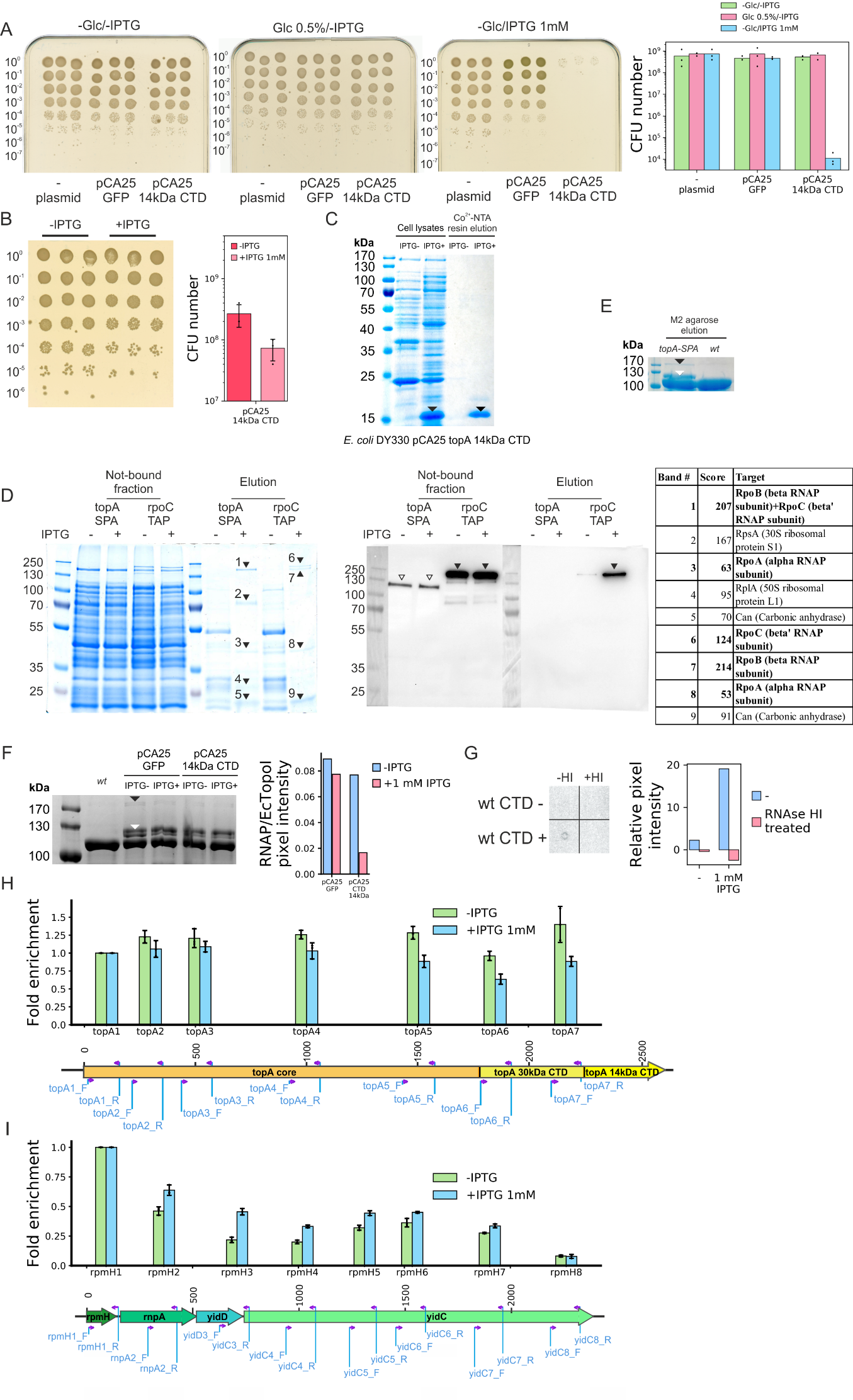

**Figure S6** related to Figure 4. Factors associated with high or low EcTopoI signal in intergenic regions. (**A**) Distribution of length of intergenic regions in *E. coli* W3110 genome. In total 3459 intergenic regions were identified. For further analysis regions has been filtered by length (1000bp>x>50bp), by expression of adjacent genes (non-zero expression level). Additionally, *dps* gene and genes of rRNA operons were excluded due to extremely high fold enrichment of EcTopoI for *dps* and extremely high expression level of rRNA operons. Finally, 1529 intergenic regions were analyzed. (**B**) Effect of a high level of expression of adjacent genes. Intergenic regions were classified as high expression if the cumulative expression of adjacent genes was higher than 5 (**EL>5**) or as low expression if opposite true (**EL<5**). Then, for each group of intergenic regions RNAP fold enrichment (**pink**), expression level of adjacent genes (**yellow**), and fold enrichment of EcTopoI in Rif-/CTD- (**red**) and Rif+/CTD- (**green**) conditions were identified. Violin plots demonstrate distribution of parameter values, vertical axis is log-scaled. Sample size is indicated above the plots, means are marked with a horizontal black line and written explicitly below (), medians are marked as a blue horizontal line. If present, statistically significant differences between means are visualized with dashed lines with stars (p-value<0.01, Welch t-test). (**C**) Effect of a high fold enrichment of RNAP in an intergenic region. Regions were classified as high RNAP signal if fold enrichment was higher than 2 (**RNAP>2**) or as low RNAP signal if opposite true (**RNAP<2**). Color coding and marking is as for picture B. (**D**) Intergenic regions were classified by orientation of adjacent genes: genes can lay in a same orientation on positive or negative strand (🡪🡪 or 🡨🡨); genes can by divergent (🡨🡪) or convergent (🡪🡨). (**E**) Intergenic regions containing annotated sites of transcription factors (**+TF**) and not containing (**-TF**). Information about localization of transcription factor sites is taken from RegulonDB [2]. (**F**) Intergenic regions were classified by the following: 1) both adjacent genes encode membrane protein (**MM**), 2) if only one of them encodes membrane protein (**M-**), 3) if neither of them encodes membrane protein (--). Information about subcellular localization of *E. coli* proteins was taken from Ecocyc database. (**G**) Same analysis as in F, but information about subcellular localization of *E. coli* proteins was taken from PSORT database 4.0 [3]. (**H**) Effect of a combination of the “expression-related” features – level of expression and signal of RNAP: high expression and high RNAP signal intergenic regions (**EL>5, RNAP>5**) compared with low expression and low RNAP signal set (**EL<5, RNAP<2**). (**I**) Effect of a combination of “expression-non-related” features - orientation of adjacent genes (🡨🡪 or ~🡨🡪); presence of transcription factors sites (**+TF** or -**TF**); membrane localization of protein encoding by the adjacent genes (**M** or **~M**). Information about localization of transcription factor sites is taken from RegulonDB [2]. Information about the subcellular localization of *E. coli* proteins was taken from PSORT database 4.0 [3].

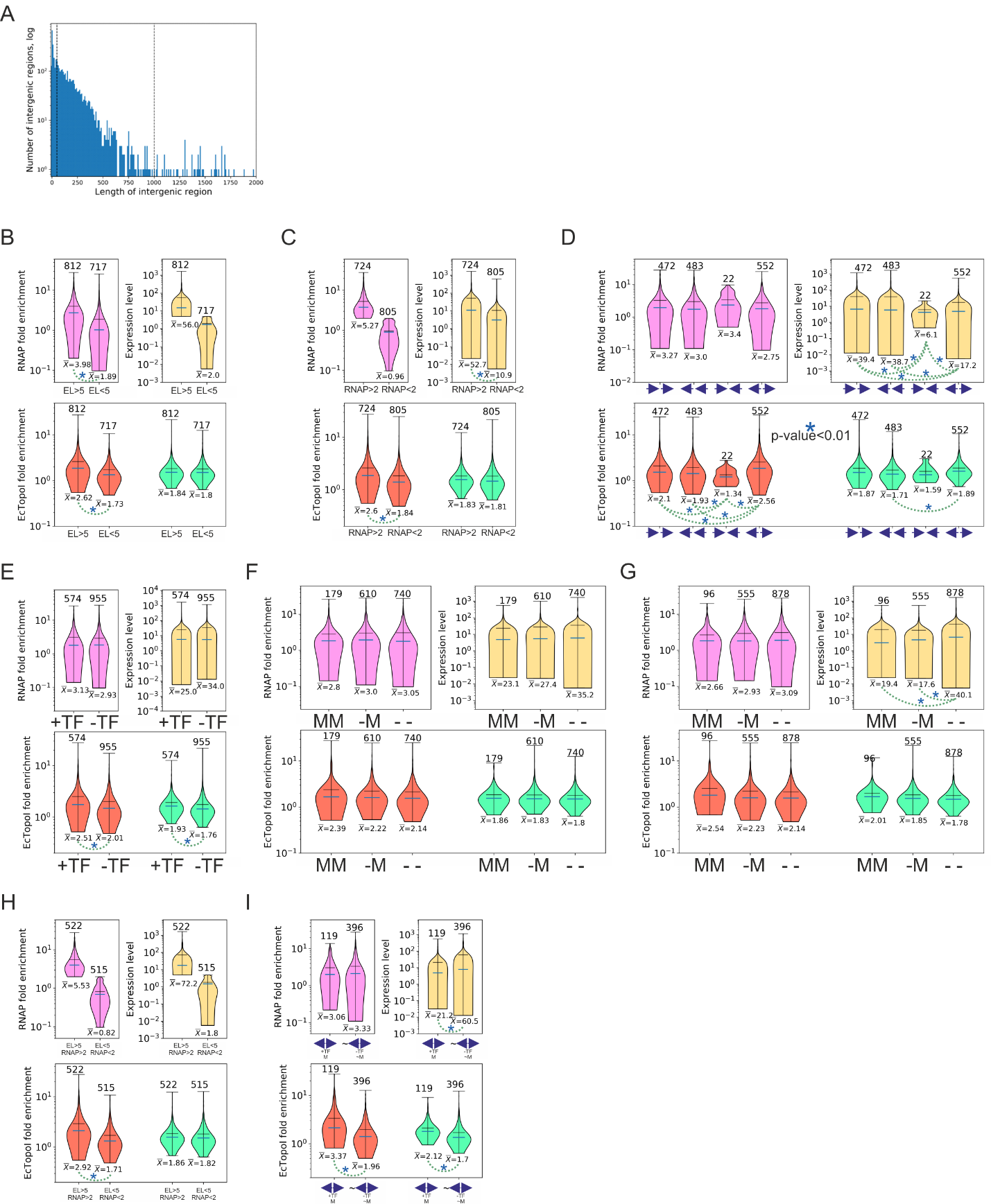

**Figure S7** related to Figure 4. Association of EcTopoI ChIP-Seq enrichment with intergenic regions contain transcription factor sites of different types (**A**) and promoters recognized by different sigma-factors (**B**). (**A**) Intergenic region is classified by a type of transcription factor site it contains: **LexA**, **Fur**, **NsrR**, **CRP**, **PhoP**, **OxyR**, **IHF**, **ArcA**, **RcsB**, **SoxS**, **PhoB**, **CpxR**, **Fis**, **NarL**, **FNR**, **MarA**, **Cra**. Only transcription factors with annotated sites in more than 15 intergenic regions were considered. Totally, 1529 sites were analyzed. For each set of intergenic regions RNAP fold enrichment (**pink**), expression level of adjacent genes (**yellow**), and fold enrichment of EcTopoI in Rif-/CTD- (**red**) and Rif+/CTD- (**green**) conditions were identified. Violin plots demonstrate distribution of the parameter values, vertical axis is log-scaled. Sample size is indicated above the plots, means are marked with a horizontal black line on violin-plots and written explicitly below the plots (), medians are marked as a blue horizontal line on plots. If present, statistically significant difference between mean of sets visualized with a dashed line (p-value<0.01, Welch t-test). Annotation of transcription factor sites taken from RegulonDB [2]. Set comprising of all intergenic regions is marked with a black arrow. (**B**) Intergenic region was classified by a type of a sigma factor recognizing annotated promoters within the region. Totally, 1529 regions were analyzed. If promoter(s) in an intergenic region is/are recognized by a same type of a sigma factor, regions were classified as **Sigma70**, **Sigma38**, **Sigma54**, **Sigma32**, **Sigma24**, **Sigma28** depending on a type of a factor. If promoter(s) is/are recognized by several different sigmas, region was marked as **Mixed**. If promoter’s specificity is unknown, region was classified as **Unknown**. If all the promoters are annotated as unknown within some intergenic region, it was called **Only unknown**. For each set of intergenic regions RNAP fold enrichment (**pink**), expression level of adjacent genes (**yellow**), and fold enrichment of EcTopoI in Rif-/CTD- (**red**) and Rif+/CTD- (**green**) conditions were identified. Violin plots demonstrate distribution of parameter values, vertical axis is log-scaled. Sample size is indicated above the plots, means are marked with a horizontal black line and written explicitly below (), medians are marked as a blue horizontal line. If present, statistically significant differences between means of subsets are visualized with dashed lines (p-value<0.01, Welch t-test). Annotation of *E. coli* promoters was taken from RegulonDB [2].

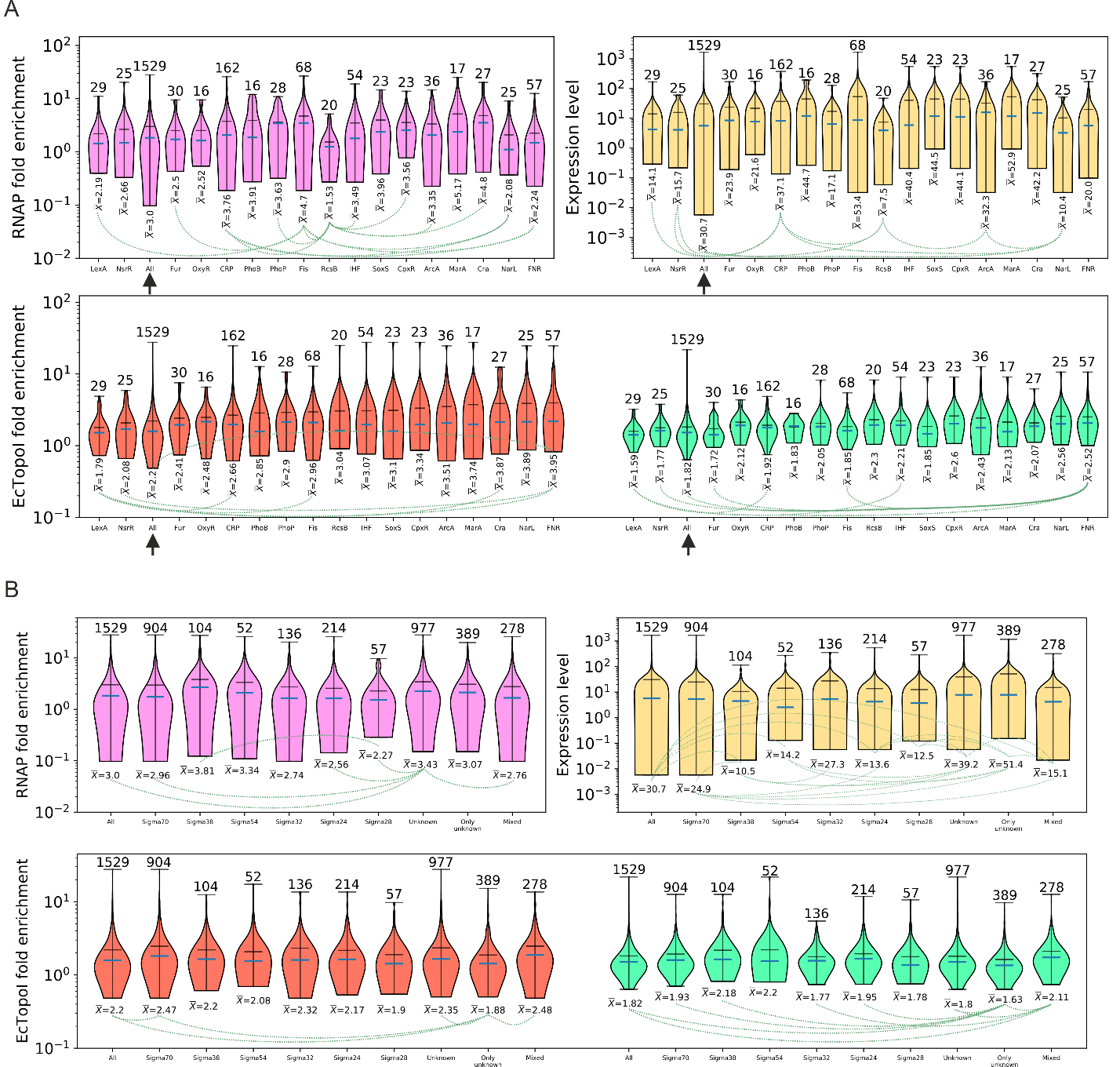

**Figure S8** related to Figure 4. (**I**) DNA-gyrase enrichment in transcription units, their upstream and downstream regions. Enrichment is shows for all TUs (black curve), highly-expressed (HETU, orange curve), and low-expressed (LETU, grey curve) sets. The number of TUs in each group is indicated in parentheses. DNA-gyrase data is from [4], experiments with Cfx as a gyrase poison were taken. Data for gyrase in native conditions and for cells pre-treated with Rif are shown with solid and dashed curves respectively. (**II**) Snapshots from IGV genomic browser representing EcTopoI fold enrichment around rRNA operons for Rif-/CTD- conditions. rRNA operons listed from A to H, genes corresponding to the operons colored in red. (**III**) Averaged fold enrichment over 7 rRNA operons. Solid gray line represents Rif-/CTD- ChIP-Seq experiments.

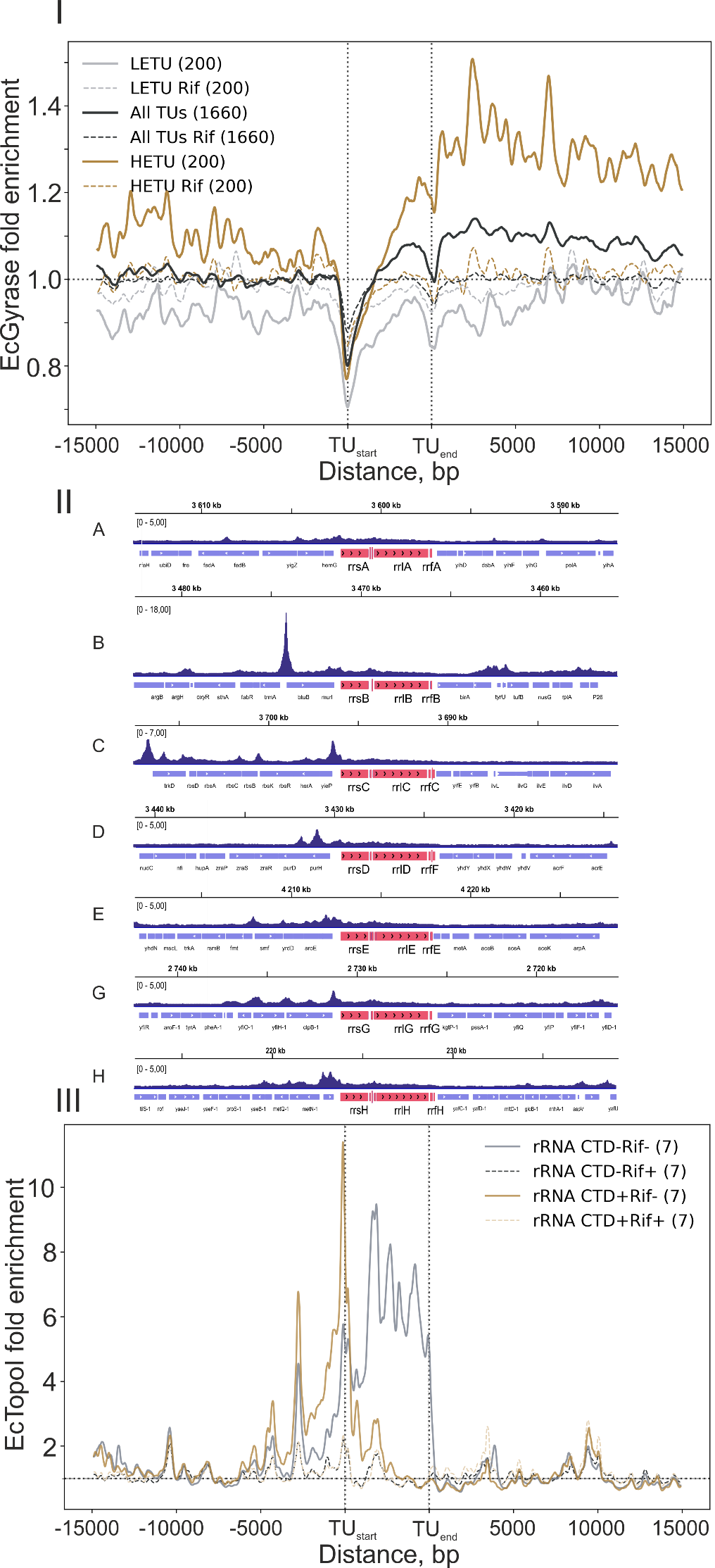

**Figure S9** related to Figure 5. EcTopoI Topo-Seq and Topo-Seq data analyis. (**A**) Detection of SOS-response in cells expressing EcTopoI G116S M320V double-mutant from pBAD33 and in cells expressing EcTopoI 14-kDa CTD from pCA25 plasmid. For the detection a *E. coli* CSH50 strain [5] transformed with the indicated plasmids was cultivated on MacConkey agar supplemented with lactose and 10 mM Ara. As a control the strain was transformed with pCA25 GFP. (**B**) Growth curve of *E. coli* DY330 topA-SPA harboring pBAD33 EcTopoI G116S M320V plasmid in LB supplemented with 0.5% glucose. At OD_600_~0.2 the culture was bisected and 10 mM Ara was added to one half (+ Ara 10 mM, light blue curve), the second half served as a non-induced control (- Ara, black curve). Shade is represented 0.95 confidential interval for mean of 3 biological experiments. See OD_600_ values in **STab. 11**. (**C**) Overexpression and purification of EcTopoI G116S M320V from pBAD33 plasmid during the Topo-Seq procedure (for details see **Materials &** **Methods** section). Bands correspond to EcTopoI G116S M320V are marked with white triangles (verified by MS). (**D**) Correspondence between EcTopoI ChIP-Seq peaks, topoisomerase cleavage sites (TCSs) identified with Topo-Seq, and EcRpoC ChIP-Seq peaks. Venn diagram demonstrating number of overlapping peaks and TCSs. (**E**), (**F**) Monte-Carlo simulation of the overlay beetween EcTopoI peaks and TCSs sets and between EcTopoI TCSs and EcRpoC peaks correspondingly. Simulations are performed with **Peak_overlap_simulation.py** script, 10000 simulations were performed. (**G**) Metagene plot represents enrichment of EcTopoI cleavage activity (N3E) over all transcription units (All TUs), silent transcription units (LETU), active transcription units (HETU), and rRNA operons (rRNA). Number of TUs considered for each set is indicated in parentheses. Enrichment of EcTopoI cleavage activity was obtained in two steps. First, N3E data for “+Ara Mock” sample was subtracted from “+Ara IP” sample and “-Ara Mock” sample was subtracted from “-Ara IP” sample. Second, resultant “-Ara [N3E IP – N3E Mock]” track was subtracted from resultant “+Ara [N3E IP – N3E Mock]” track. The final track was utilized in metagene analysis. To smooth the data, cleavage signal was binned (bin width 200 nt). For the cleavage tracks confidential interval ±SEM is shown. Barplots represent a comparison of EcTopoI cleavage signal in different regions relative to TUs. Differences in the signal were tested using Welch t-test. Significance is indicated by stars. Error bars are represented by ±SEM.

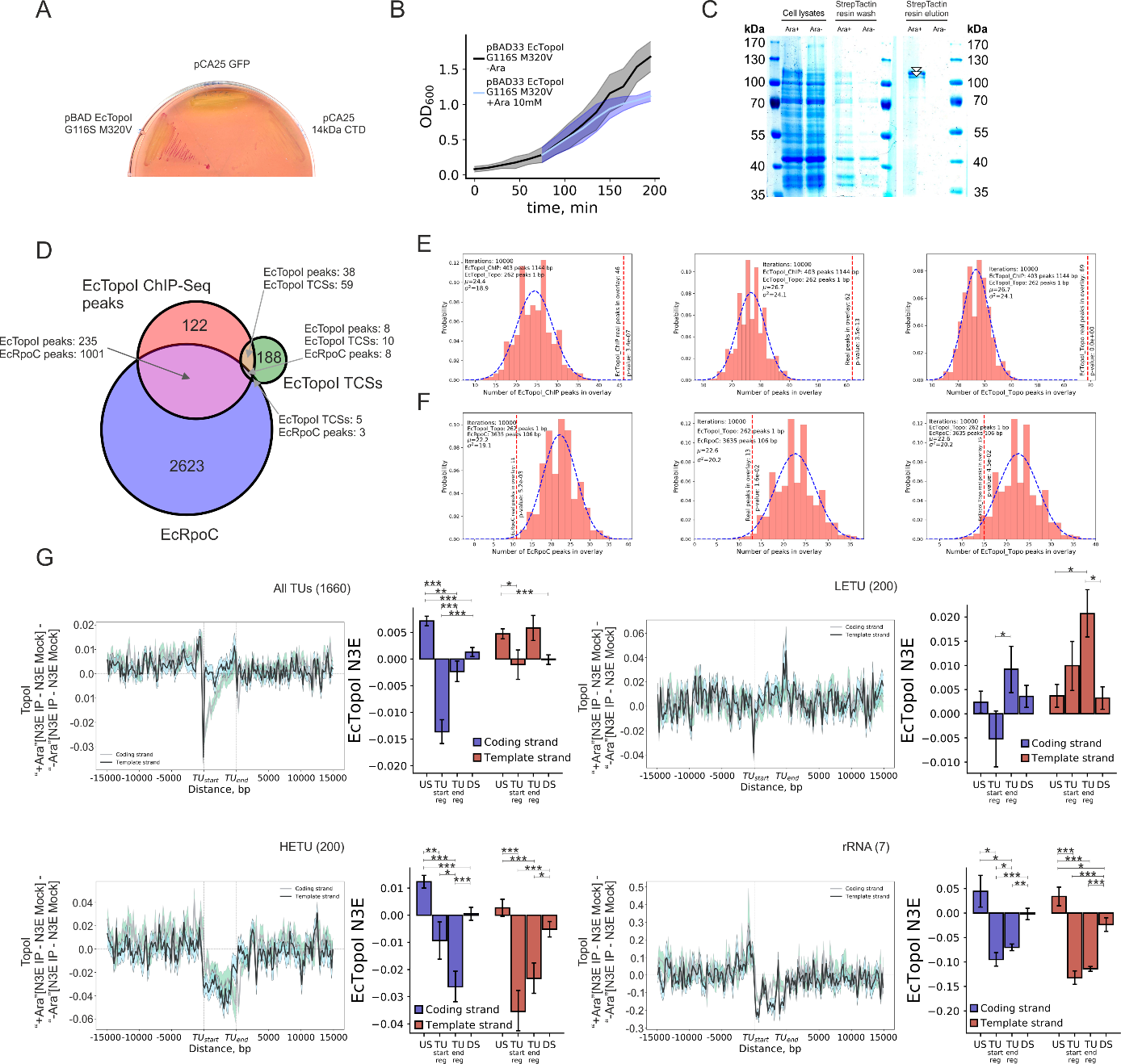

**Figure S10** related to Figure 5. EcTopoI binding motif. (**A**) Binding motifs of EcTopoI identified in 3 independent runs of ChIPMunk for different ChIP-Seq experiments: -CTD/-Rif, -CTD/+Rif, +CTD/-Rif, +CTD/+Rif. Each motif is given for “+” and “-” strands. 3 independent runs were performed in accordance with ChIPMunk manual, because of the stochastic nature of the algorithm [6]. Significance of the motifs expressed as a Kullback Discrete Information Content value (KDIC) is indicated for each algorithm run. (**B**) Number of peaks identified by MACS2 [7] taken for motif discovery and number of peaks containing motif indentified by ChIPMunk. (**C**) Motifs of *E. coli* DNA-binding proteins found to be similar with EcTopoI motif by Tomtom [8]. Also, statistically significant (p-value<0.05) similarity was found between EcTopoI motif and *E. coli* Fur, ArcA, *V. cholerae* ToxT, *S. meliloti* LexA motifs. (**D**) ChIP-qPCR validation of ChIP-Seq peaks near *dps* and *potF* genes. Dps and potF regions have dramatically increased fold enrichment in comparison with the two control regions – dif and H2394. See raw qPCR data in **STab. 17.** (**E**) EMSA competition experiments: 0.4 pmole of a labeled oligonucleotide (final concentration 20 nM) was mixed with 8 pmole of EcTopoI (final concentration 400 nM and this is a saturating concentration) and increasing amount of non-labeled competitor oligonucleotide (0-12.8 pmole, that correspond to final concentrations 0-640 nM) in 20 μL rxn. Separation was performed in 10% acrylamide gel in TGB with magnesium (see **Materials &** **Methods**). Ratio between labeled oligonucleotide and a non-labeled competitor is indicated above a gel. Oligonucleotide pairs are indicated below gels (Cy5-labeled oligo: non-labeled oligo). Dye used to track separation progression is visible as a black dot in the first lanes of several gels: Cy5-Consensus (F):poly-T and Cy5-Poly-A:Consensus (R). Minimal concentration at which a non-labeled competitor starts to displace a labeled oligo is indicated by a red star.

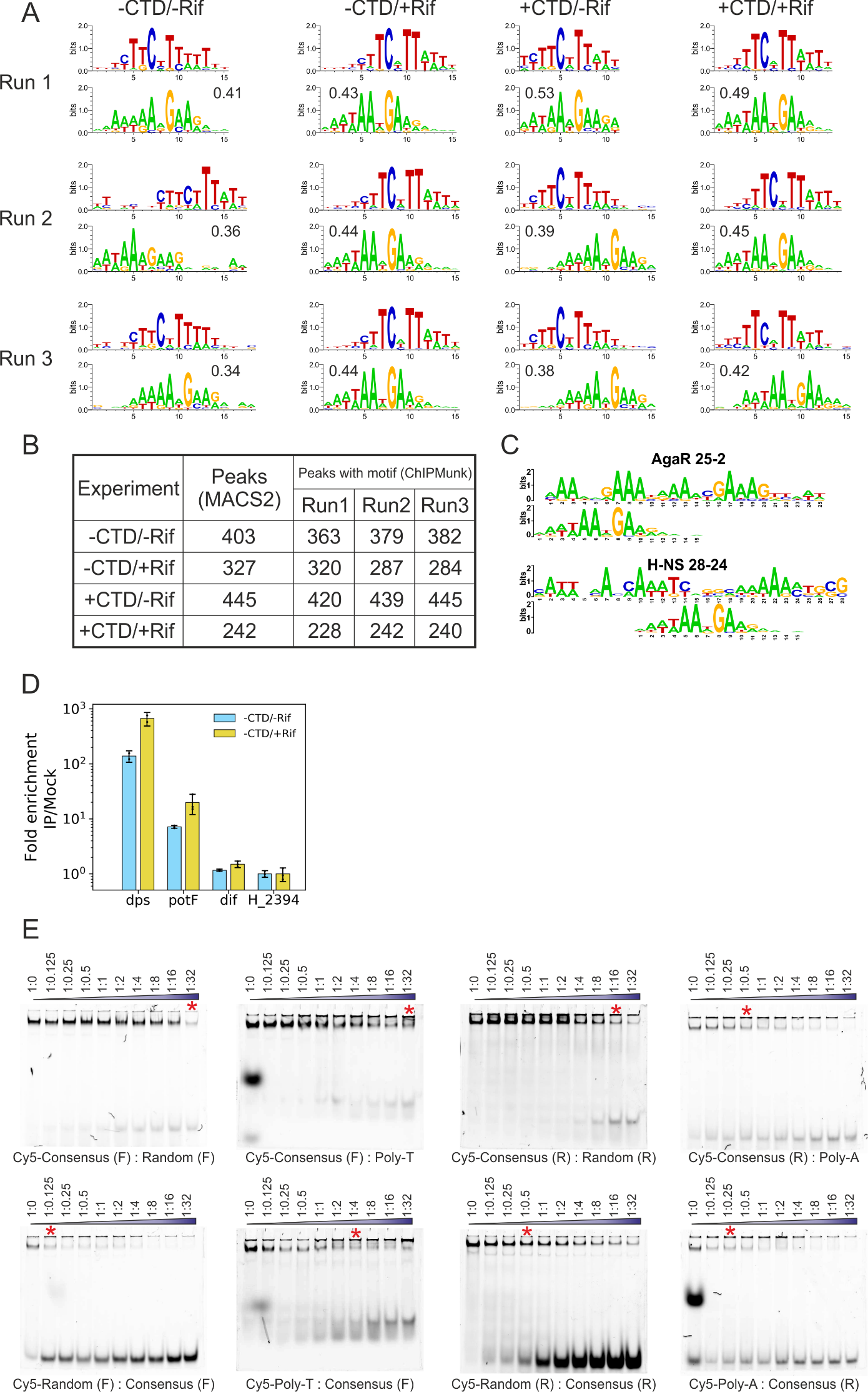

**Figure S11** related to Figure 6. Viability of *E. coli* BW25113 strains carrying truncated versions of a *topA* gene. Competition between BW25113 *wt* and BW25113 *topA* mutants (*topAΔ11* or *topAΔ14*). The strains mixture was cultivated in LB and every 24h inoculated in a fresh LB batch. Strain abundance was monitored with strain-specific primers using qPCR. Welch t-test was used to assess differences between mutant/wt ratio in the beginning of the competition experiment (day 0) and at the end of the experiment (day 5). See raw qPCR data in **STab. 15**

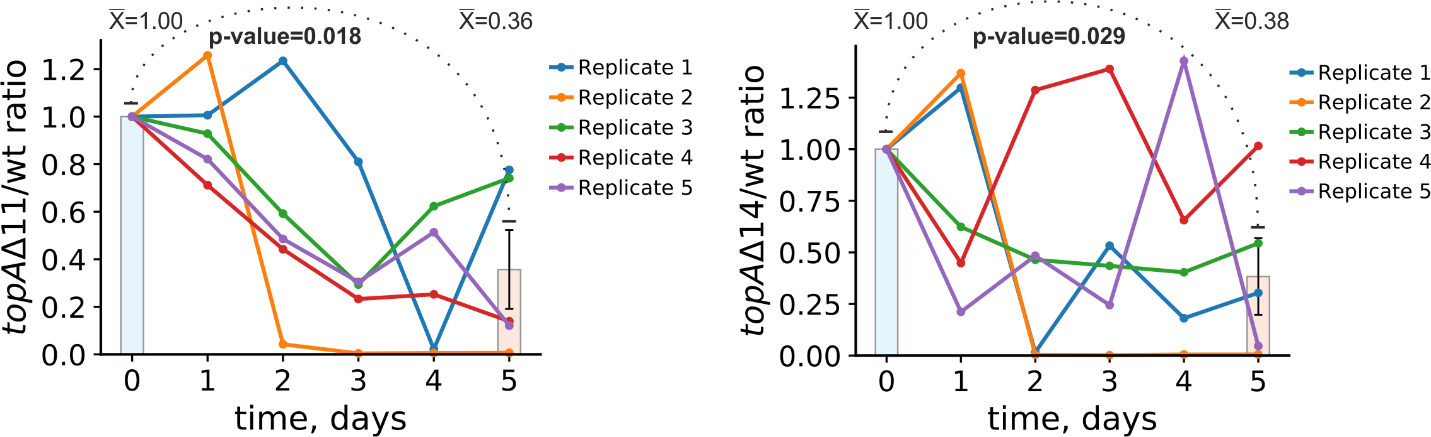

**Table S1**. Primers used for cloning, recombineering or qPCR; Primers used for quantification of transcripts coverage with RT-qPCR; Sequences of oligonucleotides used in binding and cleavage in vitro experiments.

| **Primer** | **Sequence** | **The length of amplicon, bp** | **Description** |
| --- | --- | --- | --- |
| topA_NcoI_fw | GTAAGGTGACCATGGGTAAAGCTCTTG | 2508 | topA amplification fragment 1 |
| NcoI_mut_rev | GGCTAAAGCGAACCACGGTCTTATTACC |  |  |
| NcoI_mut_fw | GGTAATAAGACCGTGGTTCGCTTTAGCC | 170 | topA amplification fragment 2. StrepII-coding sequence indicated by capitals. |
| topA_HindIII_strepII_rev | atataaagcttttaTTTTTCGAACTGCGGGTGGCTCCAcgcgcttttttttccttcaacccatttgccatc |  |  |
| topA_NdeI_fw | GTAAGGTGCatATGGGTAAAGCTCTTGTCATCG | 2654 | topA amplification |
| topA_strepII_HindIII_rev | ttcgatatcaagcttTTATTTTTCGAACTGCG |  |  |
| TopoA_14_kDa_CTD_BamHI_fw | GGATCGCATCACCATCACCATCACGGATCCGGCGAAGTGGCACCACCG | 423 | EcTopoI CTD amplification |
| TopoA_CTD_HindIII_rev | aacaggagtccaagctcagctaattaagctTTATTTTTTTCCTTCAACCCATTTGCC |  |  |
| pCA24_core_NcoI_rev | AATCTAACCATGGATCCGTGATGGTGATGGTGATGC | 4515 | Amplification of the pCA24 core |
| pCA24_core_HindIII_fw | AAACTGGGGCACAGATGAAGCTTAATTAGC |  |  |
| topA_XbaI_RBS_fw | tatataactctagaaataattttgtttaactttaagaaggagatataccatgggcagcagccatcatcatcatcatcacagcagcggcctggtgccgcgcggcagccataTGGGTAAAGCTCTTGTCATCG | 470 | Introduction of G116S and M320V mutations in *topA*. Introduction of C-terminal strepII-tag. Cloning into pBAD33. |
| topA_G116S_out_rev | ATGCAATGGCTTCAGATTCGCGGTCAAGGTCGG |  |  |
| topA_G116S_in_for | TTGACCGCGAATCTGAAGCCATTGCATGGCACC | 636 |  |
| topA_M320V_in_rev | AGTCGGTACGCACGTAAGTGATATAGCCTGCTTCATAC |  |  |
| topA_M320V_out_for | ATCACTTACGTGCGTACCGACTCCACTAACC | 1691 |  |
| topA_strepII_HindIII_rev | atatcaagcttTTATTTTTCGAACTGCGGGTGGCTCCAC |  |  |
| dps_850_F | CGGTGTAGAGGAAAAGTAGCGAGAAATTCTGC | 124 | Sequence under the highest EcTopoI ChIP-Seq peak. Used for ChIP-qPCR and EMSA: dps amplicon. |
| dps_850_R | ATGGGGTCTACGCTGACAGTACG |  |  |
| potF_895_F | CGGGAGAAAGTTCTCTTTTCTTACACCG | 125 | Sequence under one the highest EcTopoI ChIP-Seq peak. Used for ChIP-qPCR and EMSA: potF amplicon. |
| potF_895_R | GCGGTCATATCTCTTTCCTTCTGAAAGTTCG |  |  |
| L_2394_F | CCAGTACCAACAGTGCAGAGATC | 110 | Region with low signal of EcTopoI ChIP-Seq. Used for ChIP-qPCR and EMSA: nuoN amplicon. |
| L_2395_R | GGTGAGCCTGTATCTTCACGC |  |  |
| topA_delta_topA66_kanR_F | CACCACCGAAAGAAGATCCGGTGCCATTACCTGAGCTGCCGTGCGAAAAATAATAGTGATCGTGTAGGCTGGAGCTGCTTC | 1613 | For genome editing by recombineering. topA_SPA_kanR_cysB_R is universal for the three forward primers. |
| topA_delta_14kDa_kanR_F | GTAAATACATGGCCTGCACCAACGAAGAGTGTAAAAACACACGTAAGATTTTACGTAACTAATAGTGAATCGTGTAGGCTGGAGCTGCTTC | 1623 |  |
| topA_delta_30kDa_kanR_F | AGTTAGATAAAGCTGAAAAAGATCCGGAAGAGGGTGGTATGCGCCCGAACTAATAGTGAATCGTGTAGGCTGGAGCTGCTTC | 1614 |  |
| topA_SPA_kanR_cysB_R | ATGCAATAAAAAAGGGCCGCTTTCGCGACCCTTTGTTTATAAAAACCTGACAGAATTAAAGGTGAAGAAGTAAAACATATGAATATCCTCC |  |  |
| topA_8_F1 | GTTGAGTCCCCGGCAAAAGCC | 138 | To assess transcripts density over long transcription units with RT-qPCR upon uncoupling of EcTopoI:RNAP complex by overexpression of EcTopoI CTD: *topA* (~2600 bp) and *rpmH-rnpA-yidD-yidC* (~2200 bp). |
| topA_53_R1 | GGTGGAGGTAGAGTCGGCACTC |  |  |
| topA_73_F2 | GGTTGACCCGTGGCACAATTGG | 138 |  |
| topA_118_R2 | GCTTCCCCTTCGCGGTCAAGG |  |  |
| topA_146_F3 | CGATCCGCCAGGCATTTAACAAACC | 137 |  |
| topA_191_R3 | CAGGCCACGAGCGATCTTTTTCC |  |  |
| topA_303_F4 | GATGGCGCAGCGTTTGTATGAAGC | 139 |  |
| topA_352_R4 | CGGACTTTCCGGCAGATATTTCTTACC |  |  |
| topA_476_F5 | CCGTTTCAGTGAAGCATCGCTGG | 145 |  |
| topA_523_R5 | GCCCATTTTTTCCGCATAGAAACGACG |  |  |
| topA_589_F6 | GATGGTTCTGACCAGCATTGACTGC | 138 |  |
| topA_633_R6 | ACCAGGTTAATGGTGGTTTTGCAACG |  |  |
| topA_694_F7 | GGGCGAATTCCGCATTAAAGGTTATGAC | 131 |  |
| topA_740_R7 | CGTGTGTTTTTACACTCTTCGTTGGTGC |  |  |
| Operon_1_rpmH_4_F1 (rpmH_1_F) | GCACTTTTCAACCGTCTGTACTGAAGCG | 140 | To assess transcripts density over long transcription units with RT-qPCR upon uncoupling of EcTopoI:RNAP complex by overexpression of EcTopoI CTD: *topA* (~2600 bp) and *rpmH-rnpA-yidD-yidC* (~2200 bp). |
| Operon_1_rpmH_47_R1 (rpmH_1_R) | GCTTTATTACTTAGAAACGGTCAGACGAGCG |  |  |
| Operon_1_rnpA_44_F2 (rpmH_2_F) | CATCCCCGTATCGGTCTTACAGTCG | 138 |  |
| Operon_1_rnpA_R89 (rpmH_2_R) | CGCCACCACCACGAAATCCATAGC |  |  |
| Operon_1_yidD_37_F3 (rpmH_3_F) | GGAGTGATAAAAGGCAGTTGGTTGACG | 141 |  |
| Operon_1_yidC_9_R3 (rpmH_3_R) | CTAAAAGATTGCGTTGCGAATCCATCG |  |  |
| Operon_1_yidC_67_F4 (rpmH_4_F) | CGTGCTTGATCTGACCATCAACACC | 140 |  |
| Operon_1_yidC_113_R4 (rpmH_4_R) | CGCTCTGTGCCTGATAAATAAACTGCG |  |  |
| Operon_1_yidC_166_F5 (rpmH_5_F) | GTCCTGAAACGTGGTGATTACGCTGTC | 128 |  |
| Operon_1_yidC_208_R5 (rpmH_5_R) | GTATCGAGATGCGGTGGCAGAGTG |  |  |
| Operon_1_yidC_238_F6 (rpmH_6_F) | TGCCGATAACGAAAACCTGAACATCTCTTC | 139 |  |
| Operon_1_yidC_284_R6 (rpmH_6_R) | GGCGATGCCGTTACCCAGATTAGC |  |  |
| Operon_1_yidC_362_F7 (rpmH_7_F) | ACCTTTATCGTTCGTGGCATCATGTACC | 135 |  |
| Operon_1_yidC_401_R7 (rpmH_7_R) | CTGGCTGATACGCTGTTTGTCATCG |  |  |
| Operon_1_yidC_482_F8 (rpmH_8_F) | TCGCCGACCACAGTGACCGACC | 139 |  |
| Operon_1_yidC_528_R8 (rpmH_8_R) | GCTGAATAATGGTTACCAGGTTGCTGACG |  |  |
| BW25113_topA_wt_F | GCCGCTTTCGCGACCCTTTG | 103 | To assess the composition of a mixed culture undergo the competition test (*E. coli* BW25113 vs BW25113 *topA* mutant) by qPCR. |
| BW25113_topA_wt_R | CGACTGGCTGGTCAGCATTTTATGTTGATGG |  |  |
| BW25113_topA_delta11_SPA_F | GAGCTGCCGTGCGAAAAATCCATG | 106 |  |
| BW25113_SPA_qPCR_R | CAAGTGCCCCGGAGGATGAGATTTTC |  |  |
| BW25113_topA_delta14_SPA_F | ACACACGTAAGATTCTGCGCAATTCCATG | 111 |  |
| BW25113_SPA_qPCR_R | CAAGTGCCCCGGAGGATGAGATTTTC |  |  |
| EcTopoI_Consensus_F_Cy5 | Cy5GCGTTTTTTTCTTCATTATTTTTGCG |  | Oligonucleotides used in binding experiments with EcTopoI. Oligos carrying Cy5 labels on 5'-end. |
| EcTopoI_Consensus_R_Cy5 | Cy5CGCAAAAATAATGAAGAAAAAAACGC |  |  |
| Random_F_Cy5 | Cy5GCGCACGTAATCCGCAGTGCGGTGCG |  |  |
| Random_R_Cy5 | Cy5CGCTCCGCACTGCGGATTACGTGCGC |  |  |
| Poly-T_Cy5 | Cy5GCGTTTTTTTTTTTTTTTTTTTTGCG |  |  |
| Poly-A_Cy5 | Cy5CGCAAAAAAAAAAAAAAAAAAAACGC |  |  |
| EcTopoI_Consensus_F | GCGTTTTTTTCTTCATTATTTTTGCG |  | Oligonucleotides used in binding experiments with EcTopoI. Oligos are label-free. |
| EcTopoI_Consensus_R | CGCAAAAATAATGAAGAAAAAAACGC |  |  |
| Random_F | GCGCACGTAATCCGCAGTGCGGTGCG |  |  |
| Random_R | CGCTCCGCACTGCGGATTACGTGCGC |  |  |
| Poly-T | GCGTTTTTTTTTTTTTTTTTTTTGCG |  |  |
| Poly-A | CGCAAAAAAAAAAAAAAAAAAAACGC |  |  |

**Table S2**. NGS datasets used in the study.

| **Dataset** | **ID** | **Link** | **Description** | **Study** |
| --- | --- | --- | --- | --- |
| *E. coli* DY330 RNA-Seq | GSE181687 | <https://www.ncbi.nlm.nih.gov/geo/query/acc.cgi?acc=GSE181687> | Total RNA-Seq for exponentially growing culture in LB. Performed in triplicate. | Current study |
| *E. coli* DY330 RpoC ChIP-Seq | GSE182850 | <https://www.ncbi.nlm.nih.gov/geo/query/acc.cgi?acc=GSE182850> | ChIP-Seq performed for exponentially growing culture in LB. Single replicate. | Current study |
| *E. coli* DY330 EcTopoI ChIP-Seq | GSE181915 | <https://www.ncbi.nlm.nih.gov/geo/query/acc.cgi?acc=GSE181915> | ChIP-Seq performed for exponentially growing culture in LB. Performed in triplicate. | Current study |
| *E. coli* DY330 EcTopoI ChIP-Seq with cells pre-treated with Rif | GSE181915 | <https://www.ncbi.nlm.nih.gov/geo/query/acc.cgi?acc=GSE181915> | ChIP-Seq performed for exponentially growing culture in LB. Culture was pre-treated with 100 mkg/ml rifampicine for 20 min before crosslinking. Performed in triplicate. | Current study |
| *E. coli* DY330 EcTopoI ChIP-Seq with cells expressing 14kDa TopoI CTD | GSE181915 | <https://www.ncbi.nlm.nih.gov/geo/query/acc.cgi?acc=GSE181915> | ChIP-Seq performed for exponentially growing culture in LB. CTD expression was induced when OD600 reached 0.2. Performed in triplicate. | Current study |
| *E. coli* DY330 EcTopoI ChIP-Seq with cells expressing 14kDa TopoI CTD and pre-treated with Rif | GSE181915 | <https://www.ncbi.nlm.nih.gov/geo/query/acc.cgi?acc=GSE181915> | ChIP-Seq performed for exponentially growing culture in LB. CTD expression was induced when OD600 reached 0.2. Culture was pre-treated with 100 mkg/ml rifampicine for 20 min before crosslinking. Performed in duplicate. | Current study |
| *E. coli* DY330 EcTopoI Topo-Seq | GSE182473 | <https://www.ncbi.nlm.nih.gov/geo/query/acc.cgi?acc=GSE182473> | Topo-Seq for EcTopoI cleavage sites mapping performed with exponentially growing culture in LB. Performed in triplicate. | Current study |
| *E. coli* DY330 DRIP-Seq | GSE181945 | <https://www.ncbi.nlm.nih.gov/geo/query/acc.cgi?acc=GSE181945> | DRIP-Seq for R-loops mapping performed with exponentially growing culture in LB. Performed in triplicate. | Current study |
| *E. coli* DY330 DRIP-Seq with cells pre-treated with Rif | GSE181945 | <https://www.ncbi.nlm.nih.gov/geo/query/acc.cgi?acc=GSE181945> | DRIP-Seq for R-loops mapping performed with exponentially growing culture in LB. Culture was pre-treated with 100 mkg/ml rifampicine for 20 min before cells harvesting. Performed in triplicate. | Current study |
| *E. coli* DY330 DRIP-Seq with cells expressing 14kDa TopoI CTD | GSE181945 | <https://www.ncbi.nlm.nih.gov/geo/query/acc.cgi?acc=GSE181945> | DRIP-Seq for R-loops mapping performed with exponentially growing culture in LB. CTD expression was induced when OD600 reached 0.2. Performed in triplicate. | Current study |
| *E. coli* DY330 DRIP-Seq with cells expressing 14kDa TopoI CTD and pre-treated with Rif | GSE181945 | <https://www.ncbi.nlm.nih.gov/geo/query/acc.cgi?acc=GSE181945> | DRIP-Seq for R-loops mapping performed with exponentially growing culture in LB. CTD expression was induced when OD600 reached 0.2. Culture was pre-treated with 100 mkg/ml rifampicine for 20 min before cells harvesting. Performed in triplicate. | Current study |
| WGS of *E. coli* BW25113, *E. coli* BW25113 *topAΔ11*, *E. coli* BW25113 *topAΔ14*, *E. coli* BW25113 *topAΔ30*, *E. coli* BW25113 *topAΔ11-SPA*, *E. coli* BW25113 *topAΔ14-SPA*, *E. coli* BW25113 *topAΔ30-SPA* | PRJNA757761 | <http://www.ncbi.nlm.nih.gov/bioproject/757761> | WGS of *E. coli* strains with truncated versions of topA gene constructed by lambda-red recombeneering. | Current study |
| *E. coli* DY330 DNA-gyrase Topo-Seq | GSM3273139, GSM3273141, GSM3273143 | https://www.ncbi.nlm.nih.gov/geo/query/acc.cgi?acc=GSE117186 | Topo-Seq performed for exponentially growing culture in LB. Ciprofloxacin was used to trap covalent intermediate complexes between gyrase and DNA. Performed in triplicate. | Sutormin D, Rubanova N, Logacheva M, Ghilarov D et al. Single-nucleotide-resolution mapping of DNA gyrase cleavage sites across the Escherichia coli genome. Nucleic Acids Res 2019 Feb 20;47(3):1373-1388. PMID: 30517674 |
| *E. coli* DY330 DNA-gyrase Topo-Seq for cells pre-treated with Rif | GSM3273145, GSM3273147, GSM3273149 | <https://www.ncbi.nlm.nih.gov/geo/query/acc.cgi?acc=GSE117186> | Topo-Seq performed for exponentially growing culture in LB. Culture was pre-treated with 100 mkg/ml rifampicine for 20 min before adding the ciprofloxacin to trap covalent intermediate complexes between gyrase and DNA. Performed in triplicate. | Sutormin D, Rubanova N, Logacheva M, Ghilarov D et al. Single-nucleotide-resolution mapping of DNA gyrase cleavage sites across the Escherichia coli genome. Nucleic Acids Res 2019 Feb 20;47(3):1373-1388. PMID: 30517674 |
| *E. coli* MG1655 RpoB ChIP-Seq | [GSM613808](https://www.ncbi.nlm.nih.gov/geo/query/acc.cgi?acc=GSM613808) | https://www.ncbi.nlm.nih.gov/geo/query/acc.cgi?acc=GSE24991 | ChIP-Seq performed for exponentially growing culture in LB. Single replicate. | Kahramanoglou C, Seshasayee AS, Prieto AI, Ibberson D et al. Direct and indirect effects of H-NS and Fis on global gene expression control in Escherichia coli. Nucleic Acids Res 2011 Mar;39(6):2073-91. PMID: 21097887 |
| *E. coli* MG1655 RpoC ChIP-chip for cells pre-treated with Rif | [GSM351003](https://www.ncbi.nlm.nih.gov/geo/query/acc.cgi?acc=GSM351003) | https://www.ncbi.nlm.nih.gov/geo/query/acc.cgi?acc=GSM351003 | ChIP-chip performed for exponentially growing culture in M9 medium. The culture was pre-treated with Rif before the experiment. | Mooney RA, Davis SE, Peters JM, Rowland JL et al. Regulator trafficking on bacterial transcription units in vivo. Mol Cell 2009 Jan 16;33(1):97-108. |
| *E. coli* MG1655 RpoS ChIP-exo | GSM1633277, GSM1633278 | https://www.ncbi.nlm.nih.gov/geo/query/acc.cgi?acc=GSE66441 | ChIP-exo performed for exponentially growing culture in M9 medium supplemented with 0.2% glucose and under acidic stress (pH=5.5). Performed in duplicate. | Seo SW, Kim D, O'Brien EJ, Szubin R et al. Decoding genome-wide GadEWX-transcriptional regulatory networks reveals multifaceted cellular responses to acid stress in Escherichia coli. Nat Commun 2015 Aug 10;6:7970. PMID: 26258987 |
| *E. coli* MG1655 RpoD ChIP-Seq | [GSM1010221, GSM1072326](https://www.ncbi.nlm.nih.gov/geo/query/acc.cgi?acc=GSM1010221) | https://www.ncbi.nlm.nih.gov/geo/query/acc.cgi?acc=GSE41195 | ChIP-Seq performed for exponentially growing aerobic culture. Performed in duplicate. | Myers KS, Yan H, Ong IM, Chung D et al. Genome-scale analysis of escherichia coli FNR reveals complex features of transcription factor binding. PLoS Genet 2013 Jun;9(6):e1003565. PMID: 23818864 |
| *E. coli* MG1655 GapR-Seq | GSM4628314, GSM4628313, GSM4628312, GSM4628311 | https://www.ncbi.nlm.nih.gov/geo/query/acc.cgi?acc=GSE152882 | ChIP-Seq performed for exponentially growing aerobic culture in LB. | Monica S Guo, Ryo Kawamura, Megan L Littlehale, John F Marko, Michael T Laub. High-resolution, genome-wide mapping of positive supercoiling in chromosomes. eLife 2021;10:e67236. PMID: 34279217 |
| *Mycobacterium tuberculosis* DNA-gyrase ChIP-Seq | [GSM2538162](https://www.ncbi.nlm.nih.gov/geo/query/acc.cgi?acc=GSM2538162) | https://www.ncbi.nlm.nih.gov/geo/query/acc.cgi?acc=GSE85357 | ChIP-Seq performed on exponentially growing culture of MtbRa cells. | Ahmed W, Sala C, Hegde SR, Jha RK et al. Transcription facilitated genome-wide recruitment of topoisomerase I and DNA gyrase. PLoS Genet 2017 May;13(5):e1006754. PMID: 28463980 |
| *Mycobacterium tuberculosis* RNAP ChIP-Seq | [GSM1003214, GSM1003215](https://www.ncbi.nlm.nih.gov/geo/query/acc.cgi?acc=GSM1003214) | https://www.ncbi.nlm.nih.gov/geo/query/acc.cgi?acc=GSE40862 | ChIP-Seq performed on exponentially growing culture. | Uplekar S, Rougemont J, Cole ST, Sala C. High-resolution transcriptome and genome-wide dynamics of RNA polymerase and NusA in Mycobacterium tuberculosis. Nucleic Acids Res 2013 Jan;41(2):961-77. PMID: 23222129 |
| *Mycobacterium tuberculosis* RNA-Seq | GSM1003224, GSM1003225 | https://www.ncbi.nlm.nih.gov/geo/query/acc.cgi?acc=GSE40846 | RNA-Seq performed for exponentially growing culture. | Uplekar S, Rougemont J, Cole ST, Sala C. High-resolution transcriptome and genome-wide dynamics of RNA polymerase and NusA in Mycobacterium tuberculosis. Nucleic Acids Res 2013 Jan;41(2):961-77. PMID: 23222129 |
| *Mycobacterium smegmatis* RNAP ChIP-Seq | [GSM1171544, GSM1171545](https://www.ncbi.nlm.nih.gov/geo/query/acc.cgi?acc=GSM1171544) | https://www.ncbi.nlm.nih.gov/geo/query/acc.cgi?acc=GSE48164 | ChIP-Seq performed on exponentially growing culture. | Landick R, Krek A, Glickman MS, Socci ND et al. Genome-Wide Mapping of the Distribution of CarD, RNAP σA, and RNAP β on the Mycobacterium smegmatis Chromosome using Chromatin Immunoprecipitation Sequencing. Genom Data 2014 Dec;2:110-113. PMID: 25089258 |
| *Mycobacterium smegmatis* TopoI ChIP-Seq | [SRX4970107](https://www.ncbi.nlm.nih.gov/sra/SRX4970107%5baccn%5d) | https://www.ncbi.nlm.nih.gov/bioproject?LinkName=biosample_bioproject&from_uid=10371555 | ChIP-Seq performed on exponentially growing culture. | Rani P et al., "Genome-wide mapping of Topoisomerase I activity sites reveal its role in chromosome segregation.", Nucleic Acids Res, 2019 Feb 20;47(3):1416-1427 |
| *Mycobacterium smegmatis* Mock DNA for ChIP-Seq normalization | [GSM4274349, GSM4274350](https://www.ncbi.nlm.nih.gov/geo/query/acc.cgi?acc=GSM4274349) | https://www.ncbi.nlm.nih.gov/geo/query/acc.cgi?acc=GSE143764 | Mock DNA data | Feng S, Liu Y, Liang W, El-Sayed Ahmed MAE et al. Involvement of Transcription Elongation Factor GreA in Mycobacterium Viability, Antibiotic Susceptibility, and Intracellular Fitness. Front Microbiol 2020;11:413. |
| *Mycobacterium smegmatis* RNA-Seq | [GSM2756262, GSM2756263](https://www.ncbi.nlm.nih.gov/geo/query/acc.cgi?acc=GSM2756262) | https://www.ncbi.nlm.nih.gov/geo/query/acc.cgi?acc=GSE103158 | RNA-Seq performed for exponentially growing culture. | Li X, Mei H, Chen F, Tang Q et al. Transcriptome Landscape of Mycobacterium smegmatis. Front Microbiol 2017;8:2505. PMID: 29326668 |
| *Streptococcus pneumoniae* TopoI and RNAP ChIP-Seq | [SRR12427932, SRR12427936, SRR12427942](https://trace.ncbi.nlm.nih.gov/Traces/sra/?run=SRR12427932) | https://www.ncbi.nlm.nih.gov/bioproject/PRJNA656439 | ChIP-Seq performed for exponentially growing culture. | Maria-Jose Ferrandiz, Pablo Hernandez, Adela G. de la Campa, Genome-wide proximity between RNA polymerase and DNA topoisomerase I supports transcription in Streptococcus pneumoniae. PLOS Genetics 2021. PMID: 33930020 |
