## Supplementary material for "Interaction Between Transcribing RNA Polymerase and Topoisomerase I Prevents R-loop Formation in *E. coli*": Supplemetary Materials & Methods

***E. coli* TopoI ChIP-Seq and DRIP-Seq**

**Purification of IP-DNA with AMPure XP magnetic beads**

To one volume of DNA solution one volume of binding buffer was added (10 mM Tris-HCl pH=8, 1 mM EDTA, 250 mM NaCl, 20% PEG-8000, 0.05% Tween-20) and one volume of AMPure XP beads. Suspension was incubated for 10 min at room temperature. Magnetic beads were separated on a magnetic stand and supernatant was discarded. Beads’ pellet was carefully washed 3 times with 75% ethanol on a magnetic stand. Beads were dried and resuspended in 15 µL of deionized water. Beads were trapped on a magnetic stand and supernatant was collected and stored at -20°C.

***E. coli* RNAP ChIP-Seq**

For TAP-tagged RNAP-ChIP, we used *E. coli* DY330 strain (Dharmacon) carrying chromosomally TAP-tagged *rpoC* gene (Butland *et al.*, 2005). Cell culture (V=200 mL) grown to mid-exponential phase OD600~0.5-0.7 in LB containing 50 ug/mL of kanamycin was crosslinked by adding formaldehyde to a final concentration of 1%, followed by incubation for 20 min at room temperature, with agitation. Crosslinking was stopped by the addition of sterile glycine to a final concentration of 0.25M. Cells were incubated for 20 min at room temperature and harvested by centrifugation at 3000 x g, 10min, 4°C. Cell pellets were washed twice with 20 mL of ice-cold 20 mM Tris-HCl buffer pH 7.6, containing 60 mM NaCl (Tris-buffered saline, TBS), resuspended in 0.5 mL of 2x ChIP Lysis Buffer 1 (10 mM tris pH 8.0, 50 mM NaCl, 10 mM EDTA, 20% Sucrose), followed by addition of 0.5 mL of 2x ChIP Lysis Buffer 2 (200 mM Tris pH 8.0, 600 mM NaCl, 4% Triton X-100), containing RNaseA (1 µg/mL). Resuspended cells were pre-lysed by the addition of 1 ug of Lysozyme and incubation at 37°C for 10 min, followed by chilling on ice. Next, cells were sonicated in a 2.0 mL Eppendorf tube in an ice-water bath, using microtip to achieve a high yield of 300-400bp fragments. Lysates were clarified by centrifugation at 13,000 x g for 10 min at 4°C, and the resulting supernatant was used for further analysis.

For the Input DNA fragment size analysis, a 100 µL aliquot of the clarified lysate was first treated with proteinase K (Thermo-Fisher Scientific, 25530049) at 0.125 mg/mL in the presence of 1% SDS for 4 h at 37°C followed by DNA de-crosslinking by incubation at 65°C for 6 h. The Input DNA material was purified using Qiagen PCR purification kit (Qiagen, 28104) and quantified by Nano-Drop spectrophotometer. The DNA fragment size distribution (in 30-60 ng aliquots) was assessed by 8% DNA PAGE using Low Molecular Weight DNA markers (New England Biolabs, N3233S), visualized by SYBR-gold (Thermo fisher scientific, S11494) staining and quantified by Kodak Imaging station.

For the preparation of RNAP-ChIP DNA, the remaining lysate (~900 ul) was purified by immunoprecipitation (IP). The lysate was mixed with 10 µL of IgG-agarose (GE Healthcare) and incubated at 4°C overnight on a rotating mixer in the presence of Protease Inhibitor Cocktail (Sigma, s8830).

Then affinity resin was washed consecutively with 1 mL each of the following solutions: 40 mM Tris pH 7.9, 0.5% Tween 20, 2 M NaCl; 40 mM Tris pH 7.9, 0.5% Tween 20, 1 M NaCl; 40 mM Tris pH 7.9, 0.5% Tween 20, 200 mM NaCl; and twice with RIPA buffer (50 mM Tris pH 7.4, 140 mM NaCl, 1% NP-40, 0.1% Deoxycholate, 0.1% SDS). All wash procedures were carried out by resuspending the beads in the wash solution and inverting the tube several times, followed by brief centrifugation and removing the supernatant by vacuum aspiration. The last wash was done for 20 min at 4°C on a rotary mixer. The RNAP-DNA crosslinks were eluted by incubation with ChIP Elution Buffer (10 mM Tris pH 8.0, 30 mM EDTA, 1% SDS) at 65°C for 4 hours (or O/N) on a shaker at 800-900 rpm. The resulting material was treated with Proteinase K (Thermo Fisher Scientific, 25530049) (0.2 mg/mL) at 37°C for 3-5h, followed by IP-DNA decrosslinking by incubation at 95°C for 2h. The IP-DNA was finally purified using ChIP DNA Cleaning & Concentrator kit (Zymo Research, D5205).

A 50-100 ng of IP-DNA or Input-DNA was subjected to the following enzymatic and purification steps:

1. *End repair*

Reactions were performed in the final volume of 50 µL in the presence of a 1x T4 DNA ligase buffer (NEB, B0202s), 0.4 mM dNTP mix, 3 units of T4 DNA polymerase (NEB, M0203s), 10 units of T4 polynucleotide kinase (PNK) (NEB, M0201s) and 5 units of Klenow DNA polymerase (NEB, M0210s). Reactions were incubated for 30 min at 20°C, and DNA was purified by Qiaquick PCR DNA purification kit (Qiagen, 28104) using 36 µL of Qiagen Elution Buffer (EB).

2. *Addition of “A” bases to the 3’-end of the DNA fragments*

The eluted DNA material from the previous step was combined with 0.2 mM dATP and 5 units of Klenow Fragment (3ʼ to 5ʼ exo minus) (NEB, M0212s) in 50 µL of Klenow buffer (NEB buffer 2). Reactions were incubated for 30 min at 37°C, and DNA was purified by MinElute PCR purification Kit (Qiagen, 28004) using 15 µL of EB.

3. *Adapter ligation.*

Multiplex Adapter (MPA) mix was prepared by combining 1 umole of Illumina DNA adapters MP_Adapt1 and MP_Adapt2 with 1x NEB Buffer 2 in the final volume of 50 µL followed by incubation in a thermal cycler at 95°C for 5min, and then 80°C, 70°C, 60°C, 50°C, 40°C, and 30°C for 1 min each step. The MPA mix was diluted 1:20 in EB, to a final concentration of ~2 µM (needed for ligation). The DNA material from the previous step was used in a ligation reaction with 2 pmoles of diluted MPA oligo mix and 2000 units of the Quick DNA ligase (NEB, M2200s) in 30 µL of Quick DNA ligase Buffer (NEB, M2200s). Reactions were incubated for 15 min at 20°C, and DNA was purified by Agencourt Ampure XP beads (Beckman Coulter, A63880) using 11 µL of elution buffer.

4. *DNA size selection using 2% agarose gel electrophoresis*

Samples were separated on a 2% Low Range Ultra-Agarose gel (BioRad, 161-3106) in standard Tris-acetate-EDTA buffer (TEA) using Low Molecular Weight Ladder and stained by SYBR-gold DNA dye. DNA bands corresponding to 220 bp size were excised by Gel X-tracta tool (USA scientific, 5454-2500) and purified with Gel Extraction Kit (Qiagen, 28704) using 38 µL of EB.

5. *PCR enrichment of the Adapter-Modified DNA fragments*

The purified DNA material was PCR-amplified using 2 uM PCR primer 1_1, 2 uM PCR primer 2_1 and 1 unit of Phusion polymerase (NEB, M0530s) in the presence 0.3mM dNTP in 1x Phusion HF buffer, under the following conditions: first cycle - 30 seconds at 98°C, followed by 18 cycles with 30 sec at 65°C and 30 sec at 72°C, completed by the final step for 5 minutes at 72°C. The resulting PCR-amplified DNA was purified by MinElute PCR purification kit (Qiagen, 28004) using 16 µL of EB.

Input and IP DNA samples were analyzed by Nano-Drop spectrophotometer and 8% DNA-PAGE with SYBR-gold (Thermo Fisher scientific, S11494) staining quantified by Kodak Gel Logic 200 Imager. The final library DNA concentrations were analyzed by qPCR using KAPA Illumina library quantification kit (Kapa Biosystems, KK4824).

DNA sequencing was performed by Illumina Hi-Seq 50 bp paired-end conditions using 24 TruSeq Index Primers (Illumina).

Bioinformatics analysis of ChIP-Seq data was performed by Galaxy Project Open source cloud-based software (Afgan *et al.*, 2016) and R-studio using appropriate Bioconductor R-packages (https://www.bioconductor.org/). First, de-multiplexed and de-barcoded, QC-filtered short-read sequences were aligned to the *E. coli* W3110 MuSGS genome based on NC_007779.1 sequence (https://github.com/sutormin94/TopoA_ChIP-Seq/blob/master/Additional_genome_features/E_coli_w3110_G_Mu.fasta) using Bowtie for Illumina short read aligner tool (Langmead *et al.*, 2009). Mapping statistics, read depth, and ChIP signal quality assessment for all samples, as well as reproducibility between biological replicas, was calculated using DeepTools (Ramírez *et al.*, 2014). Log2 ratio profiles of IP to input samples were calculated using BAMcompare tool. To compare the BAM files to each other, the reference genome was partitioned into bins of equal size (10000bp), then the number of reads found in each BAM file was counted per bin, and finally, a summary value was reported as the log2 ratio.

Reproducibility of ChIP biological replicas was assessed by calculating Pearson correlations on mapped reads and log2 ratio profiles of IP samples normalized to input DNA using multiBigWigSummary and plotCorrelationImage tools. Peak calling, signal normalization, and PCR duplicate removal were performed using MACS2 (Zhang *et al.*, 2008). Fold enrichment of RNAP ChIP signal over Input signal was performed using bdgcmp (“BedGraph Compare”) function of MACS2 using Fold Enrichment (FE) method.  The resultant signal tracks represented normalized binding profiles were visualized using Integrated Genome Viewer (IGV) (Thorvaldsdóttir, Robinson and Mesirov, 2013). Fold enrichment track was used for further analysis using custom python scripts (<https://github.com/sutormin94/TopoA_ChIP-Seq>).

***Mycobacterium tuberculosis* ChIP-Seq (MtbRNAP, MtbGyrase) and RNA-Seq data analysis**

Expression level of *M. tuberculosis* TUs in a log-phase was taken from Uplekar et al., 2013 (Supplementary material, Table 7) (Uplekar *et al.*, 2013). Top 200 of 606 TUs by expression level comprise HETU set (highly-expressed TUs, FPKM > 4.75), similarly 606 TUs with the lowest expression level comprise LETU set (low-expressed TUs, FPKM < 1.9).

Publicly available RNAP β-subunit ChIP-Seq datasets for exponentially growing culture of *M. tuberculosis* H37Rv were taken from GEO (GSM1003214, GSM1003215) (Uplekar *et al.*, 2013). Raw reads were trimmed and filtered using Trimmomatic and aligned to the *M. tuberculosis* H37Rv genome (NC_000962.3) using BWA-mem (Li and Durbin, 2010). PCR duplicates were not removed, because the dataset comprised of single-stranded reads. Coverage depth bed track was obtained using samtools and converted to wig format with a custom script (**MtbRNAP IP tracks**). Mock control datasets from the same study (GSM1003222, GSM1003223) were processed identically (**Mock tracks**). Fold enrichment tracks for RNAP β-subunit of *M. tuberculosis* were obtained as a position-wise ratio of normalized **IP tracks** and corresponding **Mock tracks**. Fold enrichment tracks of biological replicates (GSM1003214/GSM1003222, GSM1003215/GSM1003223) were position-wise averaged. Regions of a genome having fold enrichment > 3 were defined as MtbRNAP peaks. Totally, 2464 peaks were identified.

Publicly available DNA-gyrase ChIP-Seq dataset for exponentially growing culture of *M. tuberculosis* H37Rv was taken from GEO (GSM2538162) (Ahmed *et al.*, 2017). Raw reads were trimmed and filtered using Trimmomatic and aligned to the *M. tuberculosis* H37Rv genome (NC_000962.3) using BWA-mem (Li and Durbin, 2010). PCR duplicates were removed, using samtools. Coverage depth bed track was obtained using samtools and converted to wig format with a custom script (**MtbGyrase IP track**). Mock control dataset from the same study (GSM2538163) was processed identically (**Mock track**). Fold enrichment track for DNA-gyrase of *M. tuberculosis* was obtained as a position-wise ratio of normalized **IP track** and a **Mock track**. Regions of a genome having fold enrichment > 2 were defined as MtbGyrase peaks. Totally, 286 peaks were identified.

***Mycobacterium smegmatis* ChIP-Seq (MsmRNAP, MsmTopoI) and RNA-Seq data analysis**

Publicly available RNA-Seq datasets for mid-exponential culture of *M. smegmatis* were taken from GEO (GSM2756262, GSM2756263) for estimation of the expression level of TUs (Li *et al.*, 2017). Raw reads were trimmed and filtered using Trimmomatic and aligned to the *M. smegmatis MC^2^-155* genome (CP009496.1) using BWA-mem (Li and Durbin, 2010). PCR-duplicates were removed with samtools (Li *et al.*, 2009). Genome annotation and TUs coordinates was taken from BioCyc (Karp *et al.*, 2018) for genome assembly NC_008596.1 and transferred for CP009496.1 using blast. Expression level of TUs was calculated as FPKM value using FPKM_count.py from RSeQC package (Wang, Wang and Li, 2012). Final TU FPKM was calculated as a mean of two biological replicates (GSM2756262, GSM2756263). Top 200 of 4098 TUs by expression level comprise HETU set (highly-expressed TUs, FPKM > 634.8), similarly 200 TUs with the lowest expression level comprise LETU set (low-expressed TUs, FPKM < 3.2).

Publicly available RNAP β-subunit ChIP-Seq datasets for exponentially growing culture of *M. smegmatis* were taken from GEO (GSM1171544, GSM1171545) (Landick *et al.*, 2014). Raw reads were trimmed and filtered using Trimmomatic and aligned to the *M. smegmatis MC^2^-155* genome (CP009496.1) using BWA-mem (Li and Durbin, 2010). PCR duplicates were not removed, because the datasets comprised of single-stranded reads. Coverage depth bed tracks were obtained using samtools and converted to wig format with a custom script. Tracks of biological replicates (GSM1171544, GSM1171545) were position-wise averaged (**MsmRNAP** **IP track**).

Mock control datasets for *M. smegmatis* ChIP-Seq experiments were taken from GEO (GSM4274349, GSM4274350) (Feng *et al.*, 2020). Raw reads were trimmed and filtered using Trimmomatic and aligned to the *M. smegmatis MC^2^-155* genome (CP009496.1) using BWA-mem (Li and Durbin, 2010). PCR-duplicates were removed with samtools (Li *et al.*, 2009). Coverage depth bed tracks were obtained using samtools and converted to wig with custom script. Tracks of biological replicates (GSM4274349, GSM4274350) were position-wise averaged (**Mock track**).

Fold enrichment track for RNAP β-subunit of *M. smegmatis* was obtained as a position-wise ratio of normalized **MsmRNAP** **IP track** and **Mock track**. Regions of a genome having fold enrichment > 4 were defined as MsmRNAP peaks. Totally, 2623 peaks were identified.

Publicly available TopoI ChIP-Seq dataset for exponentially growing culture of *M. smegmatis* was taken from SRA (BioSample SAMN10371555, SRA ID [SRR8149480](https://trace.ncbi.nlm.nih.gov/Traces/sra/?run=SRR8149480)) (Rani and Nagaraja, 2018). Raw reads were trimmed and filtered using Trimmomatic and aligned to the *M. smegmatis MC^2^-155* genome (CP009496.1) using BWA-mem (Li and Durbin, 2010). PCR duplicates were removed with samtools (Li *et al.*, 2009). Coverage depth bed track was obtained using samtools and converted to wig with custom script (**MsmTopoI** **IP track**). Fold enrichment track for TopoI of *M. smegmatis* was obtained as a position-wise ratio of normalized **MsmTopoI** **IP track** and **Mock track**. Regions of a genome having fold enrichment > 3 were defined as MsmTopoI peaks. Totally, 153 peaks were identified.

***E. coli* GapR-Seq data analysis**

Publicly available GapR-Seq datasets for *E. coli* MG1655 in exponential phase were taken from GEO (GSM4628314, GSM4628313, GSM4628312, GSM4628311) for mapping of positive and negative supercoiling over the *E. coli* genome (Guo *et al.*, 2021). Raw reads were trimmed and filtered using Trimmomatic and aligned to the *E. coli* W3110 MuSGS genome (*E. coli* W3110 genome with the insertion of *cat*-Mu SGS cassette may be downloaded from GEO: GSE95567) using BWA-MEM (Li and Durbin, 2010). BAM and bed files were prepared with Samtools (Li *et al.*, 2009) and visualized in IGV (Thorvaldsdóttir, Robinson and Mesirov, 2013). Biological replicates for MG1655 pKVS45-GapR experiments (GSM4628312, GSM4628311) and MG1655 pKVS45-GapR-3xFLAG experiments (GSM4628314, GSM4628313) were scaled by the total amount of mapped reads and averaged separately. Fold enrichment tracks were prepared by position-wise division of GapR-FLAG resultant track by GapR track (see <https://github.com/sutormin94/TopoA_ChIP-Seq>).

***E. coli* RpoC ChIP-chip data analysis**

Publicly available RpoC ChIP-chip datasets for *E. coli* MG1655 treated with RNAP inhibitor rifampicin in exponential phase were taken from GEO (GSM351003) for mapping of RNAP enrichment over the *E. coli* genome (Mooney *et al.*, 2009). The coordinates of the microarray oligonucleotides were identified in reference *E. coli* W3110 MuSGS genome (*E. coli* W3110 genome with the insertion of *cat*-Mu SGS cassette may be downloaded from GEO: GSE95567) using blast. For each genomic position a fold enrichment was calculated as an average enrichment of oligonucleotides aligned to this position (https://github.com/sutormin94/TopoA_ChIP-Seq/blob/master/Convert_chip_oligos_to_coordinates_Mooney_data.py).

**Meta-gene analysis**

To produce scaled meta-gene plots, fold enrichment of protein of interest or other signal was extracted in vicinity of transcription units (15kb upstream, TU body, 15kb downstream) in concordance to their orientation. Regions were scaled to have the same number of positions (5000bp) by omitting of randomly chosen points (if region is longer than 5000) or by random duplication of points (if region is shorter than 5000). Data extraction and scaling was performed with *FE_over_US_GB_DS.py* custom script. Resulting scaled arrays were averaged by position, smoothed with averaging sliding window 200bp and plotted with *Plot_signal_over_transcription_units.py* custom script. Zoom-in meta-gene plots representing the proximity of transcription start and transcription and sites (+300bp:-200bp and -200bp:+300bp respectively) were produced without smoothing.

Unscaled meta-gene plots for transcription units (TU) were produced as following. Fold enrichment of protein of interest was extracted for transcription units in concordance to their orientation. Resulting arrays were averaged by position. If some region was shorter than position examined, it was omitted. Additionally, for each position mean standard error of fold enrichment was calculated. EcTopoI fold enrichment typically has a maximum at transcription start site (TSS) and is going down toward the transcription end site (TES). The analysis was performed with *Compare_signal_US_TSS_GB.py* custom script. All scripts mentioned are available from the <https://github.com/sutormin94/TopoA_ChIP-Seq> repository.

DRIP-Seq data was analyzed similarly, but in respect to the data strand-specificity, the signals for TUs in a reverse orientation were multiplied by -1 to flip them and make them concordant with signals for TUs in a forward orientation (see *FE_over_US_GB_DS_str_spec.py* script in the <https://github.com/sutormin94/E_coli_DRIP-Seq_analysis> repository). For Topo-Seq data, in respect to the data strands-specificity, signals for coding and template strands of TUs in a forward orientation were taken from N3E_F and N3E_R tracks respectively; signals for coding and template strands of TUs in a reverse orientation were taken from N3E_R and N3E_F tracks respectively (see *FE_over_US_GB_DS_strand_specific_binning_and_statistics.py* script in the <https://github.com/sutormin94/TopoI_Topo-Seq> repository)

**Identification of EcTopoI cleavage sites (TCSs)**

Separately for forward and reverse strands, the number of DNA fragments 3’-ends was calculated per position (N3E) based on read alignments stored in SAM files (giving N3E_F and N3E_R tracks respectively). The tracks were scaled by the total number of aligned reads to get normalized coverage across samples and biological replicates were averaged. After that -IP tracks (+Ara-IP and -Ara-IP) were subtracted from corresponding +IP tracks (+Ara+IP and -Ara+IP respectively) strand-wise resulting in +Ara and -Ara tracks. TCSs were detected as sites having a signal higher than threshold value 15 (values 10 and 20 were also tested and given similar results in respect to identified DNA motif).

**Purification of EcTopoI**

15 mL of a night culture of *E. coli* BL21(DE3) pET28 topA_strep in LB, supplemented with kanamycin 50 µg/ml, was added to 1.5 L of LB medium and cultivated at 37^o^C and 180 rpm. At OD_600_~0.6 the culture was induced with IPTG (final concentration 0.4 mM) and cultivation continued for 4 h. The cells were pelleted (5000 rpm, 10 min, 4^o^C) and frozen in liquid nitrogen. Subsequently pellet was thawed on ice and resuspended in 5 mL of Lysis Buffer (100 mM Tris HCl pH 8.0, 150 mM NaCl, 1 mM EDTA) supplemented with 1 mM PMSF, lysed using incubation with 2 mg/mL lysozyme (1 h, 4^o^C) followed by sonication for 10 min. The obtained lysate was cleared by centrifugation (14000 rpm, 20 min, 4^o^C). Strep-tagged EcTopoI was purified on StrepTrap HP column (GE Healthcare) using protocol recommended by manufacturer. Briefly, the lysate was loaded on the 5 mL column pre-equilibrated with Lysis Buffer and the column was washed with 50 mL of Lysis buffer. Then proteins bound were eluted using 10 mL of Elution Buffer (50 mM Tris HCl, pH 8.0, 150 mM NaCl, 1 mM EDTA, 2.5 mM desthiobiotin). The eluted proteins were analyzed using electrophoresis in 11% PAA gel, the bands were visualized with Instant Blue protein stain and subjected to protein identification via MS. 10 mL of eluate was dialyzed overnight at 4^o^C against 1.5 l of storage buffer (10 mM Tris-HCl pH 7.5, 50 mM KCl, 0.1 mM EDTA, 1 mM 2-mercaptoethanol) and then concentrated 5x on Amicon Ultra-15 filter (Millipore) by centrifugation (3500 g, 20 min, 4^o^C). Glycerol was added to 30% w/v and sample was stored at -20^o^C. Activity of the enzyme was confirmed by relaxation assay with a negatively supercoiled plasmid substrate.

**Relaxation assay with EcTopoI**

2 µL (330 ng/µL) of negatively supercoiled pMP1000 plasmid extracted from *E. coli* DH5α cells was combined with 2 µL of 10x Reaction buffer (100 mM Tris-HCl pH 8, 500 mM NaCl, 60 mM MgCl_2_), different amount of EcTopoI (0, 50, 100, 200, 300, 400 ng of the enzyme with concentration 170 ng/µL) and milli-Q H_2_O up to 20 µL. The mixture was incubated 30 min at 37^o^C and reaction was stopped by addition of 5 µL of 5x Stop-buffer (50 mM EDTA pH 8, 0.5% bromophenol blue, 50% glycerol). Relaxation products were separated by electrophoresis in 1% agarose gel supplemented with 2 µg/mL chloroquine. After separation, for DNA visualization gel was stained with ethidium bromide.

**Protein affinity purification**

To demonstrate that the expression of the EcTopoI 14kDa CTD fragment was induced in the samples used for physiological experiments upon the addition of IPTG, we performed a small-scale affinity purification of N-terminally 6His-tagged protein from cell lysate with subsequent SDS-PAGE analysis and MS-analysis.

Cell culture (100 mL) of *E. coli* DY330 *topA*-SPA pCA24 14kDa CTD was started by the addition of 1 mL of the corresponding overnight culture to fresh LB medium supplemented with 50 µg/mL kanamycin, 34 µg/mL chloramphenicol, and 0.5% glucose. After reaching the OD_600_~0.2 the culture was bisected and one half of it was induced with 1 mM IPTG, the other was used as a control. The cells were pelleted (5000 rpm, 10 min, 4^o^C) after 1 h of cultivation and frozen in liquid N_2_. Subsequently each cell pellet was thawed on ice and resuspended in 1.5 mL of Lysis Buffer (50 mM Tris HCl pH 8.0, 150 mM NaCl, 2.5 mM Imidazole), lysed using incubation with 2 mg/mL lysozyme (1 h, 4^o^C) followed by sonication for 10 min. The obtained lysate was cleared by centrifugation (14000 rpm, 20 min, 4^o^C) and added to 30 μl of TALON superflow affinity resin (Sigma) pre-equilibrated with Lysis Buffer. After 1 h of incubation at 4^o^C with constant rotation the resin was washed five times with Lysis Buffer, the proteins bound were eluted using 40 μl of Elution Buffer (50 mM Tris HCl pH 8.0, 150 mM NaCl, 300 mM Imidazole). The eluted proteins were analyzed using electrophoresis in 11% PAA gel, the bands were visualized with Instant Blue protein stain and subjected to protein identification via MS.

**Protein identification by MS**

Small pieces of the bands of interest were cut out from the gel, washed from the stain twice with 60 mM (NH_4_)_2_CO_3_ in 40% acetonitrile and dehydrated with 100% acetonitrile. After the addition of the digestion mixture containing 50 mM (NH_4_)_2_CO_3_ and 15 ng/μL sequencing grade trypsin the samples were incubated overnight at 37ᵒC. The peptides were extracted using 10 μl of 0.5% trifluoroacetic acid. 2-3 μl of the obtained sample were subjected to MS analysis on UltrafleXtreme MALDI-TOF-TOF mass spectrometer (Bruker Daltonics) to get the peptide mass fingerprint. NCBI protein database search with Mascot software (Matrix Science) was performed for protein identification.

**Pull-down of proteins bound to EcTopoI**

An isolated colony of *E. coli* DY330 *topA-SPA* was inoculated in 5 mL of LB and cultivated overnight at 37^o^C with shaking 180 rpm. Next day the starter was diluted 100x with fresh 2xYT to get 200 mL of culture and cultivated at 37^o^C until reaching OD_600_~0.6-0.7. Then the culture was cooled on ice and centrifuged at 4^o^C, 5000 rpm for 15 min. Cells pellet was frozen in liquid N_2_ and stored at -80^o^C. Then the pellet was dissolved in 3 mL of PBS with addition of Proteoblock cOmplete and lysozyme (2 mg/mL) and incubated for 1 h at 25^o^C. Cells were gently lysed by 2 repetitive cycles of freeze-thawing and then sonicated (5s pulse/5s relaxation for 2 min, 50% power, SONOPULS HD 3100). Cells debris was removed by centrifugation at 4^o^C, 15000 rpm for 15 min and cleared lysate was incubated with 100 µL of ANTI-FLAG® M2 affinity gel (Sigma-Aldrich) at 4^o^C for 1 h with constant rotation. Affinity resin was washed 4 times with 1 mL of PBS and bound proteins were eluted by 10 min incubation at 95^o^C with 30 µL of 3x Laemmli buffer supplemented with 2-mercaptoethanol. Proteins were separated in 10% PAAG and were identified by MS.

Pull-down experiments with *E. coli* DY330 *topA-SPA* transformed with a plasmid (pCA24 GFP or pCA24 14kDa CTD) were performed essentially as described above. The starter culture was supplemented with 0.5% glucose and 34 µg/mL chloramphenicol and grown overnight. Next the starter was diluted 100x with 2xYT supplemented with glucose and antibiotic to get 400 mL of culture. The culture was cultivated until OD_600_~0.2 and then bisected. One half of it was induced with 1 mM IPTG and the cultures were cultivated for 1 h until OD_600_~0.6-0.7.

**Pull-down of proteins bound to EcTopoI 14kDa CTD and analysis by Western-blot**

Cell cultures (100 mL) of *E. coli* DY330 *topA*-SPA pCA24 14kDa CTD or *E. coli* DY330 *rpoC*-TAP pCA24 14kDa CTD were started by the addition of 1 mL of the corresponding overnight culture to fresh LB medium supplemented with 50 µg/mL kanamycin, 34 µg/mL chloramphenicol, and 0.5% glucose. After reaching the OD_600_~0.2 the culture was bisected and one half of it was induced with 1 mM IPTG, the other was used as a control. The cells were pelleted (5000 rpm, 10 min, 4^o^C) after 1 h of cultivation and frozen in liquid N_2_. Subsequently each cell pellet was thawed on ice and resuspended in 1.5 mL of Lysis Buffer (50 mM Tris HCl pH 8.0, 150 mM NaCl, 5 mM Imidazole) with addition of Proteoblock cOmplete, and lysed by sonication for 10 min. Total concentration of proteins in the obtained lysates was measured by Bradford assay (Protein Assay Dye Reagent Concentrate, Bio-Rad). Equal amounts of lysates (by total protein amount) were added to 80 μl aliquots of TALON superflow affinity resin (Sigma) pre-equilibrated with Lysis Buffer. After over-night incubation at 4^o^C with constant rotation the resin was washed five times with 1 mL of Lysis Buffer, the proteins bound were eluted using incubation with 30 μl of Elution Buffer (50 mM Tris HCl pH 8.0, 150 mM NaCl, 300 mM Imidazole) for 15 min at room temperature. Not-bound fractions and the eluted proteins were analyzed using electrophoresis in 10% PAA gel, the bands were visualized with Instant Blue protein stain and subjected to protein identification via MS.

To specifically detect SPA-tagged EcTopoI (for *E. coli* DY330 *topA*-SPA pCA24 14kDa CTD) and TAP-tagged RpoC (for *E. coli* DY330 *rpoC*-TAP pCA24 14kDa CTD) among proteins co-purified with EcTopoI 14kDa CTD a Western blot was performed.

Not-bound fractions and the eluted proteins were analyzed using electrophoresis in 10% PAA gel and proteins were transferred to Hybond-P membrane (Amersham) using Trans-Blot SD Semi-dry transfer cell (Bio-Rad). Then the membrane was blocked with 25 mL of TBST buffer (20 mM Tris-HCl pH 7.5, 150 mM NaCl, 0.1% Tween 20) + 5% milk for 30 min with agitation 50 rpm on an orbital shaker. The incubation and all the subsequent incubations were performed at room temperature. Then membrane was washed 3 times for 5 min with 25 mL TBST. Primary anti-FLAG antibodies produced in rabbit (Sigma Aldrich, F7425) were added 1:10000 in 25 mL of TBST and hybridized with the membrane for 1.5 h. Unbound antibodies were washed out with 25 mL TBST (3 times, 5 min each). Secondary anti-rabbit antibodies produced in goat conjugated with HRP (Sigma Aldrich, A0545) were added 1:25000 to 25 mL of TBST + 5% milk and hybridized with the membrane for 1 h. Not bound antibodies were washed out with 25 mL of TBST (3 times, 5 min each). Signal of HRP was detected using Clarity Western ECL Substrate (Bio-Rad) and captured with Fusion Solo S detection system (Vilber Lourmat).

**Dot-blot with S9.6 antibodies against RNA:DNA hybrids**

Single colony of *E. coli* DY330 *topA-SPA* harboring pCA24 14kDa CTD was inoculated into 5 mL of LB supplemented with chloramphenicol 34 µg/mL and 0.5% glucose and grown overnight. The starter (1 mL) was diluted 100 times with 100 mL of LB containing the antibiotic and 0.5% glucose and cultivated until OD_600_~0.2 at 37^o^C with shacking, then the culture was bisected, and one half was induced with 1mM IPTG and cultivated for 1 h. Cells were harvested by centrifugation and total nucleic acids was purified using GeneJET Genomic DNA purification kit (ThermoFisher) according to the manufacturer protocol but omitting the RNAse A treatment. Aliquots containing 100 ng of DNA (concentration measured by Qubit using dsDNA HS assay kit – Invitrogen) were treated with 1U of RNAse III (Ambion) or with 1U of RNAse III and 5U of RNAse HI (NEB) for 2h at 37^o^C in RNAse III reaction buffer (10 mM Tris-HCl pH 7.9, 50 mM NaCl, 10 mM MgCl_2_, 1 mM DTT) and nucleic acids were purified with GeneJET Gel Extraction and DNA cleanup micro kit (General cleanup protocol, ThermoFisher). Equal amounts of samples (RNAse III or RNAseIII/RNAse HI treated) equivalent to 50 ng of DNA (concentration measured by Qubit using dsDNA HS assay kit – Invitrogen) were applied onto Hybond-N+ membrane (Amersham) and air dried. Membrane was crosslinked with UV (70000 µJoules/cm^2^) using HL2000 Hybrylinker. Then the membrane was blocked with 25 mL of TBST buffer (20 mM Tris-HCl pH 7.5, 150 mM NaCl, 0.1% Tween 20) + 5% milk for 30 min with agitation 50 rpm on an orbital shaker. The incubation and all the subsequent incubations were performed at room temperature. Then membrane was washed 3 times for 5 min with 25 mL TBST. 15 µg of primary S9.6 antibodies produced in mice (Kerafast, ENH001) were hybridized with the membrane in 25 mL of TBST for 1.5 h. Unbound antibodies were washed out with 25 mL TBST (3 times, 5 min each). Secondary anti-mouse antibodies conjugated with HRP (Sigma Aldrich, A9044) were added 1:80000 to 25 mL of TBST + 5% milk and hybridized with the membrane for 1 h. Not bound antibodies were washed out with 25 mL of TBST (3 times, 5 min each). Signal of HRP was detected using Clarity Western ECL Substrate (Bio-Rad) and captured with Fusion Solo S detection system (Vilber Lourmat).

**Probing the transcript coverage in response to CTD overexpression**

Induced and non-induced *E. coli* DY330 pCA24 14kDa CTD cultures were cultivated as described for ChIP-Seq (see ***EcTopoI ChIP-Seq***). RNA was extracted from 2 mL of culture using ExtractRNA reagent (Evrogen). RNA sample (90 µg) was treated with 5 U of DNAse I (ThermoFisher Scientific) for 30 min at 37^o^C and RNA was purified with RNAClean XP beads (Beckman Coulter). cDNA was synthesized from 7 µg of RNA using Maxima Reverse Transcriptase with random hexamer primers (Thermo Fisher Scientific). qPCR was performed using Maxima SYBR green qPCR master mix (Thermo Fisher Scientific) on CFX96 Real-Time System (Bio-Rad).

Afgan, E. *et al.* (2016) ‘The Galaxy platform for accessible, reproducible and collaborative biomedical analyses: 2016 update’, *Nucleic acids research*, 44(W1), pp. W3–W10. doi: 10.1093/nar/gkw343.

Ahmed, W. *et al.* (2017) ‘Transcription facilitated genome-wide recruitment of topoisomerase I and DNA gyrase’, *PLOS Genetics*, 13(5), p. e1006754. doi: 10.1371/journal.pgen.1006754.

Butland, G. *et al.* (2005) ‘Interaction network containing conserved and essential protein complexes in Escherichia coli’, *Nature*, 433(7025), pp. 531–37. doi: doi:10.1038/nature03239.

Feng, S. *et al.* (2020) ‘Involvement of Transcription Elongation Factor GreA in Mycobacterium Viability, Antibiotic Susceptibility, and Intracellular Fitness’, *Frontiers in Microbiology*, 11(March), pp. 1–15. doi: 10.3389/fmicb.2020.00413.

Guo, M. S. *et al.* (2021) ‘High-resolution, genome-wide mapping of positive supercoiling in chromosomes’, *eLife*, 10(e67236). doi: 10.7554/eLife.67236.

Karp, P. D. *et al.* (2018) ‘The BioCyc collection of microbial genomes and metabolic pathways’, *Briefings in Bioinformatics*, 20(4), pp. 1085–1093. doi: 10.1093/bib/bbx085.

Landick, R. *et al.* (2014) ‘Genome-wide mapping of the distribution of CarD, RNAP σA, and RNAP β on the Mycobacterium smegmatis chromosome using chromatin immunoprecipitation sequencing’, *Genomics Data*. Elsevier B.V., 2, pp. 110–113. doi: 10.1016/j.gdata.2014.05.012.

Langmead, B. *et al.* (2009) ‘Ultrafast and memory-efficient alignment of short DNA sequences to the human genome’, *Genome Biology*, 10(3). doi: 10.1186/gb-2009-10-3-r25.

Li, H. *et al.* (2009) ‘The Sequence Alignment/Map format and SAMtools’, *Bioinformatics*, 25(16), pp. 2078–2079. doi: 10.1093/bioinformatics/btp352.

Li, H. and Durbin, R. (2010) ‘Fast and accurate long-read alignment with Burrows-Wheeler transform’, *Bioinformatics*, 26(5), pp. 589–595. doi: 10.1093/bioinformatics/btp698.

Li, X. *et al.* (2017) ‘Transcriptome landscape of Mycobacterium smegmatis’, *Frontiers in Microbiology*, 8(DEC), pp. 1–16. doi: 10.3389/fmicb.2017.02505.

Mooney, R. A. *et al.* (2009) ‘Regulator trafficking on bacterial transcription units in vivo’, *Molecular Cell*, 33(1), pp. 97–108. doi: 10.1016/j.molcel.2008.12.021.

Ramírez, F. *et al.* (2014) ‘DeepTools: A flexible platform for exploring deep-sequencing data’, *Nucleic Acids Research*, 42(W1), pp. 187–191. doi: 10.1093/nar/gku365.

Rani, P. and Nagaraja, V. (2018) ‘Genome-wide mapping of Topoisomerase I activity sites reveal its role in chromosome segregation’. Oxford University Press, pp. 1–12. doi: 10.1093/nar/gky1271.

Thorvaldsdóttir, H., Robinson, J. T. and Mesirov, J. P. (2013) ‘Integrative Genomics Viewer (IGV): High-performance genomics data visualization and exploration’, *Briefings in Bioinformatics*, 14(2), pp. 178–192. doi: 10.1093/bib/bbs017.

Uplekar, S. *et al.* (2013) ‘High-resolution transcriptome and genome-wide dynamics of RNA polymerase and NusA in Mycobacterium tuberculosis’, *Nucleic Acids Research*, 41(2), pp. 961–977. doi: 10.1093/nar/gks1260.

Wang, L., Wang, S. and Li, W. (2012) ‘RSeQC: Quality control of RNA-seq experiments’, *Bioinformatics*, 28(16), pp. 2184–2185. doi: 10.1093/bioinformatics/bts356.

Zhang, Y. *et al.* (2008) ‘Model-based Analysis of ChIP-Seq (MACS)’, *Genome Biology*, 9(9). doi: 10.1186/gb-2008-9-9-r137.
